# Mammalian adipogenesis regulators (Aregs) exhibit robust non- and anti-adipogenic properties that arise with age and involve retinoic acid signalling

**DOI:** 10.1101/2021.02.24.432431

**Authors:** Magda Zachara, Pernille Y. Rainer, Julie M. Russeil, Horia Hashimi, Daniel Alpern, Radiana Ferrero, Maria Litovchenko, Bart Deplancke

## Abstract

Adipose stem and precursor cells (ASPCs) give rise to adipocytes and determine the composition and plasticity of adipose tissue. Recently, several studies have demonstrated that ASPCs partition into at least three distinct cell subpopulations: *Dpp4*+ stem-like cells, *Aoc*3+ pre-adipocyte-like cells, and the enigmatic CD142+ cells. A great challenge now is to functionally characterize these distinct ASPC populations. Here, we focus on CD142+ ASPCs since discrepant properties have been assigned to this subpopulation, from adipogenic to non- and even anti-adipogenic. To address these inconsistencies, we comprehensively characterized mammalian subcutaneous CD142+ ASPCs across various sampling conditions. Our findings demonstrate that CD142+ ASPCs exhibit high molecular and phenotypic robustness, firmly supporting their non- and anti-adipogenic properties. However, these properties emerge in an age-dependent manner, revealing surprising temporal CD142+ ASPC behavioural alterations. Finally, using multi-omic and functional assays, we show that the inhibitory nature of these adipogenesis-regulatory CD142+ ASPCs (Aregs) is driven by specifically expressed secretory factors that cooperate with the retinoic acid signalling pathway to transform the adipogenic state of CD142− ASPCs into a non-adipogenic, Areg-like one.

## Introduction

Although adipogenesis is one of the best-studied cell differentiation paradigms (Rosen and Spiegelman, 2014), we still have limited knowledge of the *in vivo* origin and composition of adipose stem and precursor cells (ASPCs, (Ferrero, Rainer and Deplancke, 2020)). This is partially due to the highly heterogeneous and unstructured nature of adipose tissue depots, which are present in multiple anatomical locations (including subcutaneous and visceral white adipose tissue) and consist of a mixture of different cell types (Cristancho and Lazar, 2011), whose origin and identity differ between distinct fat depots (Cleal, Aldea and Chau, 2017). Driven by the resolving power of single-cell RNA-sequencing (scRNA-seq), several studies have recently investigated and confirmed mammalian ASPC heterogeneity (Burl *et al.*, 2018; Hepler *et al.*, 2018; Schwalie *et al.*, 2018; Cho, Lee and Doles, 2019; Gu *et al.*, 2019; Merrick *et al.*, 2019; Spallanzani *et al.*, 2019; Sárvári *et al.*, 2021). Our own integrative analysis of publicly available scRNA-seq data has thereby allowed the compiling of an ASPC subpopulation consensus based on the fact that the three main identified subpopulations exhibit a remarkable molecular consistency throughout the analysed datasets (Ferrero, Rainer and Deplancke, 2020). These subpopulations include adipose stem-like cells with high expression of *Cd55* and *Dpp4*, pre-adipocyte-like cells with high *Aoc3* and *Icam1* expression, and a rather enigmatic, third population characterized by high *F3* (coding for CD142) and *Clec11a* expression. A hierarchy of these ASPCs has also been proposed with the highly proliferative DPP4+ stem-like cells giving rise to the two other subpopulations: ICAM1+ and CD142+ ASPCs (Merrick *et al.*, 2019) which themselves may be able to interconvert (Merrick *et al.*, 2019; Sárvári *et al.*, 2021). These distinct ASPC subpopulations have been shown to be established as early as post-natal day 12 (P12) in mouse, based on scRNA-seq data (Merrick *et al.*, 2019). At an even earlier developmental stage (P2), immunofluorescence-based *in situ* analyses revealed the presence of anatomically partitioned DPP4+ adipose stem-like and ICAM1+ pre-adipocyte-like cells (Merrick *et al.*, 2019), however, the presence of CD142+ ASPCs in such young mice has not yet been demonstrated.

A great challenge in the field now is to explore whether these molecularly distinct ASPC subpopulations also have different functional properties. In this study, we decided to focus on CD142+ ASPCs. This is because previous work has defined these cells as being not only non-adipogenic but also anti-adipogenic, which is why they were termed “adipogenesis regulators” (Aregs) (Schwalie *et al.*, 2018). More recent, independent findings supported the notion that adipose tissue may harbour a negatively regulatory cell type (Lee *et al.*, 2019), yet both its identity as well as the underlying molecular mechanisms have so far remained ill-defined (Shamsi, Tseng and Kahn, 2021). The presence of an anti-adipogenic cell population could have tremendous implications with regard to how adipose tissue development and homeostasis is regulated in health and disease. This is why further studies are warranted that explore its existence as well as its molecular and functional properties, especially in light of recent, divergent findings showing that CD142+ ASPCs are in fact adipogenic (Merrick *et al.*, 2019). The reasons for these functional discrepancies between studies have remained unclear but are hypothesized to reflect differences in cell isolation and sorting criteria, antibodies, culturing conditions, sex or age (Merrick *et al.*, 2019; Ferrero, Rainer and Deplancke, 2020; Corvera, 2021).

Given the importance of resolving the molecular and functional heterogeneity of ASPCs, here, we set out to systematically address these inconsistencies. Our findings validate the molecular and phenotypic robustness of murine CD142+ ASPCs, authenticating these cells as non-adipogenic inhibitors of adipogenesis. Interestingly however, we demonstrate that these functional properties are age-dependent. Specifically, we show that the molecular identity of CD142+ cells is already established before post-natal day 16 (P16), while their non-adipogenic properties become apparent only four weeks after birth. Importantly, we also confirm the anti-adipogenic properties of adult CD142+ ASPCs and, using a diverse range of multi-omic- and functional assays, provide insights into the molecular mechanisms that control their activity. Particularly, we show that the inhibitory nature of CD142+ ASPCs appears to be driven by a set of specifically expressed secretory factors, involving Tissue factor (CD142) itself as well as Matrix Gla protein (MGP), whose actions may possibly converge onto the retinoic acid (RA) signalling pathway. These factors/pathways seem to function to render CD142+ ASPCs refractory to adipogenesis, while exerting their anti-adipogenic activity by transforming the adipogenic state of CD142− ASPCs into a non-adipogenic, CD142+-like state.

## Results

### CD142+ ASPCs are defined by a specific transcriptomic signature and a robust non-adipogenic phenotype

In recent years, numerous studies have dissected white adipose tissue (WAT) composition at the single-cell level (subcutaneous WAT: (Burl *et al.*, 2018; Schwalie *et al.*, 2018; Cho, Lee and Doles, 2019; Merrick *et al.*, 2019); visceral WAT: (Hepler *et al.*, 2018; Spallanzani *et al.*, 2019; Sárvári *et al.*, 2021); perivascular WAT: (Gu *et al.*, 2019)). Together, these studies provide an opportunity to acquire high-resolution insights into ASPC heterogeneity through data integration, essentially allowing to validate and possibly expand or revise our initial scRNA-seq-based observations (Schwalie *et al.*, 2018). To do so, we integrated data from our own and relevant publicly available mouse subcutaneous adipose tissue scRNA-seq studies (Burl *et al.*, 2018; Schwalie *et al.*, 2018; Merrick *et al.*, 2019) revealing that ASPCs robustly partition among three principal subpopulations in line with earlier observations (**Fig. 1A, Suppl. Fig. 1**, Ferrero et al., 2020). One of these three main ASPC clusters, characterized by high and specific *F3* (coding for CD142) gene expression, proved to be highly robust and stable across increasing clustering resolution (**Fig. 1B**), indicating that these cells are characterized by a clearly delineated and specific transcriptomic signature.

**Figure 1.**
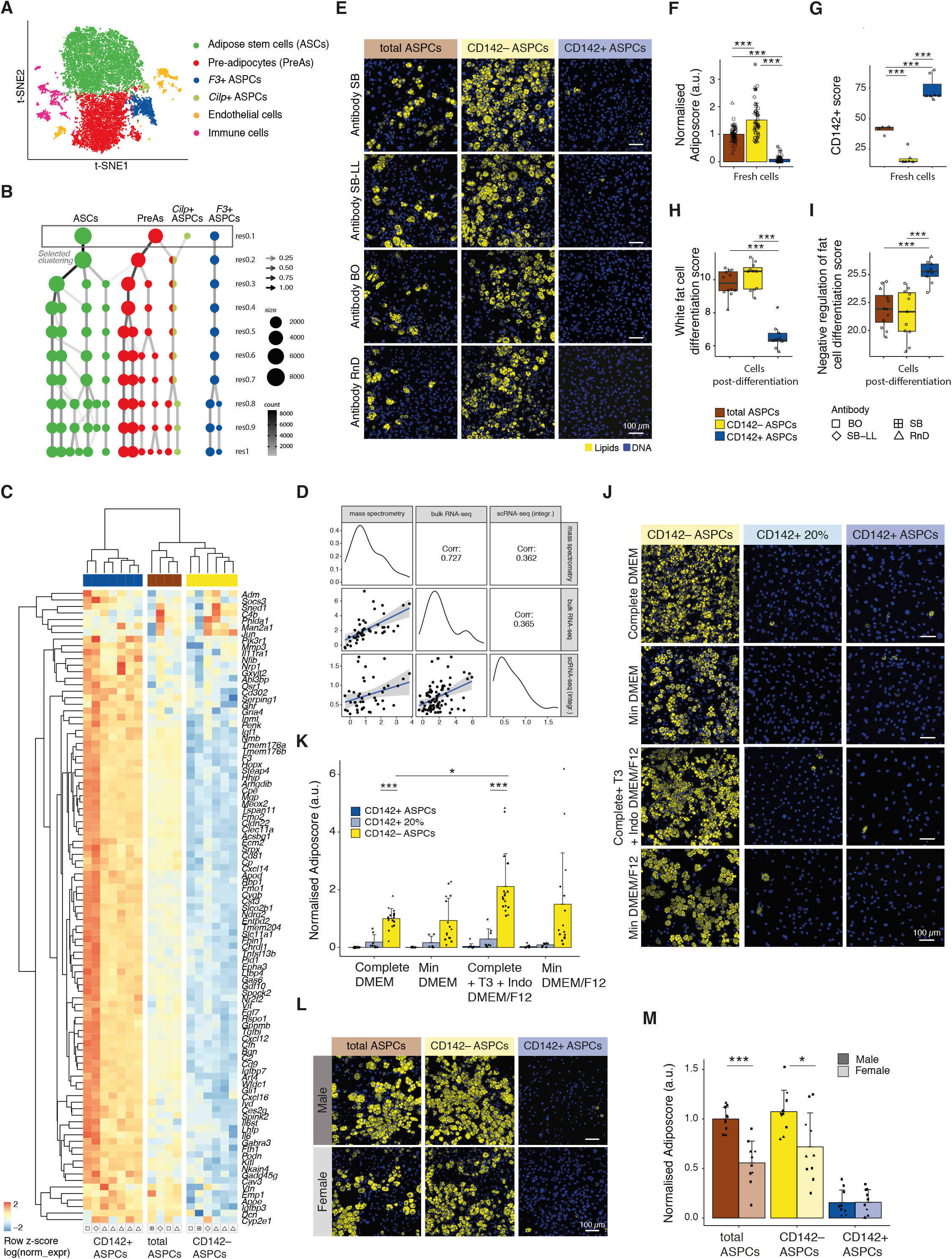
CD142+ ASPCs constitute a robust ASPC subpopulation defined by a stable non-adipogenic phenotype and a highly specific transcriptomic signature. **(A)** t-SNE cell map of integrated scRNA-seq datasets (see **Methods**) visualizing the main identified subpopulations of murine subcutaneous ASPCs: adipose stem cells (ASCs) in green, pre-adipocytes (PreAs) in red, *F3*(CD142)+ ASPCs in blue, as well as *Cilp*+ ASPCs, endothelial and immune cells (see also **Suppl. Fig. 1**); **(B)** Clustering tree of the Seurat-based clustering result of the integrated analysis described in **A**, visualizing the relationships between clustering at different resolutions of the three main ASPC subpopulations as well as *Cilp*+ ASPCs, demonstrating a high stability of the *F3*(CD142)+ ASPC cluster; **(C)** Gene expression heatmap of the top CD142+ markers (**Suppl. Table 1**) across bulk RNA-seq samples of freshly isolated total, CD142− and CD142+ ASPCs; log normalized expression scaled by row; **(D)** Correlation of the logFC of top CD142+ markers (**Suppl. Table 1**) across scRNA-seq, bulk RNA-seq and mass spectrometry data; logFC was defined as the log_2_FC of freshly isolated CD142+ over CD142− ASPCs for bulk RNA-seq and mass spectrometry, and as the average logFC of *F3*(CD142)+ over the remaining cells in scRNA-seq across integrated datasets described in **A** (**Methods**); **(E)** Representative fluorescence microscopy images of total, CD142– and CD142+ ASPCs isolated with the use of the respective anti-CD142 antibodies: SinoBiological (SB), SinoBiological-LL (SB-LL), BiOrbyt (BO) and R&D Systems (RnD) (**Suppl. Fig. 2B**, **Suppl. Fig. 5A**, **Methods**), after *in vitro* adipogenic differentiation; **(F)** Fraction of differentiated cells per ASPC type shown in **E**, as quantified by the “adiposcore”; marker shapes correspond to different anti-CD142 antibodies used for isolation as indicated, n=9-15, 3-4 biological replicates, 3-5 independent wells for each; **(G)** Boxplot showing the distribution of the “CD142+ score” based on the expression of the top CD142+ ASPC markers (**Suppl. Table 1**, **Methods**); (**H**) Boxplot showing the distribution of the “white fat cell differentiation score” based on the expression of the genes linked to the GO term “white fat cell differentiation” (GO:0050872) (**Methods**); **(I)** Boxplot showing the distribution of the “negative regulation of fat cell differentiation score” based on the expression of the genes linked to the GO term “negative regulation of fat cell differentiation” (GO:0045599) (**Methods**); **(J)** Representative fluorescence microscopy images of CD142–, 20% CD142+ and (5-7%) CD142+ ASPCs isolated following the sorting strategy shown in **Suppl. Fig. 7A**, after *in vitro* adipogenic differentiation with the indicated white adipogenic differentiation cocktails (**Methods**); **(K)** Fraction of differentiated cells per ASPC type and differentiation cocktail shown in **J**, as quantified by the “adiposcore”; marker shapes correspond to different biological replicates, n=8-17, 3-5 biological replicates, 2-5 independent wells for each; **(L)** Representative fluorescence microscopy images of male- and female-derived total, CD142– and CD142+ ASPCs, isolated following the sorting strategy shown in **Suppl. Fig. 8A** after *in vitro* adipogenic differentiation; **(M)** Fraction of differentiated cells per ASPC type and per sex shown in **L**, as quantified by the “adiposcore”; bar colour shading corresponds to male- and female-derived cells as indicated; marker shapes correspond to different biological replicates, n=9-10, 4 biological replicates, 2-3 independent wells for each; In all images, nuclei are stained with Hoechst (blue) and lipids are stained with Bodipy (yellow); scale bars, 100 μm; bar colours: total ASPCs — brown, CD142– ASPCs — yellow, CD142+ ASPCs — blue, 20% CD142+ ASPCs — turquoise; *P ≤ 0.05, **P ≤ 0.01, ***P ≤ 0.001, pairwise two-sided *t*-test (**K**, **M**) or one-way ANOVA and Tukey HSD *post hoc* test (**F-I**), for statistical details see **Methods**.

Given the particular interest in the *F3*+ cluster, previously identified to represent cells having inhibitory properties toward adipogenesis (Schwalie *et al.*, 2018), we set out to better understand the specific molecular markers of this population. Using the integrative analysis, we identified a set of robust markers differentially expressed in each individual scRNA-seq dataset that are specific to the *F3*+ cluster. Combined with the top 20 markers that were detected by bulk RNA-seq as being differentially expressed in CD142+ *versus* CD142− freshly isolated ASPCs in our previous study (Schwalie *et al.*, 2018), this resulted in a list of 100 markers that are representative of CD142+ ASPCs (top CD142+ markers, **Suppl. Table 1**, **Methods**). To assess the relevance of this list of gene expression markers, we carried out an in-depth quantitative transcriptomic (BRB-seq, Alpern *et al.*, 2019) and proteomic characterization of FACS-sorted CD142+ *versus* CD142− ASPCs (defined as: SVF Lin− (CD31− CD45− TER119−) SCA-1+ CD142+ and SVF Lin− SCA-1+ CD142− respectively, **Suppl. Fig. 2**, **Methods**). The identified top CD142+ markers revealed to be specific to CD142+ cells both at the transcriptomic and proteomic level (**Fig. 1C-D, Suppl. Fig. 3A-B, Suppl. Table 1-2**). In addition, we found that overall gene and protein expression levels correlated well between all scRNA-seq, bulk RNA-seq, and proteomic datasets (**Fig. 1D**, **Methods**), indicating that the observed transcriptomic signature of CD142+ ASPCs is a reasonable proxy of their protein/functional characteristics. To assess to which extent this signature was affected by culturing or differentiation conditions, we performed bulk RNA-seq of CD142− and CD142+ ASPCs post-expansion and post-differentiation (i.e. after exposure to adipogenic medium) as well as a proteomic analysis of expanded, respective populations (**Suppl. Table 2**). Interestingly, we observed that, under these conditions, the expression of many top CD142+ markers including *Gdf10*, *Cpe*, *Rbp1*, *F3*, *Bgn*, *Clec11a, Mgp* and *Aldh1a2* is maintained both at the transcriptomic and/or proteomic level in CD142+ ASPCs compared to their CD142− counterparts (**Suppl. Fig. 3C-D, Suppl. Fig. 4**).

To examine the functional properties of these cells, we set out to test the phenotype of CD142+ ASPCs across a wide range of experimental conditions and functional assays, aiming to possibly reconcile discrepant findings of CD142+ ASPC behaviour. First, we recapitulated our earlier findings (Schwalie *et al.*, 2018), showing that the top 5-7% most positive CD142+ ASPCs, isolated using the previously employed anti-CD142 antibody, have very low to no adipogenic capacity as compared to CD142− ASPCs when stimulated with a standard white adipogenic differentiation cocktail (**Fig. 1E-F**, **Methods**). However, since the nature of this antibody may be one of the possible reasons underlying discrepant CD142+ cell behaviour read-outs, we tested three more antibodies. While the flow cytometry profiles of the four assessed antibodies differ to a certain extent, the isolated cellular fractions yielded consistent non-adipogenic phenotypic results (**Fig.1E-F**, **Suppl. Fig. 5A**, **Methods**). Indeed, when freshly isolated with different antibodies, the respective CD142+ ASPC samples revealed a consistent transcriptional signature exhibiting a significantly higher expression score based on the top CD142+ markers (here named the “CD142+ score”, **Suppl. Table 1**, **Methods**) compared to the other tested cellular fractions (total and CD142− ASPCs) (**Fig. 1C** and **G**). Finally, the observed non-adipogenic properties of post-differentiation CD142+ cells (i.e. post-exposure to a standard, adipogenic cocktail) were consistent with the expression profiles of adipogenesis-relevant genes. Specifically, genes involved in “white fat cell differentiation” (GO:0050872) or “negative regulation of fat cell differentiation” (GO:0045599) were significantly lower or higher expressed in CD142+ ASPCs compared to total or CD142− ASPCs, respectively (**Fig. 1H-I**). Moreover, fat and lipid-related terms that were detected as significant by gene set enrichment analysis (GSEA) were negatively enriched in CD142+ *versus* CD142− ASPCs (**Suppl. Fig. 5B**).

Finally, since CD142 surface expression shows a continuum across ASPCs (**Suppl. Fig. 5A**), it is possible that the stringency of the cell isolation procedure (so far typically 5-7%) has an impact on downstream cell behaviour. To investigate this, we isolated CD142+ ASPCs using a less stringent gating (~20% **Suppl. Fig. 7A**), yet we did not observe a notable difference in overall differentiation potential compared to the more stringently isolated cells (**Fig. 1J-K, Suppl. Fig. 7B-C**).

Next, we examined the influence of the differentiation medium as it is widely established that adipogenic potential varies as a function of the utilized differentiation cocktail. To do so, we used the standard “complete DMEM” differentiation medium (with insulin, IBMX and dexamethasone) as well as three additional ones (1, “Min DMEM”, with insulin only; 2, “Complete + T3 + Indo DMEM/F12”, with insulin, IBMX, dexamethasone, T3 and indomethacin; 3, “Min DMEM/F12”, with insulin only, **Methods**, Schwalie *et al.*, 2018; Merrick *et al.*, 2019). However, we did not observe notable CD142+ ASPC differentiation differences across these distinct culturing conditions (**Fig. 1J-K**).

Another source for discrepant cell behaviour could be the sex of the animals. To test this, we isolated CD142+ and CD142− ASPCs from both male and female mice, revealing that, upon exposure to an adipogenic cocktail, the adipogenic propensity of total and CD142− ASPCs was significantly higher (adjusted p-value (p-adj) < 0.001 and < 0.05, respectively) in males compared to females (**Fig. 1L-M**, **Suppl. Fig. 8)**. However, both male and female CD142+ cells were completely refractory to adipogenic differentiation (**Fig. 1L-M**, **Suppl. Fig. 8**). This is in line with transcriptomic results, as our scRNA-seq analysis of male and female cells (**Suppl. Fig. 9A**) (Schwalie *et al.*, 2018) revealed the existence of a clearly delineated *F3*(CD142)+ cluster in both sexes with a highly consistent overlap of specific markers (**Suppl. Fig. 9B-C**). In addition, the integration of the different publicly available scRNA-seq datasets included cells from mice of different sexes and clearly showed the existence of a robust *F3*+ cluster (Burl *et al.*, 2018; Schwalie *et al.*, 2018; Merrick *et al.*, 2019) (**Fig. 1B, Suppl. Fig. 1**).

Together, these in-depth computational and experimental analyses validate the previously observed molecular identity of CD142+ ASPCs and demonstrate the robustness of their non-adipogenic phenotype across a wide range of conditions and experiments.

### Age-dependent molecular and phenotypic emergence of *bona fide* CD142+ ASPCs

The findings reported above indicate that CD142+ ASPCs constitute a distinct cell population with a well-defined molecular identity and a clear non-adipogenic character. However, all of these analyses were performed on cells derived from adult mice, prompting the question whether the observed CD142+ cell properties could perhaps be age-dependent. Analysis of publicly available scRNA-seq data of post-natal day 12 (P12) mice revealed a CD142+ cluster that shares many of the adult top CD142+ markers (Merrick *et al.*, 2019, **Suppl. Fig. 10A-B**) and overlaps with adult CD142+ ASPCs upon data integration (**Suppl. Fig. 10C-D**). Nevertheless, we uncovered that the identity of these P12 *F3*(CD142)+ cells, as defined by their “CD142+ score” (**Suppl. Table 1**), is significantly less pronounced compared to their adult counterpart (p-value < 0.001, **Fig. 2A, Suppl. Fig. 10E**). To investigate whether this more subtle molecular identity would also manifest itself at the phenotypic level, we assessed the proportion and adipogenic propensity of distinct ASPC subpopulations at distinct developmental time points including newborn (P0), 12-17-day-old (P12-17) mice as well as juvenile (4-week-old (4wo)) and adult (7wo and 11wo) animals with these two groups being separated by weaning at post-natal day 21 (P21). Given the overlap between P12 and adult CD142+ ASPCs in the scRNA-seq analysis and the specificity of *F3*(CD142) in both age groups (**Suppl. Fig. 10B**) (Merrick *et al.*, 2019), we used the CD142 marker to enrich for our cellular fractions of interest across ages. We observed significant differences in the proportions of cellular fractions assessed by flow cytometry, with the Lineage negative (SVF Lin–) portion being significantly higher (p-adj < 0.0001) and the ASPC portion (SVF Lin– SCA-1+) significantly lower (p-adj < 0.05) in pre-weaning (P0 and P12-17) compared to post-weaning mice (4wo, 7wo and 11wo) (**Fig. 2B-C**, **Suppl. Fig. 11A**). In addition, ASPCs from pre-weaning mice exhibited significantly lower (p-adj < 0.01) CD142 cell surface expression compared to post-weaning animals (**Fig. 2B-C**). Indeed, we found that at the gating stringency corresponding to 5% of CD142+ cells in adult (11wo) mice, only 2.5% of newborn (P0) CD142+ ASPCs were captured (**Fig. 2B-C**), consistent with the observed gradual decrease of the CD142− ASPC fraction as mice mature (**Suppl. Fig. 11A-B**).

**Figure 2.**
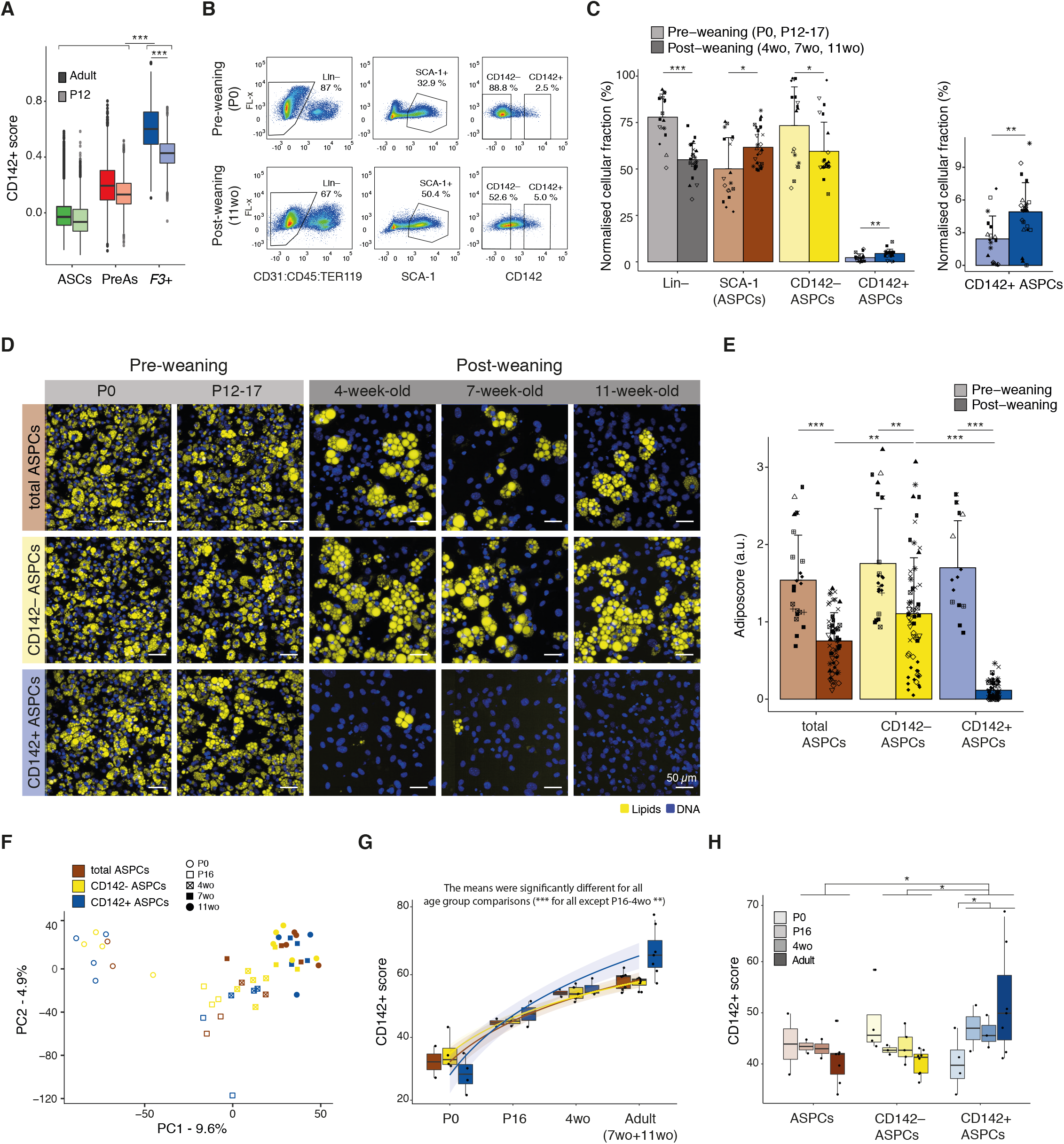
Age-dependent molecular and phenotypic emergence of *bona fide* CD142+ ASPCs. **(A)** Boxplot showing the distribution of the “CD142+ score” (**Suppl. Table 1**) in adult (darker colours) and P12 (lighter colours) ASPCs across the three main ASPC subpopulations (adipose stem cells (ASCs) in green, pre-adipocytes (PreAs) in red and *F3*(CD142)+ ASPCs in blue); **(B)** FACS-based gating strategy of pre-weaning (new-borns (P0))- and post-weaning (11-week-old (wo))-derived Lin– (defined as CD31– CD45– TER119–), SCA-1+ (total ASPC), CD142– and CD142+ ASPC cellular fractions within the subcutaneous adipose SVF; at least 8 biological replicates were performed, shown here is one representative biological replicate; **(C)** Bar plots showing the normalised parental percentage of pre- and post-weaning indicated cellular fractions; the graph on the right represents the fractions of CD142+ ASPCs plotted separately; marker shapes correspond to different biological replicates, n=8-10; **(D)** Representative fluorescence microscopy images of P0-, P12-17-, 4wo-, 7wo- and 11wo-derived total, CD142– and CD142+ APSCs after *in vitro* adipogenic differentiation; **(E)** Fraction of differentiated cells per ASPC type shown in **D**, as quantified by the “adiposcore”; bar colour shading corresponds to pre- and post-weaning-derived cells as indicated; marker shapes correspond to different biological replicates, n=14-65, 12-13 biological replicates, 2-6 independent wells for each; **(F)** PCA based on the bulk RNA-seq data of freshly isolated P0-, P16-, 4wo-, 7wo- and 11wo-mice-derived total, CD142– and CD142+ ASPCs; **(G)** Boxplot showing the distribution of the “CD142+ score” (**Suppl. Table 1**) across ages and tested cellular fractions (freshly isolated total, CD142– and CD142+ ASPCs) showing the age-dependent emergence of the CD142+ signature; see colour legend in **F**; **(H)** Boxplot showing the distribution of the “CD142+ score” (**Suppl. Table 1**) across ages and tested cellular fractions (freshly isolated total, CD142– and CD142+ ASPCs) using the log normalized expression corrected for age-driven source of variation; see colour legend in **F;** In all images, nuclei are stained with Hoechst (blue) and lipids are stained with Bodipy (yellow); scale bars, 50 μm; bar colours: total ASPCs — brown, CD142– ASPCs — yellow, CD142+ ASPCs — blue; pre-weaning data are represented in lig ter s ades; *P ≤ 0.05, **P ≤ 0.01, ***P ≤ 0.001, pairwise two-sided *t*-test (**A**, **C**, **E**, **H**) or one-way ANOVA and Tukey HSD *post hoc* test (**G**, null hypothesis: no difference in means across age), for statistical details see **Methods**.

Next, we assessed the adipogenic capacity of the different ASPC fractions as a function of age. Using the standard differentiation cocktail (**Methods**), we unexpectedly observed that all cellular fractions (total, CD142− and CD142+ ASPCs) derived from pre-weaning mice exhibited a remarkable adipogenic propensity with virtually all cells displaying lipid accumulation. However, based on visual inspection, the overall size of the pre-weaning cells and the size of the accumulated lipid droplets tended to be smaller compared to post-weaning cells (**Fig. 2D**). Perhaps most interestingly, pre-weaning-derived CD142+ ASPCs gave rise to *in vitro* adipocytes, which is in stark contrast to the marked non-adipogenic properties of their post-weaning counterparts (**Fig. 2D-E**, **Suppl. Fig. 11C-E**). Indeed, we observed that, despite non-negligible variability between independent replicates, the adipogenic propensity of total and CD142− ASPCs gradually decreases with age, whereas CD142+ ASPCs exhibit a very sharp drop (p-adj < 0.0001) in their ability to give rise to *in vitro* adipocytes between the 16^th^ and 28^th^ (4wo) day of post-natal development (**Fig. 2D-E**, **Suppl. Fig. 11C-E**).

To further explore this age-dependent functional change of CD142+ ASPCs, we performed bulk RNA-seq on freshly isolated ASPC cellular fractions at different time points after birth (P0, P16, 4wo, 7wo and 11wo, **Suppl. Table 2**). Analysis of the resulting data revealed that age is the variable that explains the largest variation across samples as they are ordered by this feature along the first principal component (PC1) (**Fig. 2F**). GSEA revealed that terms such as “cell fate commitment” (GO:0045165), “mesenchymal cell differentiation/proliferation and development” (GO:0048762, GO:0010463, GO:0014031), “stem cell differentiation” (GO:0048863), “stem cell population maintenance” (GO:0019827) and cell cycle-related terms are negatively enriched in the genes driving PC1 (**Suppl. Fig. 12A**). This suggests that genes linked to these terms are up-regulated in freshly isolated young ASPCs, likely reflecting their more naïve “stem” condition. Indeed, all ASPC fractions that were freshly isolated from the newborn bourgeoning fat pads expressed substantially higher levels of *Ccnd1* (coding for CyclinD1) and *Dlk1*/*Pref1*, and lower levels of markers such as *Pdgfrb*, *Cd34* and *Ly6a* (coding for SCA-1) compared to ASPCs from older animals (**Suppl. Fig. 12B-F**). Such a signature was proposed to select for a primitive and naïve precursor population (Wang *et al.*, 2003; Atanassova, Rancic and Georgieva, 2012; Hepler and Gupta, 2017). Interestingly, it has also been shown that foetal and early post-natal ASPCs from murine subcutaneous depots surprisingly express perilipin (*Plin1*) and adiponectin (*Adipoq*) and exhibit highly proliferative and differentiation properties (Hong *et al.*, 2015; Hepler and Gupta, 2017), consistent with our observations for newborn-derived ASPCs (**Suppl. Fig. 12G-H**, **Supp. Fig. 11C-E**). In contrast, lipid/fat-related terms were enriched in freshly isolated adult ASPCs, implying a molecular state which is less naïve and more committed towards adipogenesis (**Suppl. Fig. 12A**). We further found that the “retinol/retinoid metabolic process” (GO:0042572/GO:0001523) gene expression signature is much more prominent in adult-derived ASPCs compared to young ASPCs, which correlates with the decreased adipogenic capacity of adult *versus* young ASPCs. (**Suppl. Fig. 13A-B**). In addition to this general increase with age, this same RA-related expression signature was even more pronounced in 4wo and adult CD142+ compared to CD142− ASPCs (**Suppl. Fig. 13C**). Interestingly, the “CD142+ score” (**Suppl. Table 1**) increased in all the cellular fractions of ASPCs (CD142−, CD142+ and total ASPCs) in an age-dependent manner (**Fig. 2G, Suppl. Fig. 14A**). We thereby observed that this set of top markers becomes significantly enriched in CD142+ ASPCs (compared to CD142− ASPCs) at P16 and is particularly prominent in adult cells (**Suppl. Fig. 14B-E**). Moreover, when we removed the age-driven source of variation from the analysed samples, CD142+ ASPCs from all developmental time points, except P0, were marked by a relative and substantial increase of the “CD142+ score” compared to the other assessed fractions (all ages, total and CD142− ASPCs), while being once again more pronounced in adults, reflecting the results from our scRNA-seq analysis of P12- and adult-derived ASPCs (**Fig. 2A**) (Merrick *et al.*, 2019). This increase was accompanied by a gradual decrease of the “CD142+ score” in all CD142− ASPCs with age (**Fig. 2H**).

Together, these findings suggest that ASPCs and their considered subpopulations are molecularly and phenotypically naïve at birth, after which they gradually acquire their respective properties throughout the early post-natal developmental stages. Furthermore, the definite “CD142+-specific expression signature” appears to emerge in CD142+ ASPCs before P16, followed by the manifestation of their completely non-adipogenic phenotype by post-natal week 4.

### CD142+ ASPC-dependent adipogenic inhibition and mediating factors

The findings above validate the previously described molecular identity of CD142+ ASPCs, provide additional evidence as to the robustness of their non-adipogenic character, and add the interesting dimension of this phenotypic property emerging with age. Importantly, our results also corroborate the notion that adult CD142+ ASPCs are not only non-adipogenic, but also anti-adipogenic. This is because adult CD142− ASPCs reproducibly exhibit a greater adipogenic propensity than total ASPCs (**Fig. 1E-F**, **Fig. 2D-E**), suggesting that the presence of CD142+ ASPCs dampens the adipogenic capacity of their CD142− counterparts (as a progenitor population devoid of CD142+ ASPCs and featuring a high adipogenic potential). To better understand underlying mechanisms governing the inhibitory function of CD142+ ASPCs, we first re-explored the anti-adipogenic nature of these cells using a transwell set-up allowing specific ASPC subpopulations to be co-cultured but without cell-to-cell contact (**Methods**). These experiments revealed that CD142− ASPCs co-cultured with CD142+ ASPCs show a significantly decreased capacity to generate adipocytes (p-value < 0.01) compared to CD142− ASPCs cultured on their own (**Fig. 3A-B**). To molecularly support these results, we performed bulk RNA-seq of differentiated CD142− ASPCs and differentiated CD142− ASPCs co-cultured with CD142+ ASPCs (**Suppl. Table 2**). Consistent with our phenotypic observations, CD142− ASPCs cultured alone showed a significantly higher expression of genes related to “white fat cell differentiation” (GO:0050872) (**Fig. 3C**), while key genes linked to the “negative regulation of fat cell differentiation” (GO:0045599) were up-regulated in CD142− ASPCs co-cultured with CD142+ cells (**Fig. 3D**). These results recapitulate CD142+ ASPCs’ previously reported capacity to inhibit adipogenesis through paracrine signalling (Schwalie *et al.*, 2018). Together with a recent, independent report showing a comparable capacity of CD142+ stromal cells in muscle (Camps *et al.*, 2020), we therefore decided to reintroduce the original “Adipogenesis regulators” (Aregs) nomenclature to conceptually define CD142+ ASPCs.

**Figure 3.**
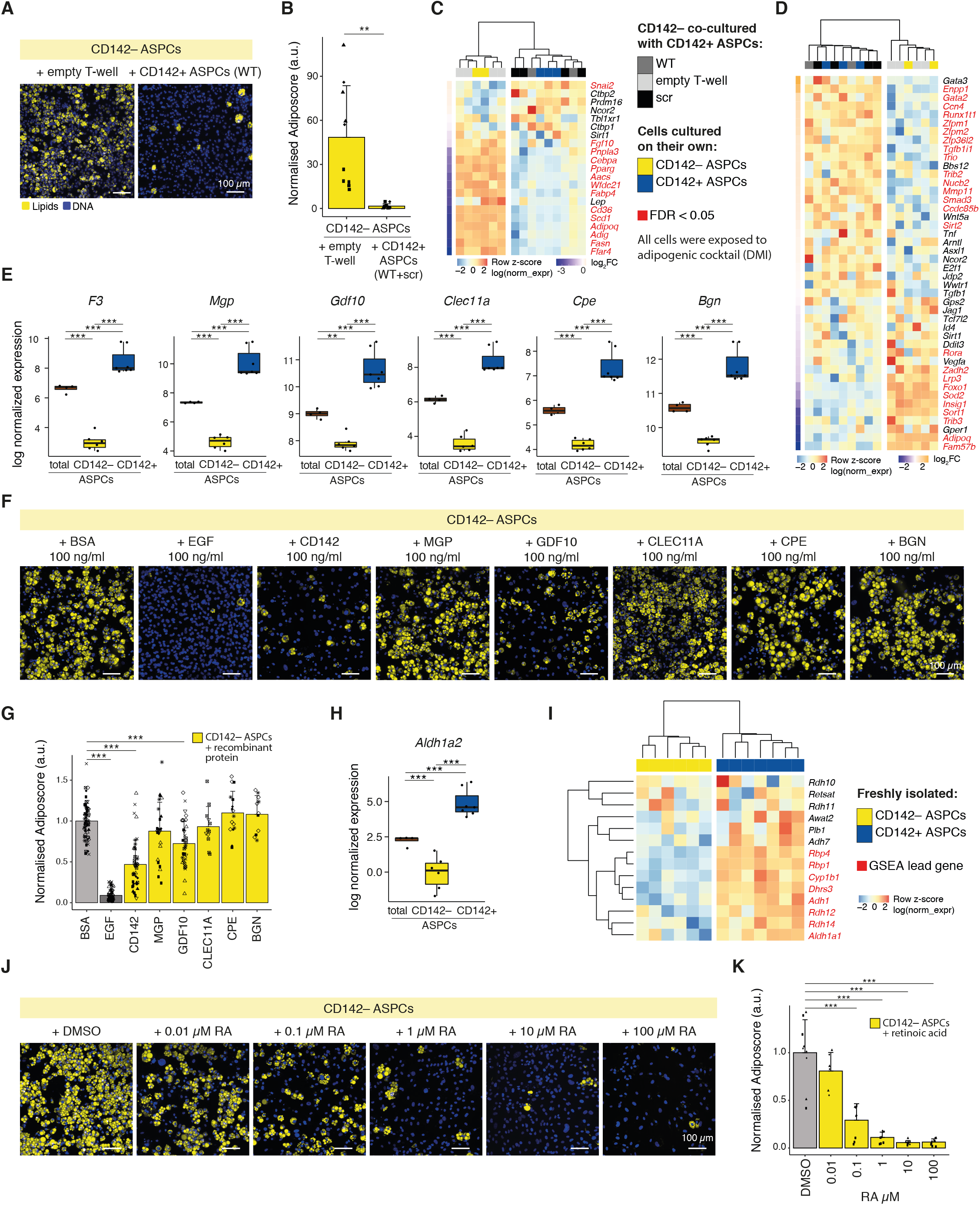
CD142+ ASPC (Areg)-specific candidates and their involvement in adipogenic inhibition. **(A)** Representative fluorescence microscopy images of CD142– ASPCs co-cultured with an empty transwell (T-well) or with wild-type (WT) CD142+ ASPCs or CD142+ ASPCs carrying control siRNA (scr) after *in vitro* adipogenic differentiation; **(B)** Fraction of differentiated CD142– ASPCs indicated in **A**, as quantified by the “adiposcore”; marker shapes correspond to different biological replicates, n=10, 4 biological replicates, 2-3 independent wells for each; **(C)** Expression heatmap listing genes linked to “white fat cell differentiation” (GO:0050872) across bulk RNA-seq samples of CD142+ and CD142– ASPCs after adipogenic differentiation (i.e. after exposure to an adipogenic cocktail, **Methods**); the genes are ordered from top to bottom by the log_2_FC of CD142+ over CD142– ASPCs after adipogenic differentiation; significantly differentially expressed genes (FDR < 0.05) are coloured in red; log normalized expression scaled by row; **(D)** Expression heatmap listing genes linked to “negative regulation of fat cell differentiation” (GO:0045599) across bulk RNA-seq samples of CD142+ and CD142– ASPCs after adipogenic differentiation; the genes are ordered from top to bottom by the log_2_FC of CD142+ over CD142– ASPCs after adipogenic differentiation; significantly differentially expressed genes (FDR < 0.05) are coloured in red; log normalized expression scaled by row; **(E)** Bulk RNA-seq-derived expression plots of CD142+ ASPC (Areg) markers coding for secreted proteins that were selected for downstream validation: *F3* (coding for CD142), *Mgp* (coding for Matrix Gla protein, MGP), *Gdf10* (GDF10), *Clec11a* (CLEC11A), *Cpe* (Carboxypeptidase E, CPE) and *Bgn* (Biglycan, BGN); **(F)** Representative fluorescence microscopy images of adult-derived CD142– ASPCs after *in vitro* adipogenic differentiation; the induction cocktail was supplemented with recombinant proteins corresponding to the selected Areg-specific candidates: CD142, MGP, GDF10, CLEC11A, CPE and BGN at 100 ng/ml (**Methods**); **(G)** Fraction of differentiated adult-derived CD142– ASPCs, as quantified by the “adiposcore”, treated with the indicated recombinant proteins shown in **F**; marker shapes correspond to different biological replicates, n=11-59, 2-10 biological replicates, 3-9 independent wells for each; **(H)** Bulk RNA-seq-derived expression plot of *Aldh1a2* (coding for Retinal dehydrogenase, RALDH2); **(I)** Expression heatmap listing genes linked to “retinol metabolic process” (GO:0042572) across bulk RNA-seq samples of freshly isolated CD142– and CD142+ ASPCs; genes identified as lead by GSEA (**Methods**) are coloured in red; log normalized expression scaled by row; **(J)** Representative fluorescence microscopy images of adult-derived CD142– ASPCs after *in vitro* adipogenic differentiation with the differentiation cocktail supplemented with DMSO (RA carrier) and RA at the indicated concentrations; **(K)** Fraction of differentiated CD142– ASPCs, as quantified by the “adiposcore”, treated with the indicated recombinant proteins shown in **J**; marker shapes correspond to different biological replicates, n=9, 3 biological replicates, 3 independent wells for each; In all images, nuclei are stained with Hoechst (blue) and lipids are stained with Bodipy (yellow); scale bars, 100 μm, bar colours: total ASPCs — brown, CD142– ASPCs — yellow, CD142+ ASPCs (Aregs) — blue, recombinant BSA or DMSO treatment (negative controls) - lig t grey, recombinant EGF treatment (positive control) - dark grey; *P ≤ 0.05, **P ≤ 0.01, ***P ≤ 0.001, pairwise two-sided *t*-test (**B**, **G**, **K**) or one-way ANOVA and Tukey HSD *post hoc* test (**E**, **H**), for statistical details see **Methods**.

Given that Aregs can exert their activity *via* paracrine signalling, we explicitly mined the transcriptome and proteome data to identify Areg-specific *secreted* factors. This resulted in a stringent set of highly Areg-specific candidates including *F3* (coding CD142) itself, *Mgp, Gdf10, Clec11a, Cpe* and *Bgn* (**Fig. 3E, Suppl. Table 1**, **2** and **Methods**). To assess the inhibitory potential of these factors, we treated CD142− ASPCs with various concentrations of recombinant candidate proteins, aiming to mimic the physiological presence of Aregs. Similar to the above-described experiments, the extent of adipogenesis was inherently variable across distinct assays and cell populations. Nevertheless, both recombinant CD142 (p-adj < 0.001) and GDF10 (p-adj < 0.001) significantly inhibited adult-derived CD142− ASPC adipogenesis at a concentration of 100 ng/ml, while recombinant MGP inhibited adipogenesis at 1 μg/ml (p-adj < 0.001) (**Fig. 3F-G**, **Suppl. Fig. 15** and **16**, **Methods**). Interestingly, P0-derived CD142− ASPCs appeared to a large extent refractory to such an inhibition, illustrated by a much less striking decrease in the extent of adipogenesis upon treatment with recombinant EGF, a well-established adipogenesis inhibitor (Harrington, Pond-Tor and Boney, 2007) that was used as a positive control (**Suppl. Fig. 17**, **Methods**). Nevertheless, we still observed a significant decrease in the extent of adipogenesis when newborn CD142− ASPCs were exposed to recombinant CD142, GDF10, MGP and BGN (p-adj < 0.01, < 0.01, < 0.001 and < 0.05 respectively), with MGP now exerting the most pronounced inhibitory effect on differentiating newborn-derived CD142– ASPCs (**Suppl. Fig. 17**).

Amongst other Areg candidates, we also identified a few retinoic acid (RA)-related genes including *Aldh1a2*, *Epha3*, *Osr1* and *Rbp1* (**Fig. 3H**, **Suppl. Fig. 18**, **Suppl. Table 1**, **Methods**). Their specificity to CD142+ ASPCs raises the possibility that this pathway could also be implicated in the anti-adipogenic character of Aregs. This is further supported by the significant enrichment of the GO term “retinol metabolic process” (GO:0042572) (with retinol being a precursor of RA) in transcriptomic and proteomic data from freshly isolated CD142+ compared to CD142− ASPCs (**Fig. 3I**, **Suppl. Fig. 19A-B**). We further uncovered that transcriptomic data from cultured CD142+ ASPCs treated with standard white adipogenic cocktail is enriched for the “cellular response to RA” (GO:0071300) and “RA receptor signalling pathway” (GO:0048384) terms compared to CD142− ASPCs post-differentiation (**Suppl. Fig. 19C-D**, **Suppl. Fig. 20**). Given that RA has been previously shown to inhibit adipogenesis of 3T3 cells (Murray and Russell, 1980), these findings suggest that Aregs might auto-suppress their own adipogenic differentiation capacity by actively responding to RA. To test the functional implication of RA in Aregs’ inhibitory properties, we treated CD142− ASPCs with RA (**Methods**) and observed a significant decrease (p-adj < 0.001) in the extent of adipogenesis (from 0.1 μM RA, **Fig. 3J-K, Suppl. Fig. 21**). Together, these findings suggest that the inhibitory character of Aregs might be mediated *via* RA signalling in concert with other factors such as CD142, GDF10 and MGP.

### Molecular (auto-)regulation of Areg-mediated inhibitory activity

To validate the involvement of factors shown implicated in the inhibitory nature of CD142+ ASPCs on adipogenesis, i.e. *F3* (CD142), *Mgp*, *Gdf10* and *Aldh1a2*, we first knocked the respective genes down in adult CD142+ and total ASPCs in an siRNA-dependent manner and examined its impact on adipogenesis. We observed that for each of these four genes, their respective knockdown (KD) lead to a variable but consistent increase of adipogenic propensity both in CD142+ and total ASPCs, with more pronounced effects in the total ASPC population (**Suppl. Fig. 22**, **Methods**). Indeed, we found that siRNA-mediated changes in lipid accumulation in CD142+ ASPCs were small, suggesting an inherent inability of these cells to give rise to adipocytes, at least in the imposed culturing conditions. Given the Areg-specific (among ASPCs) expression of *F3, Mgp*, *Gdf10* and *Aldh1a2* genes, we interpret the increase of total ASPC adipogenesis upon knockdown of these respective genes as a consequence of specifically inactivating Areg function. To support this interpretation in a more rigorous way, we performed transwell assays, allowing CD142− ASPCs to be exposed to the secretome of CD142+ ASPCs in which the respective candidate factors were knocked down. We observed that inactivating all four genes caused an increased adipogenesis of co-cultured CD142− ASPCs compared to control (scr siRNA), suggesting that they are indeed involved in the inhibitory activity of Aregs. However, while we observed that *F3* and *Mgp* inactivation caused a dramatic increase in overall differentiation of CD142− ASPCs (p-adj < 0.01 and < 0.01 respectively, **Fig. 4A-B**, **Suppl. Fig. 22H-I**), *Gdf10* and *Aldh1a2* KD had a lower effect (p-adj = 0.052 for *Gdf10*, p-adj = 0.029 for *Aldh1a2*).

**Figure 4.**
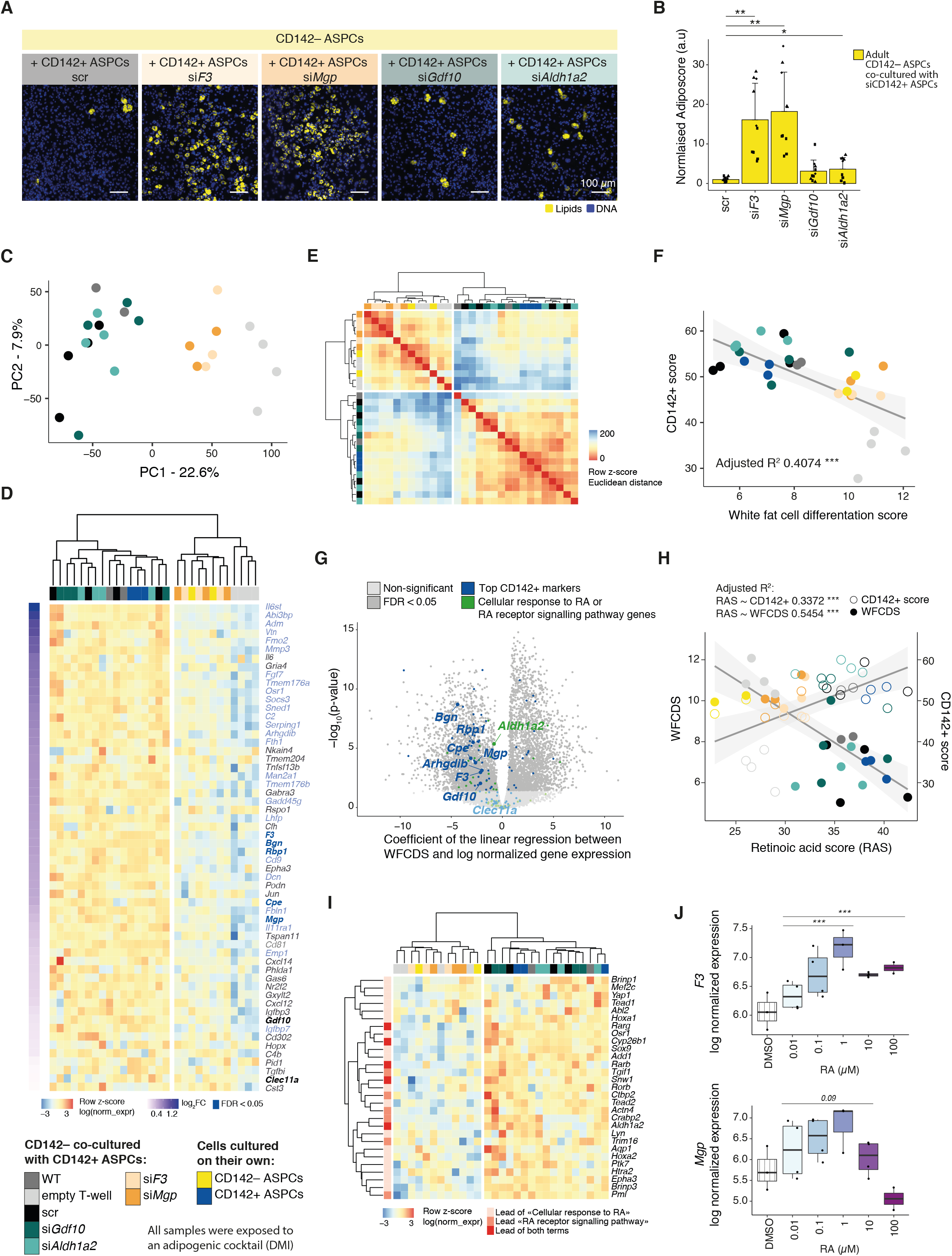
Molecular (auto-)regulation of CD142+ ASPC (Areg)’s inhibitory activity *via* RA signalling. **(A)** Representative fluorescence microscopy images of CD142– ASPCs, co-cultured with CD142+ ASPCs (Aregs) carrying control (scr) or siRNA-mediated knockdowns of selected CD142+ ASPC (Areg)-specific candidate genes: *F3*, *Mgp*, *Gdf10* and *Aldh1a2* after adipogenic differentiation (i.e. after exposure to an adipogenic cocktail, **Methods**); **(B)** Fraction of differentiated CD142– ASPCs, as quantified by the “adiposcore”, co-cultured with CD142+ ASPCs (Aregs) carrying knockdowns indicated in **A**; marker shapes correspond to different biological replicates, n=10, 4 biological replicates, 2-3 independent wells for each; **(C)** PCA based on bulk RNA-seq data of CD142– ASPCs co-cultured with CD142+ ASPCs (Aregs) carrying knock-downs as indicated in **A**; see colour legend at the bottom of the figure; **(D)** Expression heatmap of top CD142+ markers (i.e. Areg markers, **Suppl. Table 1**) with a positive log_2_FC when performing differential expression analysis between CD142– ASPCs that were exposed to “active Aregs” (WT, scr, si*Gdf10*, siAldh1a2) *versus* CD142– ASPCs that were exposed to “dysfunctional Aregs” (si*F3*, si*Mgp*) across the same bulk RNA-seq samples; genes are ordered from the highest (top) to the lowest (bottom) log_2_FC; significantly differentially expressed genes (FDR < 0.05) are coloured in blue; Areg candidates are highlighted in bold; log normalized expression scaled by row; see colour legend at the bottom of the figure; **(E)** Heatmap showing the Euclidian distance of the transcriptomic data of CD142– ASPCs co-cultured (*via* transwell) with distinct CD142+ ASPC knockdown types or controls as well as CD142+ or CD142– ASPCs post-differentiation, calculated on the five first principal components of the PCA shown in **Suppl. Fig. 23D**; **(F)** Correlation of “white fat cell differentiation score” *versus* “CD142+ score”; see colour legend at the bottom of the figure; **(G)** Volcano plot showing the coefficient of the linear regression performed between “white fat cell differentiation score” (WFCDS) and log normalized gene expression (WFCDS | log(normalized expression)) (x axis) *versus* the −log_10_(p-value) (y axis); top CD142+ markers (i.e. Areg markers, **Suppl. Table 1**) are highlighted in blue and genes linked to “cellular response to RA” (GO:0071300) and “RA receptor signalling pathway” (GO:0048384) terms in green; **(H)** Correlation plot of “RA score” (RAS) (based on the expression of genes linked to “cellular response to RA” (GO:0071300) and “RA receptor signalling pathway” (GO:0048384)) *versus* “CD142+ score” (**Suppl. Table 1**) or “white fat cell differentiation score” (WFCDS) (GO:0050872); **(I)** Expression heatmap of the lead genes identified by GSEA of the significantly enriched terms “cellular response to RA” (GO:0071300) and “RA receptor signalling pathway” (GO:0048384); GSEA identified these two terms as enriched when performed on the genes driving PC1 shown in **C**; see colour legend at the bottom of the figure; **(J)** Boxplots showing the distribution of the log normalized expression of *F3* (**top**) or *Mgp* (**bottom**) across transcriptomic data of CD142– ASPCs treated with an adipogenic cocktail supplemented with DMSO or different concentrations of RA (0.01 to 100 *μ*M); the indicated significance is based on the result of differential expression analysis between all RA treated samples or samples treated with RA of a concentration between 0.01 and 1 μM *versus* controls (treatment with DMI with or without DMSO); In all images, nuclei are stained with Hoechst (blue) and lipids are stained with Bodipy (yellow); scale bars, 100 μm; *P ≤ 0.05, **P ≤ 0.01, ***P ≤ 0.001, pairwise two-sided *t*-test (**B**), for statistical details see **Methods**.

To further characterise the molecular mechanism(s) underlying the observed Areg-mediated inhibitory signalling, we profiled the transcriptomes of CD142− ASPCs responding to CD142+ ASPCs, whose activity was modulated *via* individual knockdowns (**Suppl. Table 2**). We observed that the respective gene expression profiles reflected the image-based differentiation results to a great extent, with adipogenic propensity differences driving the first principal component (PC1) and correlating with the overall transcriptomics-based “white fat cell differentiation score” (GO:0050872) (**Fig. 4C**, **Suppl. Fig. 23A-B**). Interestingly, distinct CD142− ASPC samples grouped as a function of how effectively they were impacted by CD142+ ASPC signalling, forming two clusters corresponding to “active Areg” signalling (CD142− ASPCs co-cultured with si*Gdf10*, si*Aldh1a2*, as well as WT and scr Aregs) and what we interpret as “dysfunctional Areg” signalling (co-cultured with si*F3*, si*Mgp* Aregs and with an empty transwell, **Fig. 4C-E**). Furthermore, we found that in CD142− ASPCs that responded to “active Aregs”, the expression of most top CD142+ markers (i.e. Areg-specific markers) including *F3*, *Bgn*, *Rbp1*, *Osr1*, *Cpe*, *Mgp* and *Gdf10* is higher compared to the other CD142− ASPC samples, suggesting that the molecular state of CD142− ASPCs that were exposed to “active Aregs” became itself more Areg-like (**Fig. 4D**, **Suppl. Fig. 23C**). To test this hypothesis, we integrated transcriptomic profiles of CD142− and CD142+ ASPCs that were on their own exposed to an adipogenic cocktail for 6-8 days (**Suppl. Table 2**) into the analysis of bulk RNA-seq data derived from CD142− ASPCs that were co-cultured with distinct KD CD142+ ASPCs. Remarkably, post-differentiation CD142+ and CD142− ASPCs fell into the two distinct clusters corresponding to “active” and “dysfunctional Aregs” respectively, indicating that the CD142+ ASPCs were transcriptionally similar to CD142− ASPCs that were exposed to “active Aregs” (**Fig. 4E-D**, **Suppl. Fig. 23D**). Furthermore, we observed a strong anti-correlation between the Areg *versus* “white fat cell differentiation” signatures inferred from all the considered samples (**Fig. 4F**, **Suppl. Fig. 23A** and **C**, **Suppl. Table 1**). Specifically, we found that the expression of most top Areg markers, and particularly those of all tested candidates, strongly anti-correlated with the “white fat cell differentiation score” (**Fig. 4G**, **Suppl. Fig. 23E**, **Methods**). Our findings therefore point to a strong association between Areg marker (**Suppl. Table 1**) expression (especially of the tested candidates) and the inability of ASPCs to undergo adipogenic differentiation.

Finally, we aimed to provide additional evidence that the effect of functional Aregs on differentiating CD142− ASPCs is at least in part regulated by RA signalling-related genes. Remarkably, the “cellular response to RA” (GO:0071300) and “RA receptor signalling pathway” (GO:0048384) terms were enriched in CD142− ASPCs subjected to “active Areg” signalling (**Suppl. Fig. 23F-G**), and the expression of the genes involved in these two terms were anti-correlated with “white fat cell differentiation score” (**Fig. 4G-I, Suppl. Fig. 23A, F** and **G**). To further demonstrate the involvement of RA signalling in this Areg-mediated inhibitory effect, we performed bulk RNA-seq of CD142− ASPCs treated with RA or EGF in order to compare their gene expression profiles to those of CD142− ASPCs that were co-cultured with “active Aregs”. This analysis revealed that CD142− ASPCs that were exposed to “active Aregs” were transcriptionally more similar to CD142− ASPCs treated with RA than those treated with EGF (**Suppl. Fig. 24**). In addition, we observed that the receiving CD142− ASPCs exhibited remarkably coherent transcriptional dynamics of a number of genes and pathways that were previously reported to be involved in the RA-mediated inhibition of adipogenesis, a pattern of expression supported by the analysis of CD142− ASPCs treated with RA. Specifically, Wnt signalling pathway-related terms, RA receptors (*Rarb* and *Rarg*) as well as Catenin beta-1 (*Ctnnb*) were enriched upon the Areg-mediated inhibition of differentiating CD142− ASPCs (**Suppl. Fig. 25**), consistent with previously reported findings on the effect of RA on preadipocytes (Goldstein, Scalia and Ma, 2009; Kim *et al.*, 2013). Moreover, CD142− ASPCs exposed to the secretome of CD142+ ASPCs during adipogenic differentiation showed a decreased expression (compared to CD142− ASPCs being exposed to “dysfunctional” or no Aregs) of a substantial collection of genes reported to be downregulated in RA-mediated adipogenesis suppression, namely *Rxra*, (Sagara *et al.*, 2013), *Pparg*, *Cebpa* (Schwartz *et al.*, 1996), *Mapk14* (Lee *et al.*, 2011), *Zfp423* (Wang *et al.*, 2017) and *Asct2* (Takahashi *et al.*, 2015) (**Suppl. Fig. 25**). Conversely, the expression of genes involved in the EGF-mediated inhibition of adipogenesis, *Erk1* (MAPK3), *Erk2* (MAPK1) and *Prkaca* (PKA C alpha) (Boney, Smith and Gruppuso, 1998; MacDougald and Mandrup, 2002; Harrington, Pond-Tor and Boney, 2007) did not show substantial changes in CD142− ASPCs that were exposed to “active Aregs” compared to CD142− ASPCs exposed to “dysfunctional Aregs” (**Suppl. Fig. 25**). Finally, we found that RA-treated CD142− ASPCs, next to an impaired adipogenic capacity (**Fig. 3J-K**), exhibit also a strikingly consistent transcriptomic signature, with a number of highly specific Areg markers being regulated by RA in a concentration-dependent manner. Indeed, we observed that the expression of *F3* and *Mgp* but also of *Cpe*, *Gdf10*, *Bgn* and *Clec11a* and other Areg-specific genes is gradually upregulated by increasing concentrations of RA (**Fig. 4J, Suppl. Fig. 26**). However, for a number of genes (*F3*, *Mgp*, *Bgn*, *Clec11a*), RA administered at concentrations higher than 10 μM tended to reverse this up-regulation, potentially pointing to a specific RA concentration range (0.01-1 μM) that may be physiologically relevant for the Areg-dependent inhibition of adipogenesis. Together, these findings strongly suggest that the RA signalling pathway plays an important role in mediating the inhibitory activity of Aregs through a synchronised regulation of Areg-specific genes.

## Discussion

In this study, we systematically examined the molecular and phenotypic properties of murine subcutaneous CD142+ ASPCs, motivated by the reported discrepancy regarding their behaviour in the context of adipogenesis (Schwalie *et al.*, 2018; Hwang and Kim, 2019; Merrick *et al.*, 2019; Corvera, 2021). Using numerous functional and multi-omic assays across distinct experimental settings and sampling conditions, we unambiguously validated the previously proposed non-adipogenic and anti-adipogenic nature of these cells, which is why we suggest to retain their initially proposed name: “Aregs” for adipogenesis regulators (**Fig. 1** and **Suppl. Fig. 1-9**).

These analyses also led to unexpected findings regarding in particular the age-dependent nature of CD142+ ASPCs’ functional properties. It has been shown that the developmental timing of adipose tissue formation varies largely between species (Carberry, Colditz and Lingwood, 2010; Louveau *et al.*, 2016) and different anatomical fat depots (Birsoy *et al.*, 2011; Han *et al.*, 2011; Rosen and Spiegelman, 2014; Hong *et al.*, 2015). Subcutaneous stromal vascular fraction (SVF) cells were shown to be capable of differentiating into lipid-filled adipocytes *in vitro* under adipogenic differentiation medium from embryonic day E16.5 (Birsoy *et al.*, 2011). However, the dynamics of post-natal ASPC differentiation, as well as the emergence of their cellular heterogeneity, are still poorly understood. Having experimentally investigated diverse murine post-natal developmental stages, we found that all fractions of newborn (P0) ASPCs displayed a molecular identity and behaviour (high proliferative and adipogenic propensity) that resemble those of “naïve preadipocytes” (**Suppl. Fig. 12**) (Wang *et al.*, 2003; Atanassova, Rancic and Georgieva, 2012; Hong *et al.*, 2015; Hepler and Gupta, 2017). Importantly, P0-derived CD142+ ASPCs did not show a higher “Areg/CD142+ score” (**Suppl. Table 1, Fig. 2G-H**, **Suppl. Fig. 14**) nor higher expression of “retinol metabolic process”-related genes (**Suppl. Fig. 13**) compared to the other tested ASPC fractions. This is consistent with the notion that all P0 ASPCs are likely still naïve and indicates that the mesenchymal cellular landscape that is observed in adults has not yet been established in newborns. Further inquiry revealed an age-dependent evolution of both the molecular signature as well as the diverse adipogenic phenotypes of ASPCs. Indeed, we found that the Areg-specific molecular characteristics emerge in CD142+ ASPCs before P16, consistent with their detection in P12 scRNA-seq data (Merrick *et al.*, 2019) and become most prominent in adulthood (**Fig. 2A**, **Suppl. Fig. 10**). Interestingly however, the non- and anti-adipogenic phenotype of CD142+ ASPCs only emerged between post-natal day 16 and 28 (4wo), thus after the establishment of their molecular identity, with, intriguingly, the weaning of the litters occurring during this time period (**Fig. 2D-E**). We cannot formally point to weaning as the primary causal factor for the observed timing offset between Aregs’ molecular and functional appearance. Yet, the fact that weaning entails a dramatic nutritional alteration makes it an intriguing candidate for further investigation.

While the observed adipogenic phenotype of pre-weaning CD142+ ASPCs is rather striking, it still does not fully resolve the reported functional discrepancy for CD142+ ASPCs given that an adipogenic propensity has also been reported for adult CD142+ APSCs (Merrick *et al.*, 2019), which contrasts with our results. Amongst remaining possible reasons for this discrepancy is genetic background. Indeed, Merrick and colleagues used CD1 as opposed to C57BL/6J mice used in other studies, including our own (Burl *et al.*, 2018; Hepler *et al.*, 2018; Schwalie *et al.*, 2018; The Tabula Muris Consortium *et al.*, 2018; Cho, Lee and Doles, 2019; Zhang *et al.*, 2019; Sárvári *et al.*, 2021). Metabolic variation as a function of genetic background is widely recognized in the field (Fontaine and Davis, 2016). Yet, how ASPC heterogeneity and function may vary across individuals / strains has not yet been investigated and will thus constitute an exciting downstream research avenue.

Given the demonstrated non-adipogenic and inhibitory properties of Aregs, understanding how these functional properties are molecularly regulated is highly relevant. Since our findings demonstrated that Aregs exert their inhibitory properties *via* paracrine signalling, we focused on genes coding for secreted factors. We identified six candidates that were specific to Aregs across multi-omic datasets: *F3* (coding for CD142) itself, *Mgp*, *Gdf10*, *Clec11a*, *Cpe* and *Bgn*, with CD142 and MGP the most interesting functionally, based on recombinant protein as well as knockdown assays (**Fig. 3F-G**, **Suppl. Fig. 16**, **Fig. 4A-B**). The involvement of CD142 in Aregs’ inhibitory activity is surprising given the reported physiological role of CD142 as a coagulation factor (Chu, 2011). CD142, also known as Tissue factor, is the primary initiator in the extrinsic coagulation pathway (Petersen, Valentin and Hedner, 1995) and has not been explicitly shown to be involved in adipogenesis-related processes. MGP (Matrix Gla protein, a member of a family of vitamin-K2 dependent, Gla-containing proteins) has been demonstrated to act as an inhibitor of calcification in cartilage and vasculature (Bäck *et al.*, 2019), implying its possible specificity to mesenchymal cells with multilineage potential. Finally, while the inhibitory effect of GDF10 has been previously demonstrated in the context of adipogenesis in Areg-like/CD142+ muscle-resident stromal cells (Camps *et al.*, 2020) as well as in differentiating 3T3-L1 cells (Hino *et al.*, 2012), we found that inactivating *Gdf10* in Aregs did not majorly interfere with their inhibitory activity towards other ASPCs.

Next to these secretory proteins, we uncovered the RA-signalling pathway as another likely actor that is involved in the Areg-mediated inhibition of adipogenesis. Retinoic acid has long been linked to adipogenic inhibition ( Murray and Russell, 1980; Kuri-Harcuch, 1982; Salazar-Olivo *et al*., 1994; Schwartz *et al.*, 1996; Lee *et al.*, 2011; Sagara *et al.*, 2013; Wang *et al.*, 2017) and demonstrated to be protective against diet-induced obesity (Berry and Noy, 2009; Bonet, Ribot and Palou, 2012). Throughout our analyses, we identified a substantial number of genes related to RA signalling to be specific to Aregs and some of them, including *Aldh1a2*, *Epha3, Osr1* and *Rbp1* are *bona fide* Areg markers (**Fig. 3H**, **Suppl. Fig. 18**). Furthermore, transcriptomic profiling of freshly isolated Aregs suggests that they produce retinol (**Fig. 4I**, **Suppl. Fig. 19A-B**), a precursor of RA. In addition, RA-related terms, particularly “cellular response to RA” and “RA receptor signalling pathway” were enriched in CD142− ASPCs that were subjected to active Areg signalling (**Fig. 4I**, **Suppl. Fig. 4A**, **Suppl. Fig. 19C-D**). Together, these results suggest that Aregs might exert their anti-adipogenic activity *via* RA itself, as further supported by the remarkably coherent transcriptional dynamics of a set of genes involved in RA-mediated inhibition in CD142− ASPCs exposed to active Aregs. For example, we found that Areg-treated CD142− ASPCs were transcriptionally more similar to CD142− ASPCs treated with RA than those treated with EGF, another well-established adipogenesis inhibitor (**Fig. 4J**, **Suppl. Figs. 24-25**) (Harrington, Pond-Tor and Boney, 2007). This transcriptional similarity reflects a more general, intriguing phenomenon in that Areg-treated CD142− ASPCs exhibited a significantly increased expression of many Areg-specific genes, a number of which were also upregulated in RA-treated CD142− ASPCs (**Fig. 4D-G**, **Suppl. Fig. 26**). These results point to a potential conversion of CD142− ASPCs, when subjected to inhibitory signals, into Areg-like cells, a phenomenon of interconversion of various ASPC subpopulations that has been recently proposed to occur within the subcutaneous adipogenic stem cell niche (Merrick *et al.*, 2019). It is thereby worth noting that the transcriptome of Aregs exposed to adipogenic cocktail also showed a response to RA (**Suppl. Fig. 19C-D**), suggesting that they exert an auto-inhibitory effect that may impair their own ability of generating *in vitro* adipocytes.

In conclusion, our findings systematically authenticated Aregs as a molecularly, phenotypically and functionally robust inhibitory subpopulation of murine subcutaneous ASPCs. This subpopulation emerges within the ASPCs during early stages of post-natal development. The establishment of the molecular signature of Aregs precedes the manifestation of their phenotypic (non-adipogenic) and functional (anti-adipogenic) properties, with those two events being separated by weaning. We finally uncovered that the murine Areg-mediated inhibition of adipogenesis involves the secreted factors CD142 (encoded by *F3*) and Matrix Gla protein (MGP), which act together with the RA signalling pathway. This set of factors seems at first glance unrelated, but intriguingly, the expression of *F3* and *Mgp* (as well as *Aldh1a2* and *Bgn)*, has already been shown to be regulated by retinoic acid (Balmer and Blomhoff, 2002; Takeda *et al.*, 2016) as we also demonstrated for these and other Areg-specific markers (**Fig. 4J, Suppl. Fig. 26**). Furthermore, these genes were reported to be higher expressed in visceral compared to subcutaneous ASPCs and have as such been associated with the impaired capacity of visceral ASPCs to give rise to *in vitro* adipocytes (Reichert *et al.*, 2011; Meissburger *et al.*, 2016; Takeda *et al.*, 2016; Li *et al.*, 2020). Further studies will now be required to investigate how this seemingly diverse collection of molecules may cooperate within the retinoic acid signalling pathway to steer the developmental, phenotypic and functional properties of Aregs.

## Acknowledgements

We thank W. Chen, G. van Mierlo and Judith F. Kribelbauer for constructive discussions and careful reading of the manuscript. This research was supported by the Swiss National Science Foundation Grants (#31003A_162735, 31003A_182655, and CRSII5_186271), the Precision and Health-related Technologies Initiative Grants (PHRT #222, #307, and #502) and by institutional support from the Swiss Federal Institute of Technology in Lausanne (EPFL). We thank the EPFL Core Facilities: CPG (Centre de phénogénomique, especially Arnaud Legay), FCCF (Flow Cytometry Core Facility, especially Miguel Garcia), BIOP (BioImaging and Optics Platform, especially Olivier Buri and Romain Guiet), PCF (Proteomics Core Facility, especially Romain Hamelin and Florence Armand), GECF (Gene Expression Core Facility, especially Bastien Mangeat).

## Author contributions

B.D., M.Z. and P.Y.R. designed the study and wrote the manuscript. M.Z. and P.Y.R. conducted the experiments and analyses: M.Z. performed all the experimental assays including murine SVF extraction, FACS-based isolation, cell culture experiments, imaging, image analyses and quantification. P.Y.R. performed all single-cell-, bulk RNA-sequencing- and mass-spectrometry-related analyses. J.R. and D.A. performed bulk RNA sequencing and data pre-treatment, H.H., R.F. and M.L. assisted with experimental procedures and analyses. All authors read and approved the final manuscript.

