## Supplementary Materials for "Mammalian adipogenesis regulators (Aregs) exhibit robust non- and anti-adipogenic properties that arise with age and involve retinoic acid signalling"

1. Methods
2. Supplementary Figures 1-27
3. Supplementary Tables 1-4

### Methods

#### Bioethics

All mouse experiments were conducted in strict accordance with the Swiss law and all experiments were approved by the Ethics Commission of the state veterinary office (VD2984/2015, VD3406/2018).

#### Analysis and integration of single-cell RNA-seq data

<u>Integration of adult murine subcutaneous adipose tissue scRNA-seq data</u>

The integration of the three scRNA-seq datasets of adult murine subcutaneous adipose tissue published by (Burl *et al.*, 2018; Schwalie *et al.*, 2018; Merrick *et al.*, 2019) was performed as described in (Ferrero, Rainer and Deplancke, 2020). The clustering tree plot was generated using the function clustree() from the package *clustree* version 0.4.3. (Zappia and Oshlack, 2018).

<u>Analysis and Integration of P12 murine subcutaneous adipose tissue scRNA-seq data</u>

The scRNA-seq count matrices from Merrick *et al.* (Merrick *et al.*, 2019) (GSM3717977) were downloaded from the Gene Expression Omnibus (GEO) repository. The datasets were first analysed separately to identify the main clusters as described in (Ferrero, Rainer and Deplancke, 2020). Differential expression analysis between the different clusters was performed using the function FindAllMarkers() from *Seurat* (Stuart *et al.*, 2019). Only the markers detected as significant (FDR < 0.05 and logFC > 0.25, as defined by default) by both the Wilcoxon Rank Sum test (test.use = “wilcox”) and Likelihood-ratio test for single cell gene expression (test.use = “bimod”) were selected. This dataset was then integrated with the three adult datasets discussed above following similar methods as the one described in (Ferrero, Rainer and Deplancke, 2020). The “CD142+ score” was calculated using the function AddModuleScore() of *Seurat* (Stuart *et al.*, 2019) giving as features the top CD142+ markers (see “Selection of CD142+ top 100 markers”) (**Fig. 2A** and **Supp. Fig. 10E**).

<u>Comparison of male and female ASPC scRNA-seq data</u>

The single-cell RNA-seq dataset published in (Schwalie *et al.*, 2018) (https://www.ebi.ac.uk/arrayexpress/experiments/E-MTAB-6677) contains both male and female cells. A strict filtering based on the expression of Y-chromosome genes (*Ddx3y*, *Eif2s3y*, *Kdm5d*, *Uty*, and *Sult1e1*) and the local minimum of *Xist* expression was used to separate male and female cells. More precisely, female cells were defined as cells expressing *Xist* above the defined threshold and not expressing any of the Y-chromosome genes, and male cells were defined as cells expressing at least one of the Y-chromosome genes and not *Xist*. The other cells were filtered out. The two groups, female and male cells, were analyzed separately. For each, genes not expressed in at least three cells, were filtered out. The data were log normalized using the functions ComputeSumFactor() to which the results of the function quickCluster() was given as input to the parameter clusters, and the function normalize() from the package *SCRAN M3Drop* (Andrews and Hemberg, 2019) was used to find the highly variable genes. A *Seurat* object was created and the data were scaled for the number of features and number of UMIs. The PCA was computed, and the first x significant PCs defined by the JackStraw test implemented in *Seurat* (Stuart *et al.*, 2019) were used to build the tSNE. Clustering was performed using the function sc3() from the package *SC3* (Kiselev *et al.*, 2017). Differential expression analysis between the different clusters was performed using the function FindAllMarkers() from *Seurat* (Stuart *et al.*, 2019). Only the markers detected as significant (FDR <0.05 and logFC > 0.25, as defined by default) by both the Wilcoxon Rank Sum test (test.use = “wilcox”) and Likelihood-ratio test for single cell gene expression (test.use = “bimod”) were selected. The top differentially expressed genes of males were ranked by logFC and the top 100 were used to generate **Suppl. Fig. 10A**.

<u>Comparison of top markers between scRNA-seq results</u>
**Suppl. Fig. 9B** displays the percentage of the top 100 markers shared between the different subpopulations that are displayed. The male markers were identified using differential expression analysis as described in “Comparison of male and female ASPC scRNA-seq data”. For the left barplot, the male markers were compared to the top 100 markers published in (Schwalie *et al.*, 2018). For the right bar plot, the male markers were compared to the top 100 markers identified for the three main ASPC subpopulations based on the integration analysis that we performed as described in “Integration of adult murine subcutaneous adipose tissue scRNA-seq data”. **Suppl. Fig. 10A** shows the percentage of the top 60 markers identified for each of the three main ASPC subpopulations in P12 as described in “Analysis and Integration of p12 murine subcutaneous adipose tissue scRNA-seq data”, compared to the top 60 adult markers (ranked by logFC) of the same subpopulations identified based on the integration analysis that we performed, as described in “Integration of adult murine subcutaneous adipose tissue scRNA-seq data”. For this figure, only the top 60 markers were used as one of the P12 ASPC subpopulations did not have more than 60 markers that were defined as differentially expressed based on our filtering, which is not uncommon in scRNA-seq analyses.

**Selection of CD142+ top 100 markers**
Three single-cell RNA-seq datasets of subcutaneous murine fat tissues from (Burl *et al.*, 2018; Schwalie *et al.*, 2018; Merrick *et al.*, 2019) were integrated using the *Seurat* (Stuart *et al.*, 2019) pipeline as described in the supplementary Methods of (Ferrero, Rainer and Deplancke, 2020). The differentially expressed genes of the CD142 (*F3*+) cluster were the genes identified as differentially expressed in all three batches and with a corrected p-value below 0.05. More precisely, the cells of the integrative CD142 (*F3*+) cluster were sorted by batch. Within each respective dataset, DE genes were identified using the FindMarkers() function of *Seurat* (Stuart *et al.*, 2019) with the Wilcox test (logfc.threshold = 0.25, min.pct = 0.1). Only genes identified as differentially expressed in all the batches and with a positive log fold change were considered. The p-values were corrected using the Fisher() function from the *metaseqR* package. 85 genes of this list had an adjusted pvalue < 0.05. The top 20 differentially expressed genes in CD142+ *versus* CD142− ASPCs from bulk RNA-seq data published in (Schwalie *et al.*, 2018) were added, yielding a final list of 100 top markers, referred to as the “Top CD142+ markers” in the manuscript (**Suppl. Table 1)**.

**Isolation of the mouse stromal vascular fraction (SVF)**
Subcutaneous inguinal adipose tissue depots were dissected from wild-type C57BL/6J male and female mice of different ages (P12-17 (12-17 days post-birth), 4-week-old, 7-week-old, 11-week-old) and transferred into PBS. The tissues were subsequently minced using scissors, transferred into collagenase (Sigma-Aldrich #C6885-1G, 2 mg/ml of homemade collagenase buffer (25 mM $NaHCO_3$, 12 mM $KH_2PO_4$, 1.2 m $MgSO_4$, 4.8 mM KCl, 120 mM NaCl, 1.4 M $CaCl_2$, 5 mM Glucose, 2.5% BSA, pH = 7.4)) and incubated for 65-70 min at 37 °C under

agitation (130 rpm on a vertical shaker), after which they were resuspended after 45-50 min of digestion. The cell suspension was then diluted with PBS (with the aim of diluting the collagenase and stopping/slowing down the digestion), and filtered through a 100-μm, then 40-μm cell strainer in order to generate a single cell suspension. Next, the cells were pelleted by a 5 min centrifugation at 400 g at room temperature and red blood cells were lysed by incubating the pellet with a homemade red blood cell lysis buffer (154 mM NH4Cl, 10 mM KHCO3, 0.1 mM EDTA) for a maximum of 5 min, followed by two washes (a 5 min centrifugation at 400 g, room temperature) with a homemade FACS buffer (PBS with 3% foetal bovine serum (FBS) (Gibco #10270-106), 1 mM EDTA, 1% penicillin–streptomycin (Gibco #15140122)).

**Isolation of SVF cells from newborn mice**
Burgeoning depots of inguinal subcutaneous adipose tissue were dissected from newborn (P0) wild-type C57BL/6J mice and transferred into PBS. The tissues were then transferred into collagenase solution (see above) and incubated for 1 h at 37 °C under agitation (130 rpm on a vertical shaker), after which they were resuspended after 45 min of digestion. The cell suspension was then filtered through a 100-μm, then 40-μm cell strainer in order to generate a single cell preparation. Next, the cells were pelleted by a 5-min centrifugation at 400 g at room temperature and a red blood cell lysis was performed by incubating the pellet in a homemade red blood cell lysis buffer (see above) for 5 min, followed by two washes (a 5-min centrifugation at 400 g, room temperature) with a homemade FACS buffer (see above).

**FACS-based cell isolation**
The SVF cell suspension was diluted to 1 × 10^7 cells/ml with a homemade FACS buffer (see above) and the following fluorophore-conjugated antibodies were added (in titration-determined quantities, **Suppl. Table 3**): anti-mouse CD31–AF488, anti-mouse CD45–AF488, anti-mouse TER119–AF488 (BioLegend #303110, #304017 and #116215, respectively) to select the Lin− population; anti-mouse SCA1-PE-Cy7 (BioLegend #122513) to enrich the Lin− population with ASPCs; and four different anti-mouse CD142 antibodies: 1) anti-mouse CD142 SinoBiological (SB), monoclonal rabbit IgG clone #001, PE-conjugated, #50413-R001-PE, 2) SinoBiological, monoclonal rabbit IgG clone #001, #50413-R001, in-house conjugated to Lightning-Link PE with Conjugation Kit (Innova Biosciences #703-0010) (SB-LL), 3) BiOrbyt, monoclonal mouse Igg clone HTF-1, PE-conjugated, # ORB507485 (BO), 4) R&D Systems, polyclonal goat IgG, PE-conjugated, #FAB3178P (RnD).

The cells were incubated with the antibodies on ice for at least 30 min and protected from light, after which they were washed with the FACS buffer, stained with propidium iodide (Molecular Probes #P3566) for assessing viability, filtered through a 40-μm cell strainer and subjected to FACS with the use of a Becton Dickinson FACSAria II sorter. Compensation measurements were performed for single stains using positive/negative compensation beads (eBiosciences #01-2222-42).

The following gating strategy was applied while sorting the cells: first, the cells were selected based on their size and granularity/complexity (side and forward scatter), followed by a double elimination of any events that could represent more than one cell. Next, the Lin−(CD31− CD45− TER119–) population was selected, followed by a Lin−SCA1+ selection (ASPCs), and finally, ASPC cells that were negative or positive for CD142 marker: Lin− SCA1+ CD142− (CD142− ASPCs) and Lin− SCA1+ CD142+ (CD142+ ASPCs, Aregs). The measurements were acquired using Diva software supplemented on the Becton Dickinson FACS Aria II sorter and analysed using FlowJo analysis software.

##### *In vitro* adipogenic differentiation

Equal cell numbers were sorted or plated in flat-bottom microscopy-adapted cell culture plates (Corning #353219), cultured in high glucose DMEM medium (Gibco #61965026) supplemented with 10% FBS and 1% penicillin–streptomycin, and treated with a single dose of white adipocyte differentiation induction cocktail at confluence (0.5 µM 3-isobutyl-1-methylxanthine (IBMX, Sigma #15879), 1 µM dexamethasone (Sigma #D2915), 170 nM (1µg/ml) insulin (Sigma #19278)), followed by a maintenance treatment two-three days post-induction (170 nM insulin) and refreshed every two-three days. The culture was carried out until day six-eight after induction, at which point cells were stained for imaging or collected for RNA extraction.

##### *In vitro* adipogenic differentiation using various differentiation cocktails

Equal cell numbers were sorted or plated as described above. At confluence, the cells were treated with a single dose of one of four different versions of white adipocyte differentiation induction cocktails: 1) the classical complete cocktail in DMEM medium: 0.5 µM IBMX, 1 µM dexamethasone, 170 nM (1µg/ml) insulin; 2) minimal cocktail in DMEM medium: 170 nM insulin; 3) Complete cocktail + Indomethacin + T3 in DMEM/F12 (1:1 ratio) medium: 0.5 µM IBMX, 1 µM dexamethasone and 20 nM, 125 nM indomethacin, 1 nM Triiodothyronine (T3); 4) Minimal cocktail in DMEM/F12 medium: 20 nM insulin. After 2-3 days of induction, maintenance cocktail was applied every 2-3 days with 1) 170 nM insulin in DMEM for cocktail 1); 2) DMEM for cocktail 2); 3) 20 nM insulin in DMEM/F12 for cocktail 3); 4) DMEM/F12 for cocktail 4). DMEM for all the corresponding treatments was supplemented with 10% FBS and 1% penicillin–streptomycin. DMEM/F12 for all the corresponding treatments was supplemented with 10% FBS and 50 ng/ml Primocin. The culture was maintained till day 6-8 post-induction.

##### Imaging and quantification of *in vitro* adipogenesis

Once the cells differentiated fully, they were stained with live fluorescence dyes: Bodipy (boron-dipyrromethene, Invitrogen #D3922) for lipids and Hoechst for nuclei. Cells were incubated with the dyes in FluoroBrite, phenol red-free DMEM medium (Gibco #A1896701), supplemented with 10% FBS and 1% penicillin–streptomycin for 30 min at 37 °C in the dark, washed twice with PBS and imaged in FluoroBrite medium. Given the substantial variation in the extent of lipid accumulation across the tested cell fractions (within the same well but also across technical replicates), the imaging was optimised to cover the largest surface of the well possible. Moreover, a z-stack acquisition in a spinning-disc mode and Z-projection were performed in order to capture the extent of *in vitro* adipogenesis with the highest possible accuracy (**Suppl. Fig. 27**). Specifically, the automated platform Operetta (Perkin Elmer) was used for imaging. First, 7-8 z-stacks were acquired for every field of view (**Suppl. Fig. 27A**) in a confocal-mode of the microscope in order to produce high quality images for downstream z-projection and accurate thresholding (**Suppl. Fig. 27B**). Next, 25 images per well were acquired (**Suppl. Fig. 27C**) using a Plan Neofluar 10x Air, NA 0.35 objective, with 10% overlap for further tiling and with the aim of covering most of each well's surface to assure an accurate representation of overall lipid accumulation (**Suppl. Fig. 27D**). (For experiments in **Fig. 4A** and **Suppl. Fig. 22H**, 12 images per well were acquired using a Plan Apochromat 10x Air, NA 0.8 objective, with 10% given the use of a different cell culture plate (see "Transwell experiments"). The Hoechst, blue, channel was acquired with 40 ms exposure time and 50% of the illumination power while the Bodipy, green, channel was acquired with 20 ms exposure time and 50% of the illumination power. The images, supported by Harmony software, were exported as TIFF files. They were subsequently tiled using the acquired 10% image overlap and Z-projection was performed with the maximum intensity of acquired z-stacks. To accurately estimate and represent differences in adipogenesis, a quantification algorithm for image treatment was developed in collaboration with the EPFL BIOP imaging facility. In brief, image analysis was performed in ImageJ/Fiji, lipid droplets (yellow) and nuclei (blue) images were filtered using a Gaussian blur (sigma equal to 2 and 3, respectively) before an automatic thresholding. The automatic thresholding algorithm selections were chosen on the basis of visual inspection of output images. The areas corresponding to the thresholded lipid and nuclei signals were then used to calculate the normalised lipid accumulation (Adiposcore) for each examined sample. In the main figures, representative blown-up crop images of each sample are shown. In the supplementary figures, tiled (25 images) and thresholded (both lipid - yellow and nuclei - blue channels) images are shown. Plotted ratios were normalised to the indicated control in all figures except **Fig. 2E** and the corresponding **Suppl. Fig. 11D**, where non-normalised values from 13 independent experiments were plotted together, which explains why the observed variation is more substantial.

**Mass spectrometry sample preparation**

FACS-sorted cells were vacuum-centrifuged to near dryness and resuspended in 9µl of 100 mM HEPES pH8, 10mM tris(2-carboxyethyl)phosphine. Samples were first heated for 20 min at 95°C with permanent shaking and then sonicated in a water bath for 15 min. Extracted proteins were alkylated with 1 µl of 400 mM chloroacetamide for 30 min at 37°C in the dark with permanent shaking. Proteins were digested overnight using 200 ng mass spectrometry grade trypsin with permanent shaking. Resulting peptides were desalted on C18 StageTips (Rappsilber, Mann and Ishihama, 2007) and dried by vacuum centrifugation. For TMT labelling, peptides were first reconstituted in 8 µl HEPES 100 mM pH 8 and 3 µl of TMT solution (20 µg/µl in pure acetonitrile) was then added. Labelling was performed at room temperature for 1.5 h and reactions were quenched with hydroxylamine to a final concentration of 0.4% (v/v) for 15 min. TMT-labelled samples were then pooled at a 1:1 ratio across all samples. A single shot control LC-MS run was performed to ensure similar peptide mixing across each TMT channel to avoid the need of further excessive normalization. Quantities of each TMT-labelled sample were adjusted according to the control run. The combined sample was vacuum-centrifuged and fractionated into 8 fractions using the Pierce High pH Reversed-Phase Peptide Fractionation Kit following the manufacturer's instructions. Resulting fractions were dried by vacuum centrifugation and again desalted on C18 StageTips.

Each individual fraction was resuspended in 2% acetonitrile, 0.1% FA and nano-flow separations were performed on a Dionex Ultimate 3000 RSLC nano UPLC system on-line connected with a Lumos Fusion Orbitrap Mass Spectrometer. A capillary precolumn (Acclaim Pepmap C18, 3 µm-100Å, 2 cm x 75 µm ID) was used for sample trapping and cleaning. Analytical separations were performed at 250 nl/min over 150 min biphasic gradients on a 50 cm long in-house packed capillary column (75 µm ID, ReproSil-Pur C18-AQ 1.9 µm silica beads, Dr. Maisch). Acquisitions were performed through Top Speed Data-Dependent acquisition mode using a 3 seconds cycle time. First MS scans were acquired at a resolution of 120'000 (at 200 m/z) and the most intense parent ions were selected and fragmented by High energy Collision Dissociation (HCD) with a Normalized Collision Energy (NCE) of 37.5%

using an isolation window of 0.7m/z. Fragmented ion scans were acquired with a resolution of 50'000 (at 200 m/z) and selected ions were then excluded for the following 120s.

**Mass spectrometry data analysis**

Raw data were processed using SEQUEST, Mascot, MS Amanda (Dorfer *et al.*, 2014) and MS Fragger (Kong *et al.*, 2017, p. 20) in Proteome Discoverer v.2.4 against the Uniprot Mouse Reference Proteome (Uniprot Release: 2020_05). Enzyme specificity was set to Trypsin and a minimum of six amino acids was required for peptide identification. Up to two missed cleavages were allowed and a 1% FDR cut-off was applied both at peptide and protein identification levels. For the database search, carbamidomethylation (C), TMT tags (K and Peptide N termini) were set as fixed modifications whereas oxidation (M) was considered as a variable. Resulting text files were processed through in-house written R scripts (version 3.6.3). Two steps of normalization were applied. The first step of normalization was the sample loading normalization (Plubell *et al.*, 2017). Assuming that the total protein abundances were equal across the TMT channels, the reporter ion intensities of all spectra were summed and each channel was scaled according to this sum, so that the sum of reporter ion signals per channel equals the average of the signals across samples. Then, the Trimmed M-Mean normalization step was also applied using the package *EdgeR* (Robinson, McCarthy and Smyth, 2010) (version 3.26.8). Assuming that the samples contain a majority of non-differentially expressed proteins, this second step calculates normalization factors according to these presumed unchanged protein abundances. Proteins with high or low abundances and proteins with larger or smaller fold-changes were not considered. Differential protein expression analysis was performed using the R bioconductor package *limma* (version 3.34.9, 2018-02-22) (Ritchie *et al.*, 2015), followed by the Benjamini-Hochberg multiple-testing method (Benjamini and Hochberg, 1995). Adjusted P values lower than 0.05 (FDR $< 0.05$) and absolute $\log_2$FC $> 1$ were considered as significant. GSEA enrichment analysis (**Supp. Fig. 19B**) was performed using the gseGO() function on the proteins ranked by the logFC of protein expression in freshly isolated CD142+ *versus* CD142− ASPCs.

**Recombinant protein and retinoic acid treatment**

Equal cell numbers were sorted or plated in flat-bottom microscopy-adapted cell culture plates and cultured as described above. At confluence, the cells were treated with a single dose of classical white adipocyte differentiation induction cocktail at half of its usual concentration: 0.25 µM IBMX, 0.5 µM dexamethasone, 85 nM (0.5 µg/ml) insulin (in DMEM supplemented with 10% FBS and 1% penicillin–streptomycin), supplemented with the following recombinant proteins: BSA (Sigma, # A3059, A9418), EGF (ThermoFisher Scientific, # PMG8043), CD142 (LifeSpan BioSciences, # LS-G12283), MGP (LifeSpan BioSciences, #LS-G13865), CLEC11A (R&D Systems, #3729-SC), GDF10 (SinoBiological, #50165-M01H), CPE (ArcoBioSystems, #CAE-M5222) and BGN (R&D Systems, #8128-CM) at various concentrations including all or some of the following concentrations: 10, 50, 100, 500, 750 and 1000 ng/ml. For retinoic acid treatments, the above described differentiation and maintenance cocktails were supplemented with retinoic acid (Sigma, # R2625) diluted in Dimethyl sulfoxide (DMSO, AppliChem, #A3672,0250). The corresponding doses of the recombinant proteins and the retinoic acid, were applied after 2-3 days of induction with the maintenance medium at half of its usual concentration, 85 nM (0.5 µg/ml) insulin in DMEM supplemented with 10% FBS and 1% penicillin–streptomycin.

**siRNA-mediated knockdown experiments**

For each gene, a pool of 3 siRNA probes (IDT, TriFECTA DsiRNAs, **Suppl. Table 4**) was reverse-transfected to CD142+ ASPCs (Aregs) or total ASPCs. Seventy-five thousand cells/cm2 were plated into the well with previously deposited siRNA mix containing 20 nM of each siRNA re-suspended in 1.5% Lipofectamine RNAiMAX (Invitrogen #13778150) dissolved in Opti-MEM I reduced serum medium (Invitrogen #31985062), and high glucose DMEM medium supplemented with 2.5% FBS (w/o penicillin–streptomycin). After 24 h, the medium was changed to high glucose DMEM medium supplemented with 10% FBS and 1% penicillin–streptomycin and after 48 h, the cells were collected for determining knockdown efficiency (see 'RNA isolation and quantitative PCR') or induced with white adipogenic differentiation cocktail (see '*In vitro* adipogenic differentiation').

**Transwell experiments**

Wild-type CD142+ ASPCs or those carrying the indicated gene knockdown constructs (knockdowns performed in the transwell, see "siRNA-mediated knockdown experiments") were plated in the transwell inserts (Corning #3381). 24 h after transfection, the cells in inserts were washed with PBS and changed to high glucose DMEM medium supplemented with 10% FBS and 1% penicillin–streptomycin. 48 h post-transfection, the inserts were placed above previously plated CD142− ASPCs (cultured in the transwell-receiving cell culture Corning #3382 plates) and both cell populations were treated with white adipocyte differentiation cocktail (see '*In vitro* adipogenic differentiation') and imaged at day 6-8 after induction (see 'Imaging and quantification of *in vitro* adipogenesis').

**RNA isolation and quantitative PCR**

Cells carrying siRNA-mediated knockdown constructs were collected into Tri-Reagent (Molecular Research Center #TR118) 48 h post-transfection. Direct-zol RNA kit (Zymo Research #R2052) was used to extract RNA, followed by reverse transcription using the SuperScript VILO cDNA Synthesis Kit (Invitrogen). Expression levels of mRNA were assessed by real-time PCR using the PowerUp SYBR Green Master Mix (Thermo Fisher Scientific #A25743). mRNA expression was normalized to the *Hprt1* gene.

**Bulk mRNA-seq**

3'end bulk mRNA sequencing was performed following the BRB-seq strategy as previously described (Alpern *et al.*, 2019). In brief, 7-200 ng of total RNA from each sample was reverse transcribed in a 96-well plate using SuperScriptTM II Reverse Transcriptase (Lifetech 18064014) with individual barcoded oligo-dT primers, featuring a 12-nt-long sample barcode (IDT). Double-stranded cDNA was generated by the second strand synthesis via the nick translation method. For that, a mix containing 2 µl of RNAse H (NEB, #M0297S), 1 µl of *E. coli* DNA ligase (NEB, #M0205 L), 5 µl of *E. coli* DNA Polymerase (NEB, #M0209 L), 1 µl of dNTP (10 mM), 10 µl of 5x Second Strand Buffer (100 mM Tris, pH 6.9, (AppliChem, #A3452), 25 mM $MgCl_2$ (Sigma, #M2670), 450 mM KCl (AppliChem, #A2939), 0.8 mM β-NAD (Sigma, N1511), 60 mM $(NH_4)_2SO_4$ (Fisher Scientific Acros, #AC20587), and 11 µl of water was added to 20 µl of ExoI-treated first-strand reaction on ice. The reaction was incubated at 16 °C for 2.5 h. Full-length double-stranded cDNA was purified with 30 µl (0.6x) of AMPure XP magnetic beads (Beckman Coulter, #A63881) and eluted in 20 µl of water.

The Illumina-compatible libraries were prepared by tagmentation of 10-40 ng of full-length double-stranded cDNA with 1 µl of in-house produced Tn5 enzyme (11 µM). After tagmentation, the libraries where purified with DNA Clean and Concentrator kit (Zymo Research #D4014), eluted in 20 µl of water and PCR amplified using 25 µl NEB Next High-Fidelity 2x PCR Master Mix (NEB, #M0541 L), 2.5 µl of each i5 and i7 Illumina index adapter (IDT) following this program: incubation 72 °C—3 min, denaturation 98 °C—30 s; 15 cycles: 98 °C—10 s, 63 °C—30 s, 72 °C—30 s; final elongation at 72 °C—5 min. The libraries were purified twice with AMPure beads (Beckman Coulter, #A63881) at 0.6x ratio to remove the fragments < 300 nt. The resulting libraries were profiled with a High Sensitivity NGS Fragment Analysis Kit (Advanced Analytical, #DNF-474) and measured with the Qubit dsDNA HS Assay Kit (Invitrogen, #Q32851) prior to pooling and sequencing using the Illumina NextSeq 500 platform using a custom primer and the High Output v2 kit (75 cycles) (Illumina, #FC-404-2005). The library loading concentration was 2.4 pM and the sequencing configuration as follows: R1 21c / index i7 8c / index i5 8 c/ R2 55c.

**Analysis of bulk RNA-seq**

Preprocessing

After sequencing and standard Illumina library demultiplexing, the *.fastq* files were aligned to the mouse reference genome mm10 (GRCm38 release 100 from Ensembl) using STAR (Version 2.7.3a), excluding multiple mapped reads. Resulting BAM files were sample-demultiplexed using BRB-seqTools v.1.4 (https://github.com/DeplanckeLab/BRB-seqTools) and the “gene expression x samples” read and UMI count matrices were generated using HTSeq v0.12.4.

General methods

Samples with too low read numbers or UMIs were filtered out. Genes with a count per million greater than 1 in at least 3 samples were retained. Raw counts were then normalized as log counts per million with a pseudo count of 1, using the function cpm() from *EdgeR* (McCarthy, Chen and Smyth, 2012) version 3.30.3. If the samples were from different batches, the raw counts were first normalized using quantile normalization as implemented in voom() from the package *limma* (Ritchie *et al.*, 2015) version 3.44.3 and then corrected for batch effects using combat() from *sva* (Johnson, Li and Rabinovic, 2007) version 3.36.0. PCAs were computed using prcomp() with the parameter center and scale set to TRUE. Differential expression analyses were computed using *DESeq2* (Love, Huber and Anders, 2014) version 1.28.1 and adding batch as a cofactor if necessary.

Scores

Scores were calculated as the sum of the normalized expression scaled between 0 and 1 per gene belonging to specific gene lists defined as follows:

- the “CD142+ score” was based on the top CD142+ markers (**Suppl. Table 1**), defined in “Selection of CD142+ top 100 markers” (**Fig. 1G**, **Fig. 2G**, **H**, **Fig. 4F**, **H**)
- the “White fat cell differentiation score” was based on the genes of the GO term GO:0050872 (**Fig. 1H**, **Fig. 4F**, **G**, **H**, **Supp. Fig. 23A, B, E**)
- the “Negative regulation of fat cell differentiation score” was based on the genes of the GO term GO:0045599 (**Fig. 1I**)
- the “Retinol metabolic process score” was based on the genes of the GO term GO:0042572 (**Supp. Fig. 13C**)
- the “Retinoic acid score” (“RAS”) was based on the genes of the GO terms “Cellular response to RA” (GO:0071300) and “RA receptor signaling pathway” (GO:0048384) (**Fig. 4H**)

Gene expression heatmaps:

Heatmaps of **Fig. 1C**, **Fig. 3C**, **D**, **I**, **Fig. 4 D**, **I**, **Suppl Fig. 4A**, **Suppl Fig. 20A**, **Suppl Fig. 25B** and **Suppl Fig. 26A** display the row-normalized gene expression and were generated using *pheatmap* version 1.0.12. The columns and rows were clustered using the method "ward.2D" of hclust() of the package *stats*.

Gene set enrichment analysis:

Gene set enrichment analysis was performed using the package *clusterprofiler* (Yu *et al.*, 2012) version 3.16.1.

- **Supp. Fig. 5B**, **Supp. Fig. 19C**, **D**, **Suppl. Fig 20A**: GSEA was performed using gseGO() function on the genes ranked by the Wald statistic computed by differential expression analysis of CD142+ *versus* CD142− ASPCs post-differentiation;
- **Supp. Fig. 12A**, **Supp. Fig. 13A**, **B**: GSEA was performed using gseGO() on the genes ranked by their loadings along the first principal component (PC1) of the PCA of freshly sorted total, CD142+ and CD142− ASPCs at different ages (**Fig. 2F**), showing that samples are ordered by age along PC1;
- **Supp. Fig. 14A**: GSEA was performed on the same ranking as **Supp Fig. 12A** described just above using GSEA() imputing the top CD142+ markers (**Suppl. Table 1**) as tested gene set;
- **Supp. Fig. 14D-G**: GSEA was performed on the genes ranked by the Wald statistic computed by the differential expression analysis of freshly sorted CD142+ *versus* CD142− ASPCs performed by age (P0, P16, 4wo and adult (7 and 11wo)). The top 100 CD142+ markers were given as input gene set to GSEA();
- **Fig. 3I**, **Supp. Fig. 19A**: GSEA was performed using the gseGO() function on the genes ranked by the Wald statistic computed by the differential expression analysis of freshly sorted CD142+ *versus* CD142− ASPCs;
- **Supp. Fig. 23C**, **F, G**: GSEA was performed on the genes ranked by their loadings along PC1 of the PCA display in **Supp. Fig. 23D** using GSEA() testing for the three gene sets: top CD142+ markers, GO:0071300 "Cellular response to RA" and GO:0048384 "RA receptor signaling pathway". Significant positive enrichment was also obtained when GSEA was performed on the Wald statistic computed by the differential expression analysis of CD142− ASPCs co-cultured with "active Aregs" (si*Aldh1a2*, si*Gdf10*, scr, WT) and CD142+ ASPCs post-differentiation *versus* CD142− ASPCs co-cultured with "dysfunctional Aregs" (si*F3*, si*Mgp*) or CD142− ASPCs alone post-differentiation. Significant negative enrichment was obtained when GSEA was performed on the genes ranked by the coefficient of the linear regression performed between the "white fat cell differentiation score" and gene expression of the same dataset (see "Extra analysis specific to CD142− ASPCs co-cultured with KD CD142+ ASPCs" for further details).

Additional analyses specific to age-related samples:

Bulk RNA-seq samples of freshly isolated total, CD142+ and CD142− ASPCs at different ages were first analyzed using conventional log CPM normalization as described above (**Fig. 2F**, **G**, **Suppl. Fig. 12**, **Supp. Fig. 13**, **Suppl. Fig. 14A**, **B**, **D-G**). In a second part, the analysis was performed on the data corrected for age. More precisely, the data were normalized using the *Seurat* (Stuart *et al.*, 2019) pipeline, and scaled for Age using the function ScaleData() (**Fig. 2H**, **Suppl. Fig. 14C**).

Extra analysis specific to CD142− ASPCs co-cultured with KD CD142+ ASPCs

In a first analysis, CD142− ASPCs that were exposed via a transwell assay to WT CD142+ ASPCs, to CD142+ ASPCs with respectively si*Aldh1a2*, si*Mgp*, si*F3*, si*Gdf10*, scr or to an empty well were analyzed as described above in the "General method" section (**Fig. 4C**, differential expression analysis result of **Fig. 4D**, **Suppl. Fig 23C**, **F**, **G, Suppl. Fig. 25A**, differential expression analysis result of **Suppl. Fig. 25B**). As described in the manuscript, the samples clustered in two groups: CD142− ASPCs co-cultured with "active Aregs" (WT CD142+ ASPCs or CD142+ ASPCs treated with si*Aldh1a*2, si*Gdf10* or scr) or CD142− ASPCs co-cultured with "dysfunctional Aregs" (CD142+ ASPCs treated with si*Mgp*, si*F3*, or empty transwell (null)). Differential expression analysis was performed between these two groups and used to order the genes on the heatmaps in **Fig. 4D**, **Supp. Fig. 25B**.

In a second part, CD142+ and CD142− ASPC samples post differentiation were integrated in the analysis. The heatmap of **Fig. 4E** displays the Euclidean distance calculated on the first five principal components of the PCA (**Suppl. Fig. 23D**). The linear regression between the "white fat cell differentiation score" and the expression of each gene (white fat cell differentiation score ~ expression of gene n) was calculated. The volcano plot in **Fig. 4G** shows the estimated slope (x axis) *versus* the -$\log_{10}$(p-value), where the p-value for each term tests the null hypothesis that the coefficient is equal to zero.

An overview of all the different transcriptomic datasets is provided in **Suppl. Table 2**.

**Comparison of various datasets (scRNA-seq, bulk RNA-seq, mass spectrometry)**

The top CD142+ markers (**Suppl. Table 1**) were compared across various datasets of freshly isolated CD142+ *versus* CD142− ASPCs. **Fig. 1D** displays the spearman correlation and the linear regression fitted to the logFC of the expression of these markers in freshly isolated CD142+ ASPCs *versus* CD142− ASPCs of the different datasets by pairs. For bulk RNA-seq and mass spectrometry data, the logFC was defined as the $\log_2$FC of freshly isolated CD142+ over CD142− ASPCs. For the scRNA-seq, the logFC was defined as the average logFC across the logFC of the *F3*+ cluster over the rest of the cells calculated in the different datasets used for the integration described in the section "Integration of adult murine subcutaneous adipose tissue scRNA-seq datasets".

**Statistical methods**

The paired Student's *t*-test was used to determine statistical differences between two groups, with the null hypothesis being that the two groups are equal. Multiple comparisons were taken into account by applying false discovery rate (FDR) correction. When specified, one-way ANOVA followed by Tukey honest significant difference (HSD) *post hoc* correction was applied, the null hypothesis being defined so that the difference of means was zero. (Adjusted) * P value < 0.05, ** P value < 0.01, *** P value < 0.001 were considered statistically significant.

**Data availability**

All raw and processed RNA-seq data have been uploaded in the Array Express (www.ebi.ac.uk/arrayexpress) with the accession numbers: E-MTAB-10179, E-MTAB-10180, E-MTAB-10181, E-MTAB-10182, E-MTAB-10183, E-MTAB-10184.

The mass spectrometry proteomics data have been deposited to the ProteomeXchange Consortium *via* the PRIDE partner repository with the dataset identifier PXD024064.

Microscopy images are available upon request.

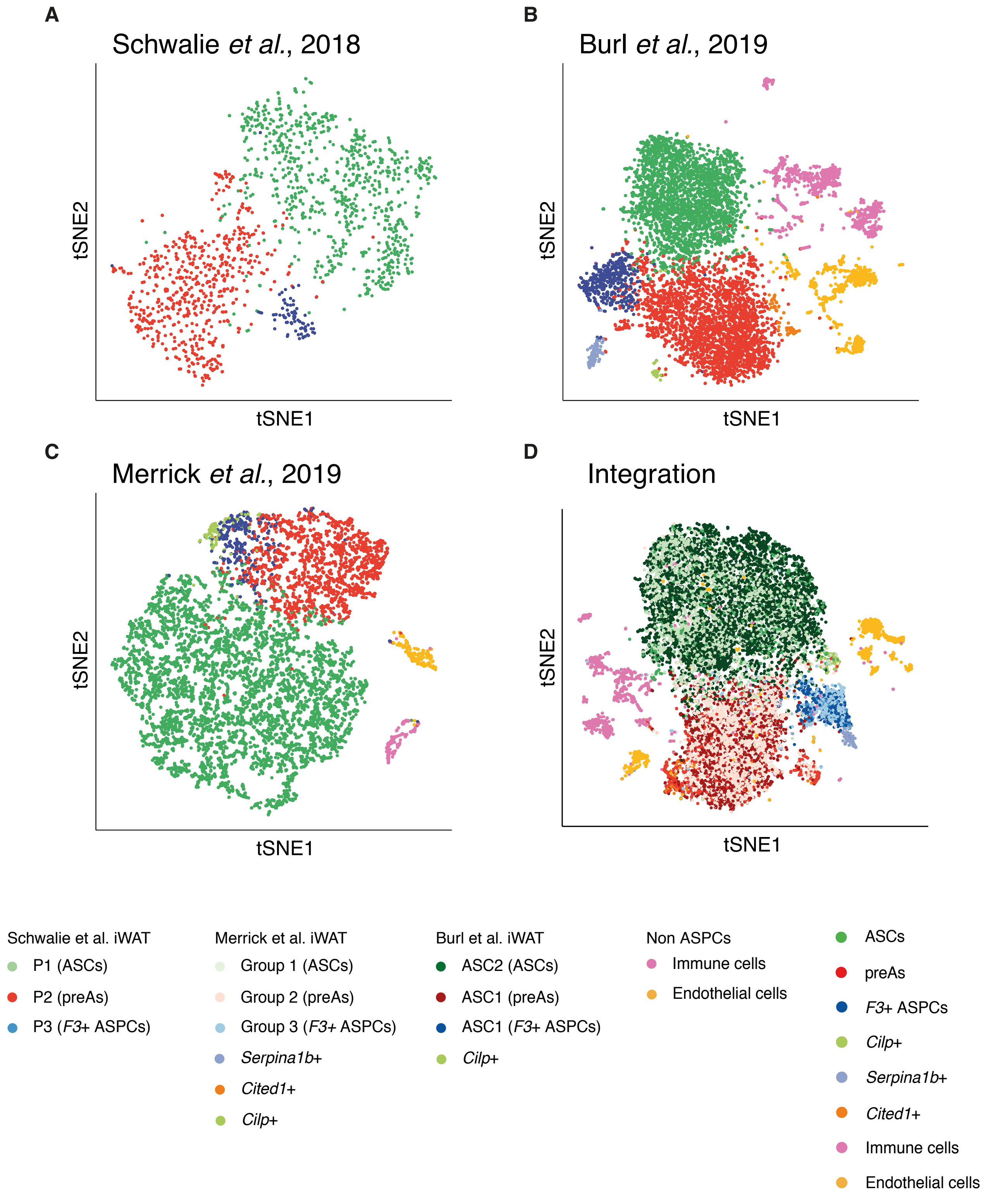


**Supplementary Figure 1. The *F3*+ cluster exists in distinct, publicly available single-cell RNA-seq datasets**

- **(A)** t-SNE cell map of the scRNA-seq dataset published in Schwalie *et al.*, 2018;
- **(B)** t-SNE cell map of the scRNA-seq dataset published in Burl *et al.*, 2019;
- **(C)** t-SNE cell map of the scRNA-seq dataset published in Merrick *et al.*, 2019;
- **(D)** t-SNE cell map of the integration analysis of the three datasets presented in **A-C**; the t-SNE is coloured by the clustering results identified in each dataset individually;

Three main ASPC subpopulations colour code: adipose stem cells (ASCs) — green, pre-adipocytes (PreAs) — red, *F3*(CD142)+ ASPCs — blue.

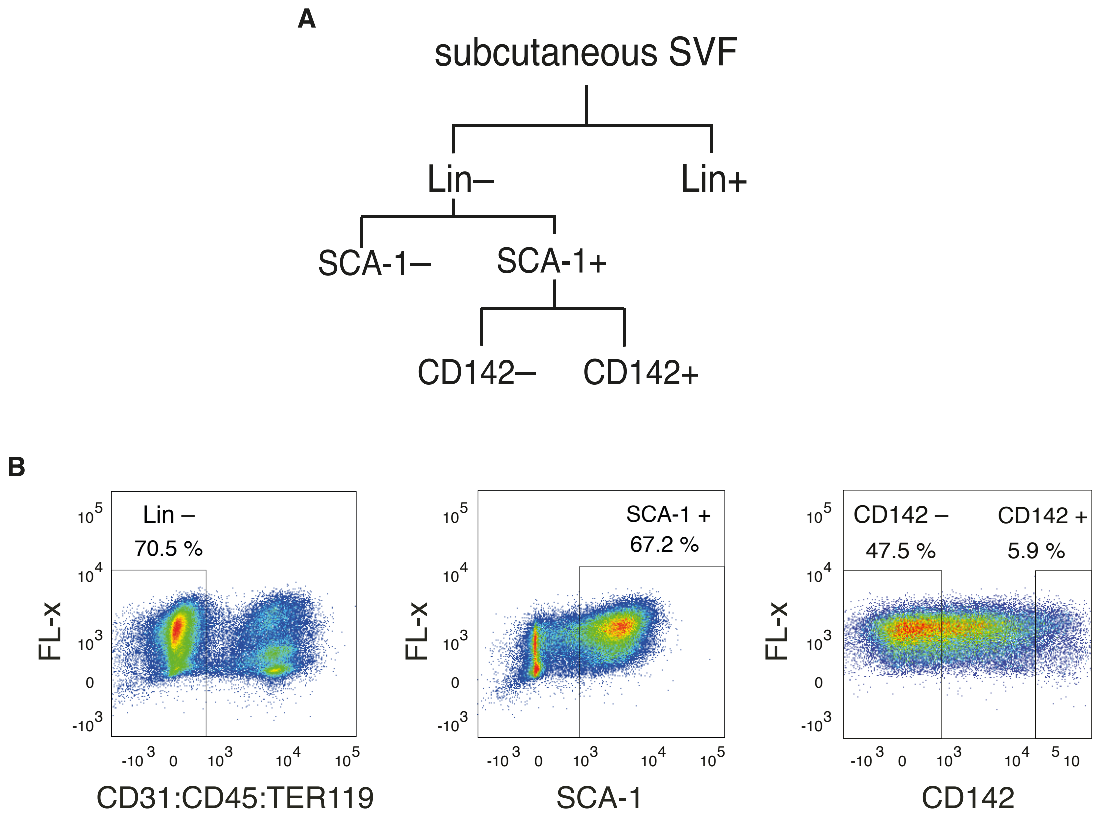


**Supplementary Figure 2. FACS-based gating strategy to isolate different ASPC fractions**

**(A)** Schematic of FACS-based selection of murine subcutaneous ASPCs;

**(B)** FACS-based gating strategy to isolate CD142+ and CD142– ASPCs defined as Lineage negative (Lin–: CD31– CD45– TER119–) SCA-1+ SVF cells (**Methods**).

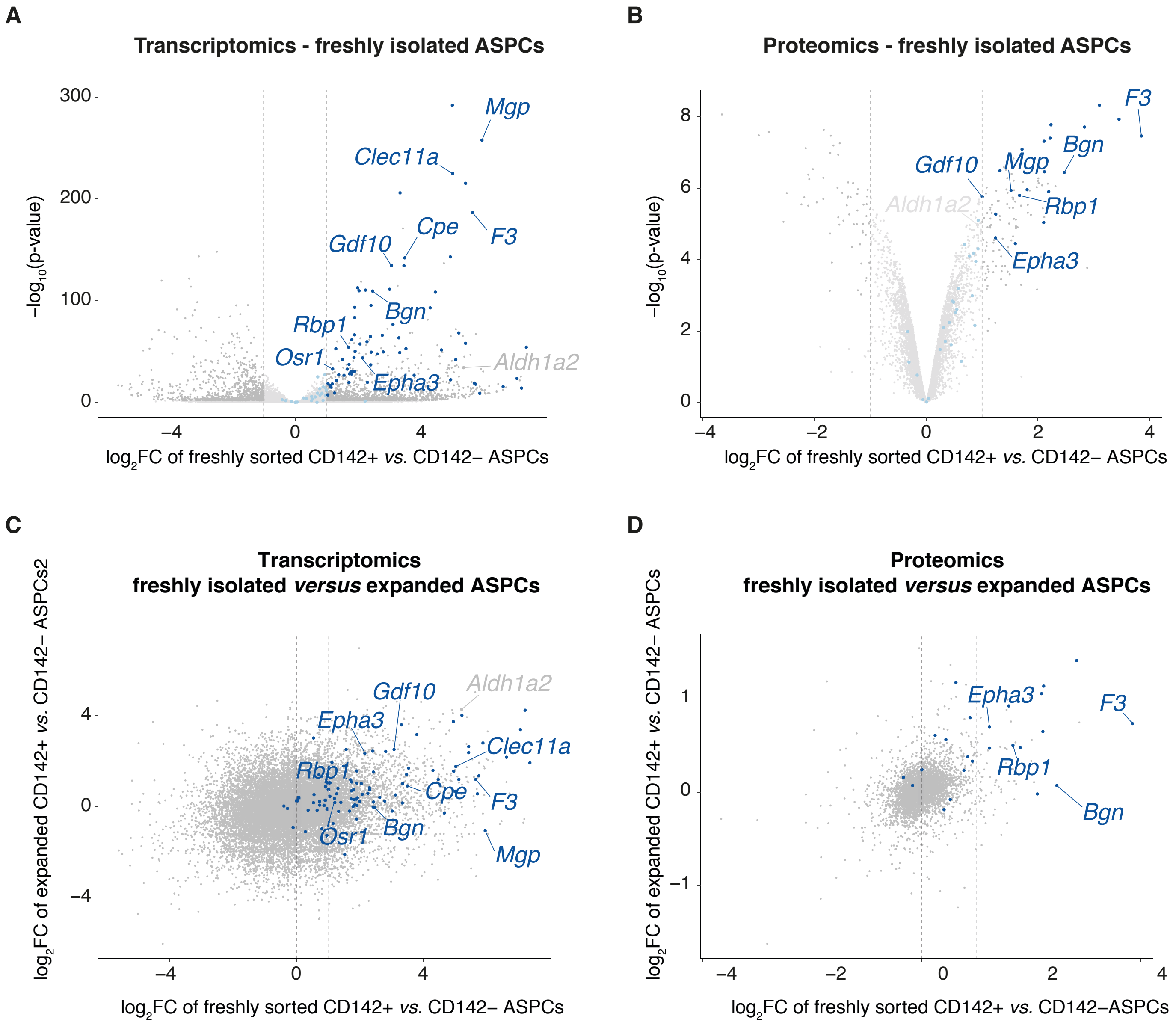


**Supplementary Figure 3. Top CD142+ markers are robust in freshly isolated CD142+ ASPCs across multiple omic datasets and some markers are maintained upon expansion**

- **(A)** Volcano plot displaying differential gene expression analysis based on bulk RNA-seq data of freshly isolated CD142+ *versus* CD142– ASPCs; top CD142+ markers (**Suppl. Table 1**) are highlighted in blue; significantly differentially expressed genes ($log_2FC > 1$, $FDR < 0.05$) are highlighted in darker colours;
- **(B)** Volcano plot displaying differential protein abundance analysis based on mass spectrometry data of freshly isolated CD142+ *versus* CD142– ASPCs; top CD142+ markers (**Suppl. Table 1**) are highlighted in blue; significantly differentially abundant proteins ($log_2FC > 1$, $FDR < 0.05$) are highlighted in darker colours;
- **(C)** Scatter plot showing the correlation of $log_2FC$ gene expression assessed by bulk RNA-seq of CD142+ over CD142– ASPCs in freshly isolated ASPCs (x axis) *versus* expanded ASPCs (y axis); top CD142+ markers (**Suppl. Table 1**) are highlighted in blue;
- **(D)** Scatter plot showing the correlation of $log_2FC$ protein abundance assessed by mass spectrometry of CD142+ over CD142– ASPCs in freshly isolated ASPCs (x axis) versus expanded ASPCs (y axis); top CD142+ markers (**Suppl. Table 1**) are highlighted in blue.

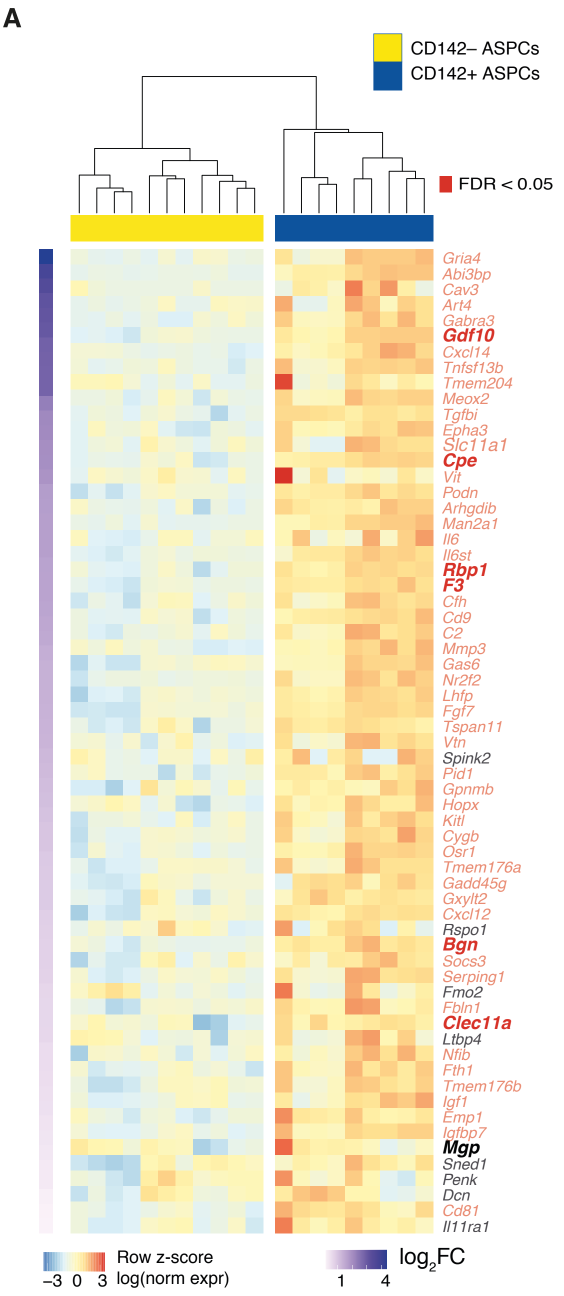

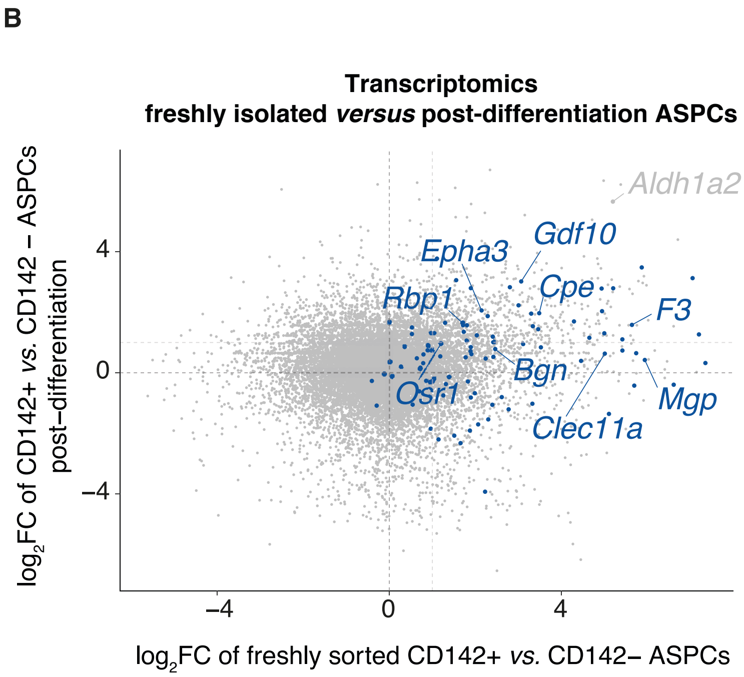


**Supplementary Figure 4. Many top CD142+ markers are maintained in CD142+ ASPCs upon exposure to an adipogenic cocktail**

**(A)** Gene expression heatmap across bulk RNA-seq samples of CD142+ and CD142– ASPCs post-differentiation (i.e. post exposure to an adipogenic cocktail); the selected genes are the top CD142+ markers (**Suppl. Table 1**) with a positive $\log_2$FC of CD142+ over CD142– ASPCs post-differentiation; the genes are ordered from biggest (top) to lowest (bottom) $\log_2$FC and significantly differentially expressed genes (FDR < 0.05) are highlighted in red; log normalized expression is scaled by row;

**(B)** Scatter plot showing the correlation of $\log_2$FC gene expression based on bulk RNA-seq of CD142+ over CD142– ASPCs in freshly isolated (x axis) *versus* post-differentiation (y axis) ASPCs; top CD142+ markers (**Suppl. Table 1**) are highlighted in blue.

**A**

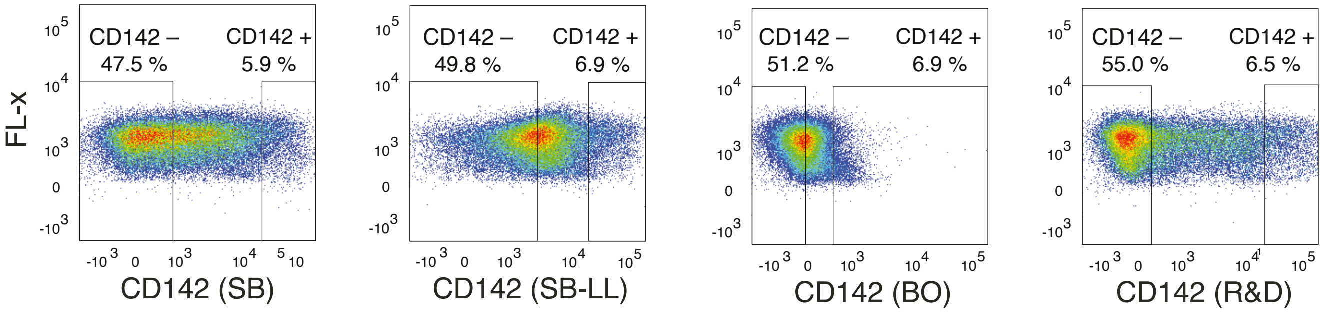


**B**

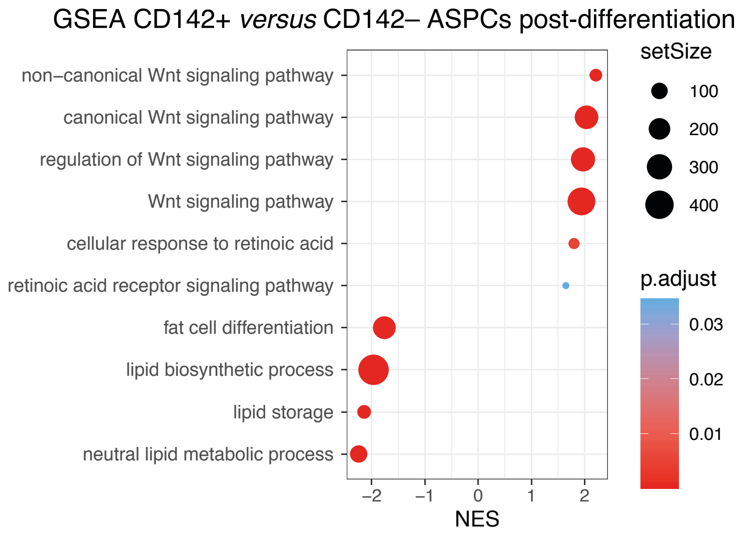


**Supplementary Figure 5. FACS-based gating strategy to isolate CD142+ ASPCs using different antibodies and GSEA of CD142+ *versus* CD142– ASPCs post-differentiation**

- **(A)** FACS-based gating strategy to isolate CD142+ ASPCs defined as 5-7% CD142+ ASPCs, and CD142– ASPCs defined as around 50% cells negative for CD142 marker, with the use of four different antibodies: “SB” – SinoBiological anti-CD142–PE, “SB-LL” SinoBiological anti-CD142 coupled to PE in house (**Methods**), “BO” – BiOrbyt anti-CD142–PE and “R&D” – R&D Systems anti-CD142–PE; the experiment was repeated four times with similar results, shown here is one representative biological replicate;
- **(B)** GSEA results performed on differential gene expression between CD142+ *versus* CD142- ASPCs post-differentiation; NES: normalized enrichment score.

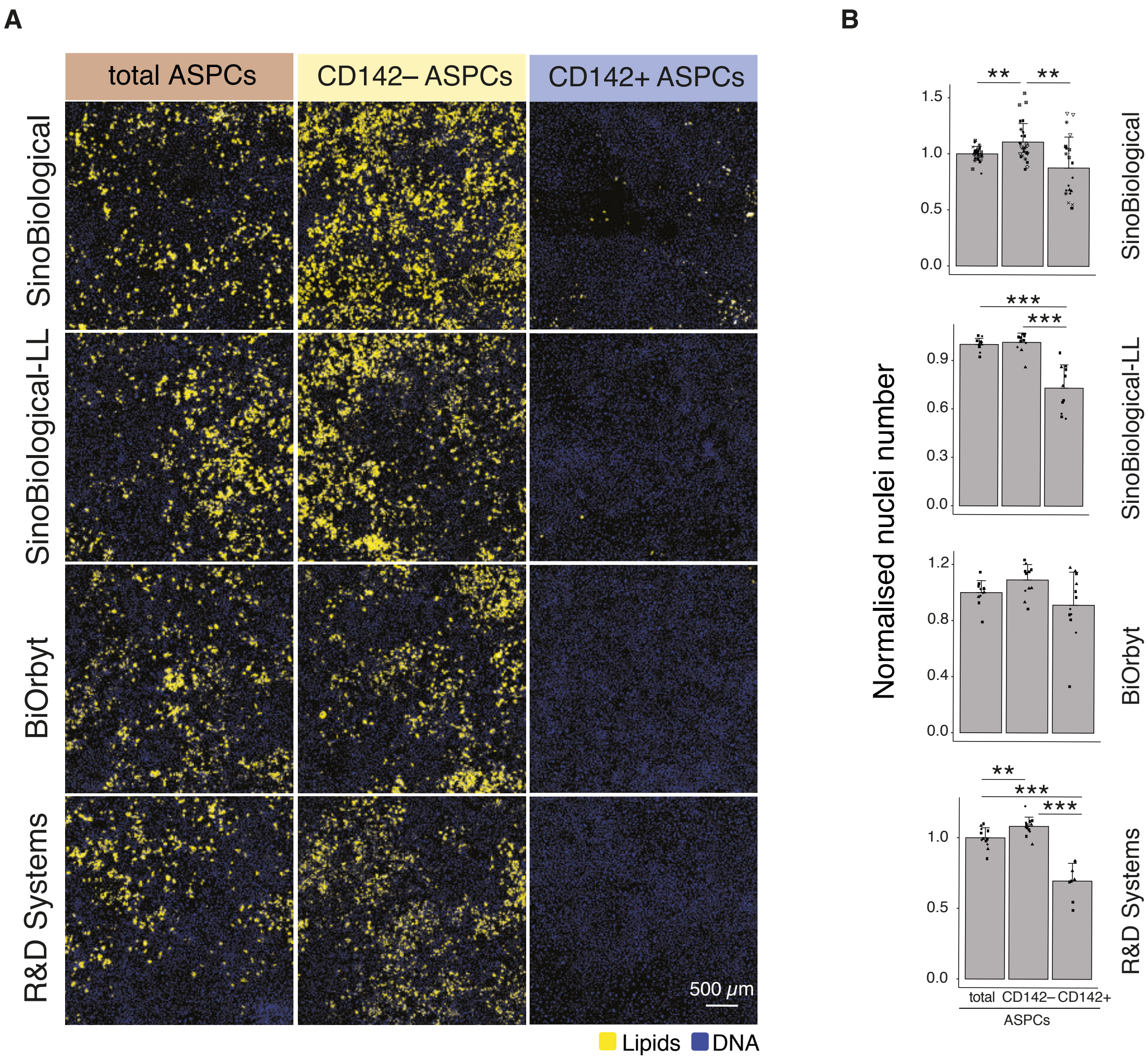


**Supplementary Figure 6. CD142+ ASPCs are non-adipogenic regardless of the antibody used for isolation**

**(A)** Representative fluorescence microscopy images of total, CD142– and CD142+ ASPCs isolated with the use of the anti-CD142 antibodies described in **Suppl. Fig. 5A**, after *in vitro* adipogenic differentiation; the presented images correspond to 25 tiled and thresholded 20x images containing 7-8 z-stacks in order to capture most of the well surface (**Methods**);

**(B)** Bar plots showing the nuclei numbers for the cellular fractions shown in **A**, n=9-15, 3-4 biological replicates, 3-5 independent wells for each;

In all images, nuclei are stained with Hoechst (blue) and lipids are stained with Bodipy (yellow); scale bars, 500 µm; *P ≤ 0.05, **P ≤ 0.01, ***P ≤ 0.001, pairwise two-sided *t*-test, for statistical details see **Methods**.

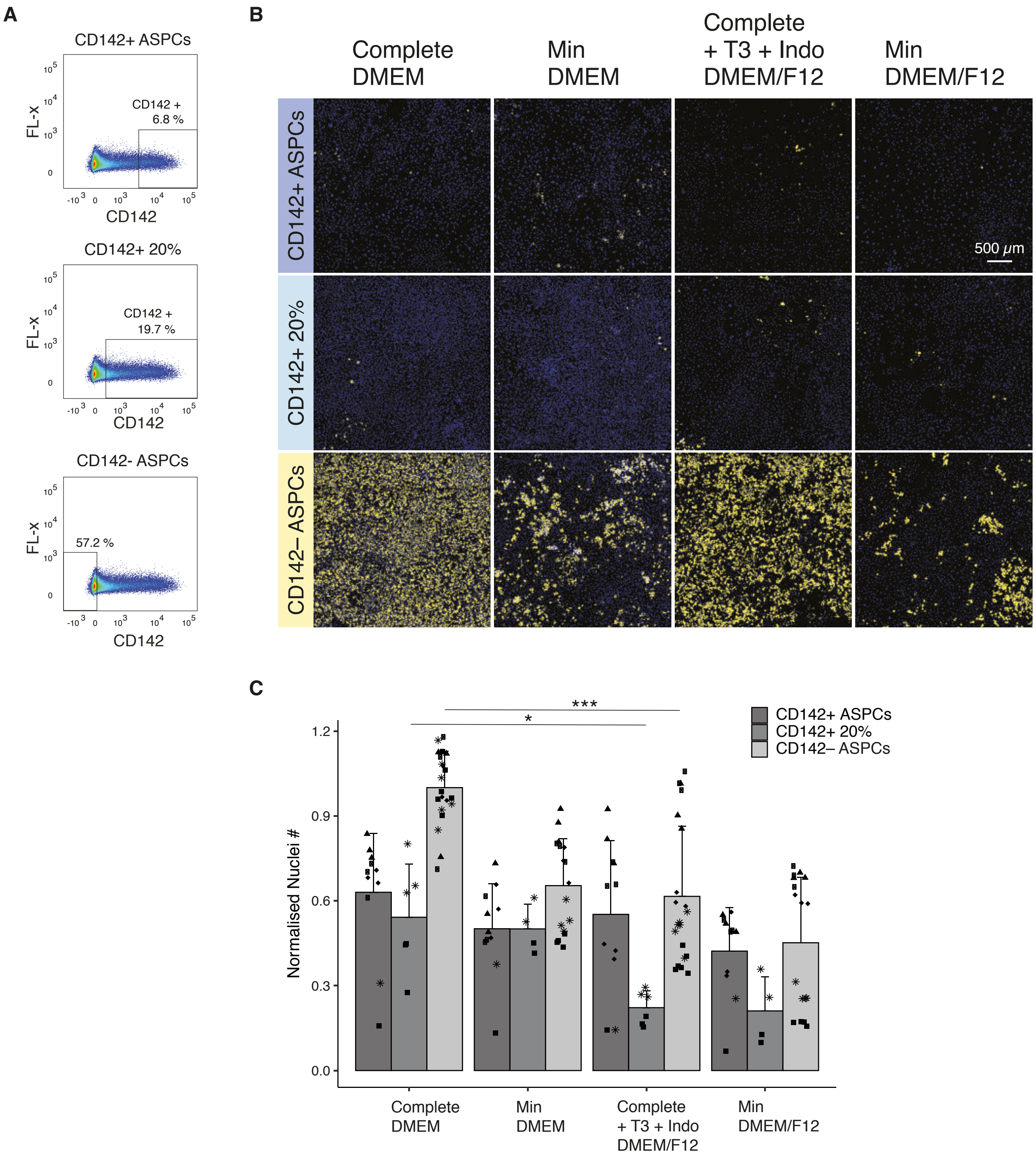


**Supplementary Figure 7. CD142+ ASPCs are non-adipogenic regardless of the stringency of the gating strategy**

- **(A)** FACS-based gating strategy to isolate CD142+ (5-7%), 20% CD142+ and CD142– ASPCs; the experiment was repeated at least five times with similar results, shown here is one representative biological replicate;
- **(B)** Representative fluorescence microscopy images of CD142+, 20% CD142+ and CD142– ASPCs isolated following the sorting strategy shown in **A**, after *in vitro* adipogenic differentiation with the indicated adipogenic differentiation cocktails (**Methods**); the presented images correspond to 25 tiled and thresholded 20x images containing 7-8 z-stacks in order to capture most of the well surface (**Methods**);
- **(C)** Bar plots showing the nuclei numbers for the cellular fractions shown in **B**, marker shapes correspond to different biological replicates, n=8-17, 3-5 biological replicates, 2-5 independent wells for each;

In all images, nuclei are stained with Hoechst (blue) and lipids are stained with Bodipy (yellow); scale bars, 500 μm; $*P \leq 0.05$, $**P \leq 0.01$, $***P \leq 0.001$, pairwise two-sided *t*-test, for statistical details see **Methods**.

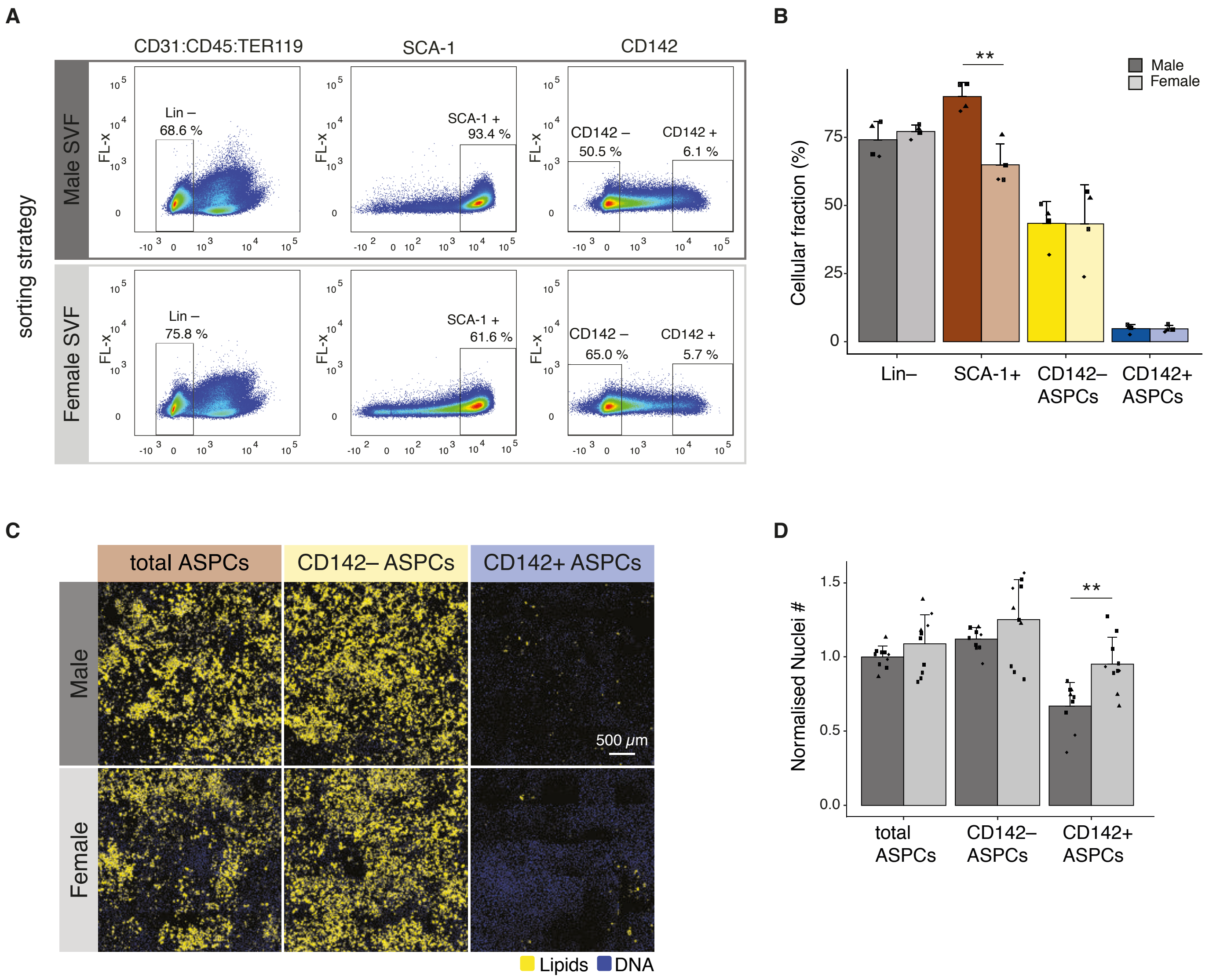


**Supplementary Figure 8. CD142+ ASPCs are non-adipogenic in male and female mice**

- **(A)** FACS-based gating strategy of male and female Lin–, SCA-1+ (ASPC), CD142– and CD142+ ASPC cellular fractions within the SVF; the experiment was repeated four times with similar results, shown here is one representative biological replicate;
- **(B)** Bar plots showing the percentage of the cellular fractions gated as shown in **A**, the experiment was repeated n=4;
- **(C)** Representative fluorescence microscopy images of male- and female-derived ASPCs, CD142– and CD142+ ASPCs, isolated following the sorting strategy shown in **A** after *in vitro* adipogenic differentiation; the presented images correspond to 25 tiled and thresholded 20x images containing 7-8 z-stacks in order to capture most of the well surface (**Methods**): n=9-10, 4 biological replicates, 2-3 independent wells for each;
- **(D)** Bar plots showing the nuclei numbers for the cellular fractions shown in **C**, marker shapes correspond to different biological replicates, n=9-10, 4 biological replicates, 2-3 independent wells for each;

In all images, nuclei are stained with Hoechst (blue) and lipids are stained with Bodipy (yellow); scale bars, 500 μm; bar colours: total ASPCs - brown, CD142– ASPCs — yellow, CD142+ ASPCs — blue, female — lighter colours, male — darker colours; $*P \leq 0.05$, $**P \leq 0.01$, $***P \leq 0.001$, pairwise two-sided *t*-test, for statistical details see **Methods**.

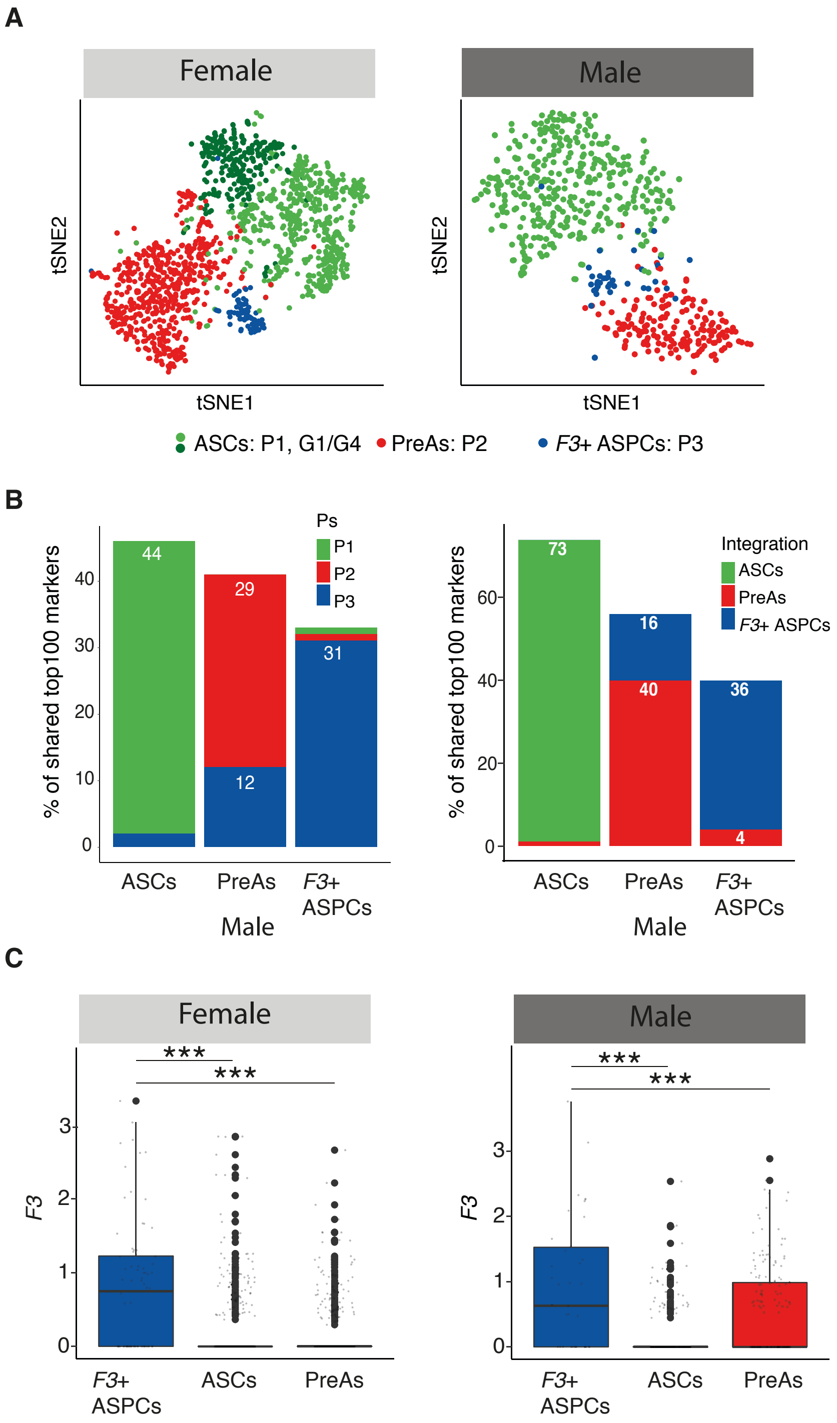


**Supplementary Figure 9. scRNA-seq validates the molecular existence of *F3*(CD142)+ ASPCs in male and female mice**

**(A)** t-SNE cell map of an scRNA-seq dataset of female (**left**) or male (**right**) murine subcutaneous ASPCs: adipose stem cells (ASCs) — green, pre-adipocytes (PreAs) — red, *F3*(CD142)+ ASPCs — blue;

**(B)** Percentage of the top 100 markers (**left**) of the three clusters (P1 (equivalent to ASCs) in green, P2 (equivalent to PreAs) in red and P3 (equivalent to *F3*(CD142)+ ASPCs) in blue) published in Schwalie *et al.*, 2018 (analysis performed on female ASPCs, t-SNE shown in panel **A left**), or (**right**) of the top 100 markers of the three main ASPC populations based on the integration of adult (male and female) ASPCs (**Fig. 1A**, Ferrero *et al.*, 2020) that overlap with the top markers of the same three subpopulations identified in male ASPCs (t-SNE shown in panel **A right**);

**(C)** Boxplots showing the distribution of log normalized expression of *F3* in female (**left**) and male (**right**) ASPCs across the three main ASPC subpopulations;

Adipose stem cells (ASCs) — green, pre-adipocytes (PreAs) — red, *F3*(CD142)+ ASPCs — blue; $^{*}P \leq 0.05$, $^{**}P \leq 0.01$, $^{***}P \leq 0.001$, ANOVA and Tukey HSD *post hoc* test, for statistical details see **Methods**.

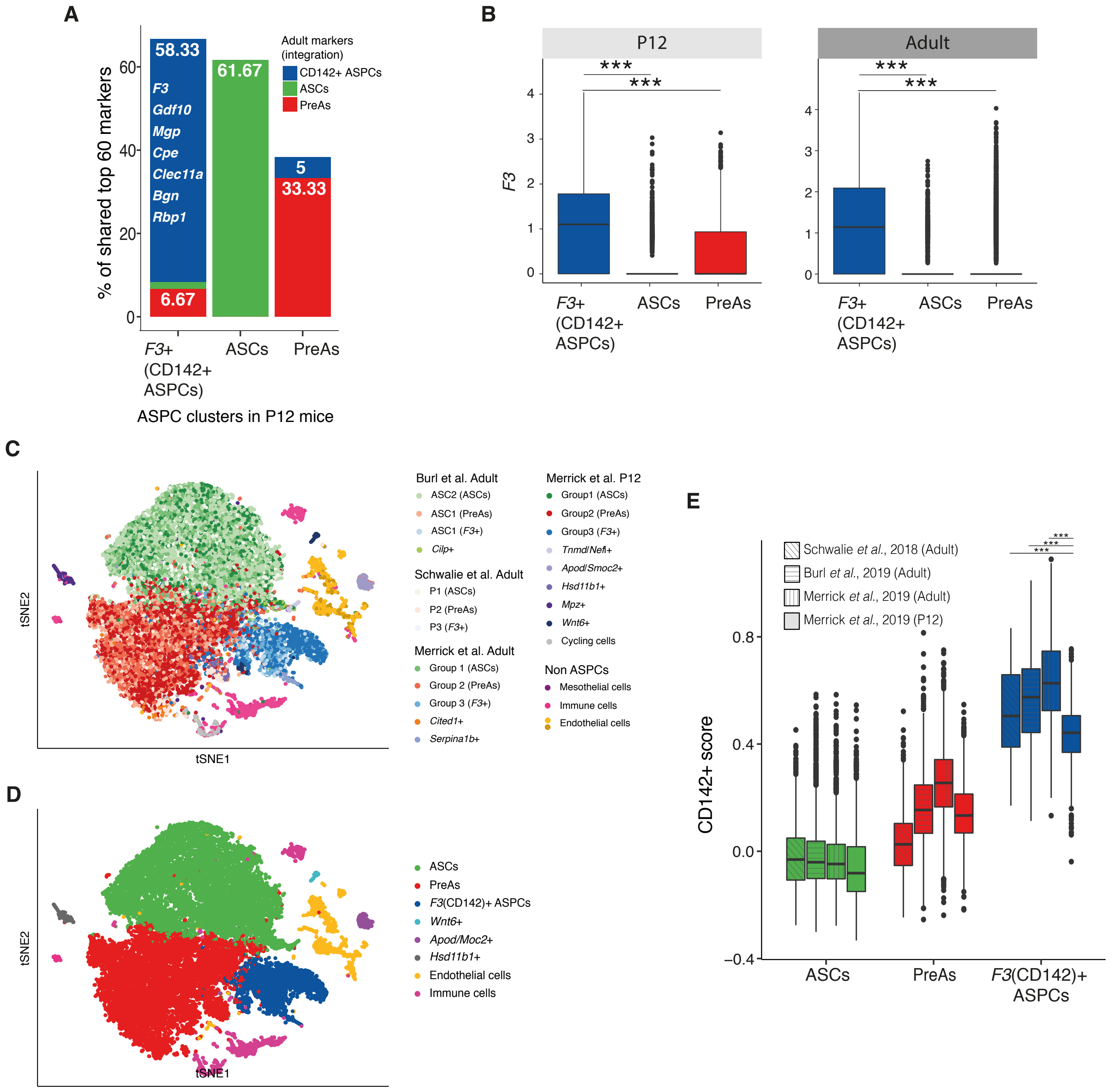


**Supplementary Figure 10. The CD142+ score of *F3*(CD142)+ ASPCs is less pronounced in P12 mice**

- **(A)** Percentage of the top 60 markers (**Methods**) of the three main ASPC subpopulations (adipose stem cells (ASCs) in green, pre-adipocytes (PreAs) in red, *F3*(CD142)+ ASPCs in blue) based on the integration of adult ASPCs (**Fig. 1A**, Ferrero *et al.*, 2020) that overlap with the top 60 markers of the same three subpopulations identified in P12 ASPCs;
- **(B)** Boxplots showing the distribution of log normalized expression of *F3* in P12 (**left**) and adult (**right**) ASPCs across the main ASPC subpopulations;
- **(C)** t-SNE cell map of the integration analysis of adult- (Schwalie *et al.*, 2018, Burl *et al.*, 2019, Merrick *et al.*, 2019) and P12- (Merrick *et al.*, 2019) derived ASPCs coloured by the clustering results identified in each dataset individually;
- **(D)** t-SNE cell map of the integration analysis described in **C** coloured by the clustering results, showing the main three ASPCs subpopulations as well as other subgroups of ASPCs, endothelial and immune cells;
- **(E)** Boxplot showing the distribution of “CD142+ score” (**Suppl. Table 1**) in the four different datasets used in the integration described in **C** across the three main ASPC subpopulations;

*F3*(CD142)+ ASPCs — blue, adipose stem cells (ASCs) — green, pre-adipocytes (PreAs) — red; *P ≤ 0.05, **P ≤ 0.01, ≤ 0.001, one-way ANOVA and Tukey HSD *post hoc* test (**B**, **E** null hypothesis: no difference between means of *F3*+ ASPCs in each dataset), for statistical details see **Methods**.

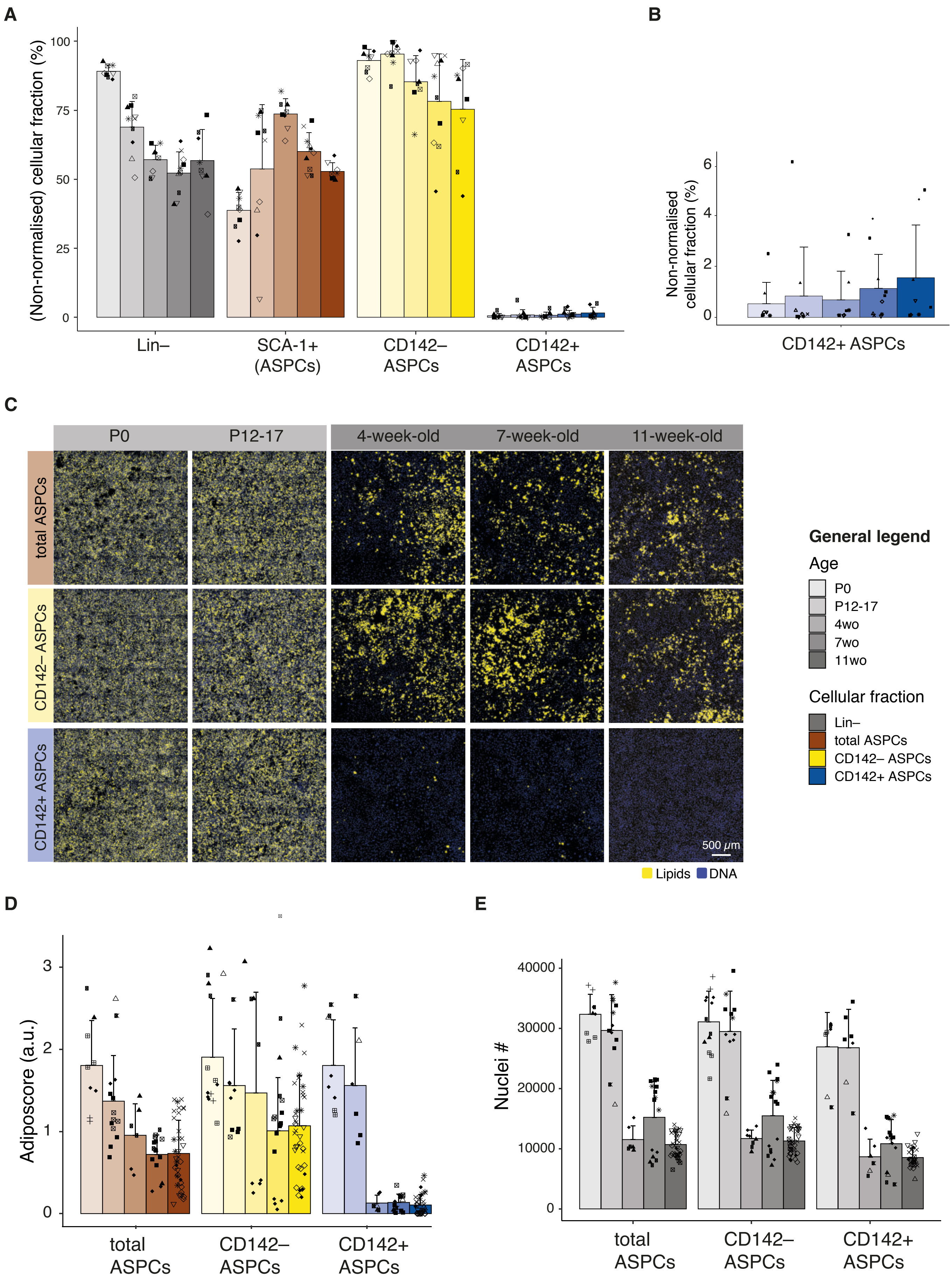


**Supplementary Figure 11** | See next page for caption

**Supplementary Figure 11. The non-adipogenic character of CD142+ ASPCs is age-dependent**

- **(A)** Bar plot showing the non-normalised parental percentage of cellular fractions gated for FACS-based isolation of the indicated subpopulations at different ages: P0, P12-17, 4wo, 7wo and 11wo, marker shapes correspond to different biological replicates, n=8-10;
- **(B)** Bar plot showing the non-normalised cellular fractions gated for FACS-based isolation of P0-, P12-17-, 4wo-, 7wo- and 11wo-derived CD142+ ASPCs, marker shapes correspond to different biological replicates, n=8-10;
- **(C)** Representative fluorescence microscopy images of P0-, P12-17-, 4wo-, 7wo- and 11wo-derived total, CD142– and CD142+ APSCs after in vitro adipogenic differentiation; the presented images correspond to 25 tiled and thresholded 20x images containing 7-8 z-stacks in order to capture most of the well surface (**Methods**);
- **(D)** Fraction of differentiated cells per ASPC type shown in **C**; bar colour shading corresponds to the individual ages as indicated, marker shapes correspond to different biological replicates, n=6-35, 3-13 biological replicates, 2-6 independent wells for each;
- **(E)** Bar plot showing the nuclei numbers for the cellular fractions shown in **C**; marker shapes correspond to different biological replicates, n=6-35, 3-13 biological replicates, 2-6 independent wells for each;

In all images, nuclei are stained with Hoechst (blue) and lipids are stained with Bodipy (yellow); scale bars, 500 µm, bar colours: total ASPCs - brown, CD142– ASPCs - yellow, CD142+ ASPCs - blue; shades are indicative of age (young - light, adult - dark); $^{*}P \leq 0.05$, $^{**}P \leq 0.01$, $^{***}P \leq 0.001$, pairwise two-sided *t*-test, for statistical details see **Methods**.

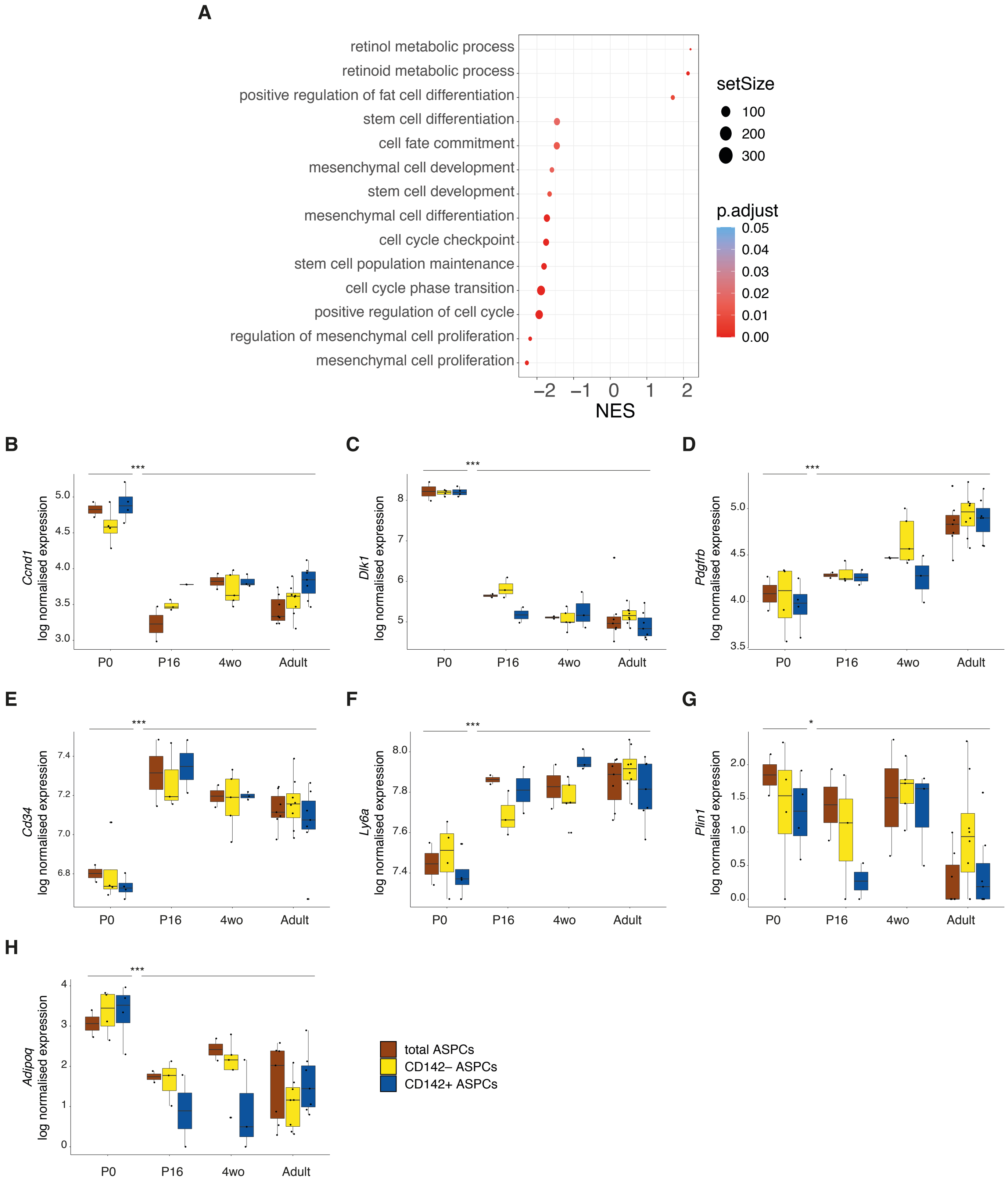


**Supplementary Figure 12. Gene expression profiling of P0 ASPCs suggests that these cells are in a "naïve" state**

**(A)** Significantly enriched terms found by GSEA performed on the genes driving the first principal component of the PCA representation of bulk RNA-seq samples of P0, P16-, 4-, 7- and 11wo-derived total, CD142– and CD142+ ASPCs (**Fig. 2F**); negative and positive NES (Normalized enrichment score) indicates enrichment in young or adult samples respectively;

**(B-H)** Boxplots showing the distribution of the log normalized expression of (**B**) *Ccnd1* (coding for Cyclin D1), (**C**) *Dlk1* (coding for PREF-1), (**D**) *Pdgfrb*, (**E**) *Cd34*, (**F**) *Ly6a* (coding for SCA-1), (**G**) *Plin1* (coding for Perilipin), (**H**) *Adipoq* (coding for Adiponectin) in total ASPCs, CD142– and CD142+ ASPCs at the indicated ages;

*P ≤ 0.05, **P ≤ 0.01, ***P ≤ 0.001, pairwise two-sided *t*-test, for statistical details see **Methods**.

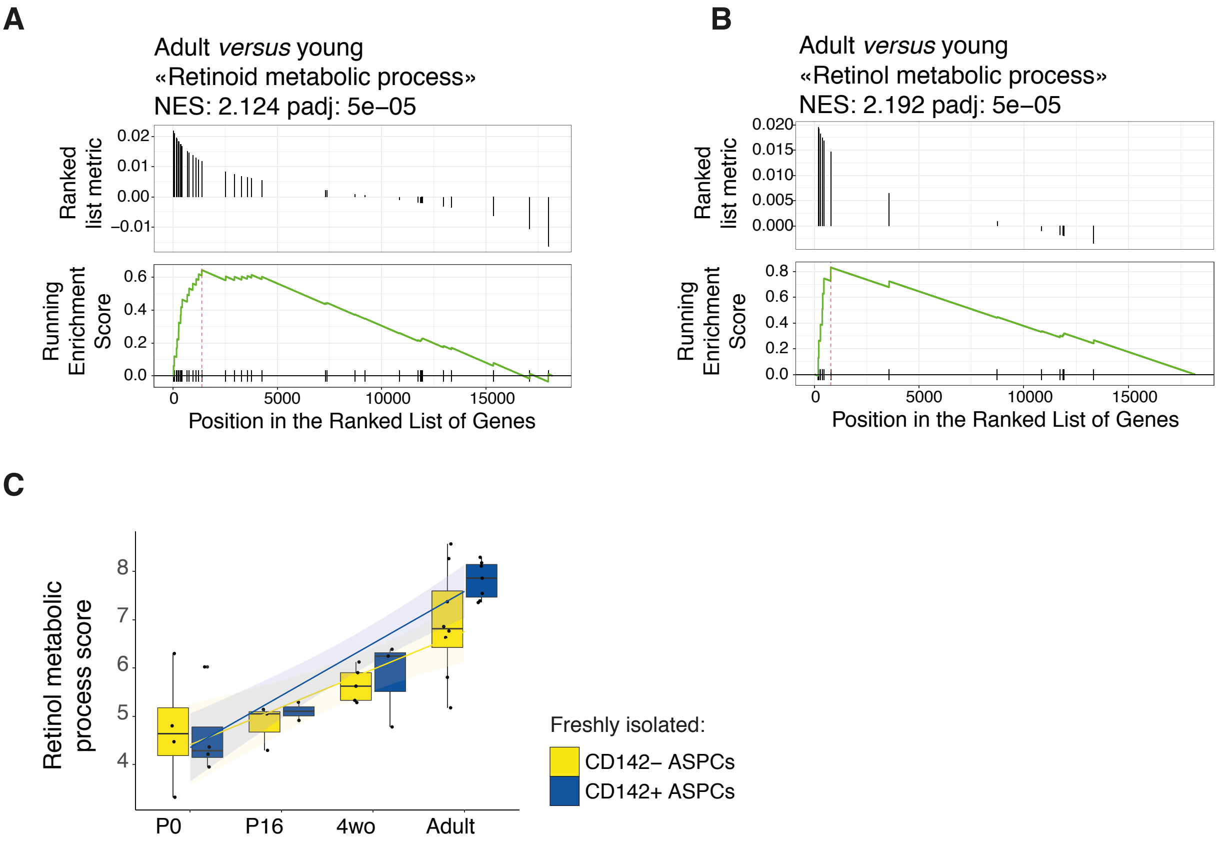


**Supplementary Figure 13. Genes linked to retinol/retinoid metabolic processes are enriched in adult compared to young ASPCs**

- **(A)** The term “retinoid metabolic process” (GO:0001523) was identified as significantly enriched in freshly isolated adult- compared to young-derived ASPCs when performed on the genes driving the PC1 of the PCA representation based on bulk RNA-seq samples of P0-, P16-, 4-, 7- and 11wo-derived ASPCs (see **Fig. 2F**);
- **(B)** The term “retinol metabolic process” (GO:0042572) was identified as significantly enriched in adult- compared to young mice-derived ASPCs when performed on the genes as described in **A**;
- **(C)** Boxplot showing the distribution of the “retinol metabolic process score” based on the expression of the genes linked to “retinol metabolic process” (GO:0001523) for CD142– and CD142+ ASPCs at the indicated ages.

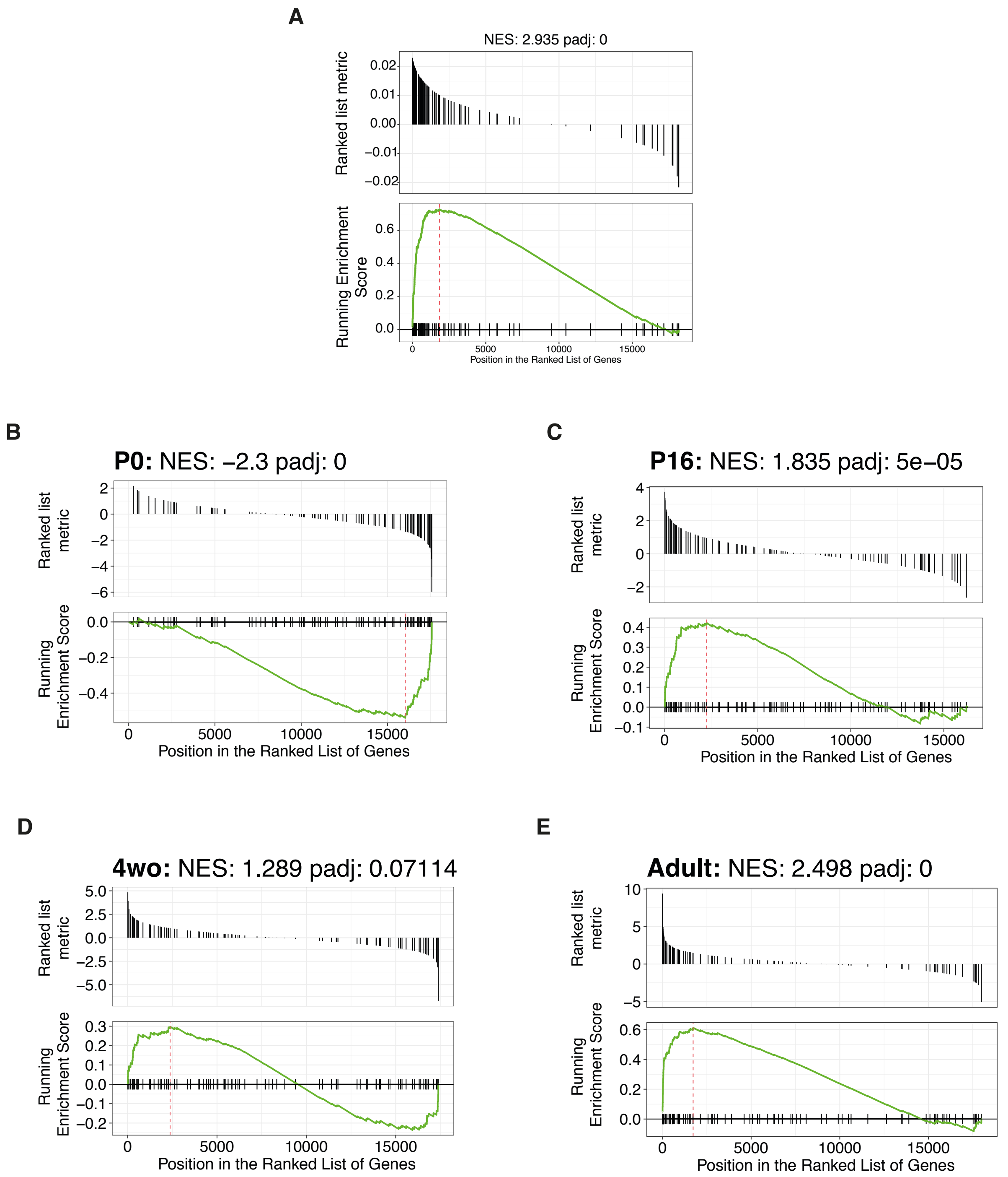


**Supplementary Figure 14. The molecular "CD142+ signature" appears in CD142+ ASPCs in an age-dependent manner**

**(A)** GSEA of the top CD142+ markers (**Suppl. Table 1**) performed on the genes driving PC1 of the PCA representation of bulk RNA-seq samples of P0-, P16-, 4-, 7- and 11wo-derived freshly isolated total, CD142– and CD142+ ASPCs (see **Fig. 2F**);

**(B-E)** GSEA of the top CD142+ markers (**Suppl. Table 1**) performed on the differential expression data between CD142+ and CD142– ASPCs of P0 (**D**), P16 (**E**), 4wo (**F**) and adults (7 and 11wo) (**G**).

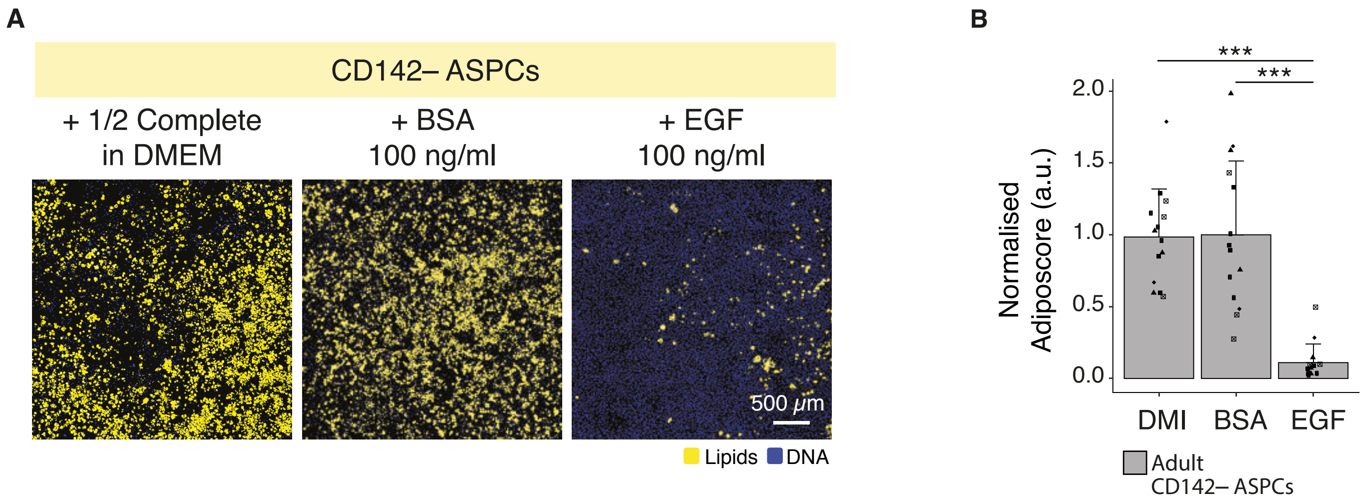


**Supplementary Figure 15. Positive and negative controls of chemical inhibition of adipogenesis**

- **(A)** Representative fluorescence microscopy images of CD142– ASPCs after *in vitro* adipogenic differentiation with the differentiation cocktail (1/2 Complete in DMEM, **Methods**) supplemented with control recombinant proteins BSA (negative control) and EGF (positive control) of inhibition of adipogenesis at 100 ng/ml; the presented images correspond to 25 tiled and thresholded 20x images containing 7-8 z-stacks in order to capture most of the well surface (**Methods**);
- **(B)** Fraction of differentiated CD142– ASPCs treated with the indicated recombinant proteins shown in **A**, marker shapes correspond to different biological replicates, n=12, 4 biological replicates, 3 independent wells for each;

In all images, nuclei are stained with Hoechst (blue) and lipids are stained with Bodipy (yellow); scale bars, 500 μm; $*P \leq 0.05$, $**P \leq 0.01$, $***P \leq 0.001$, pairwise two-sided *t*-test, for statistical details see **Methods**.

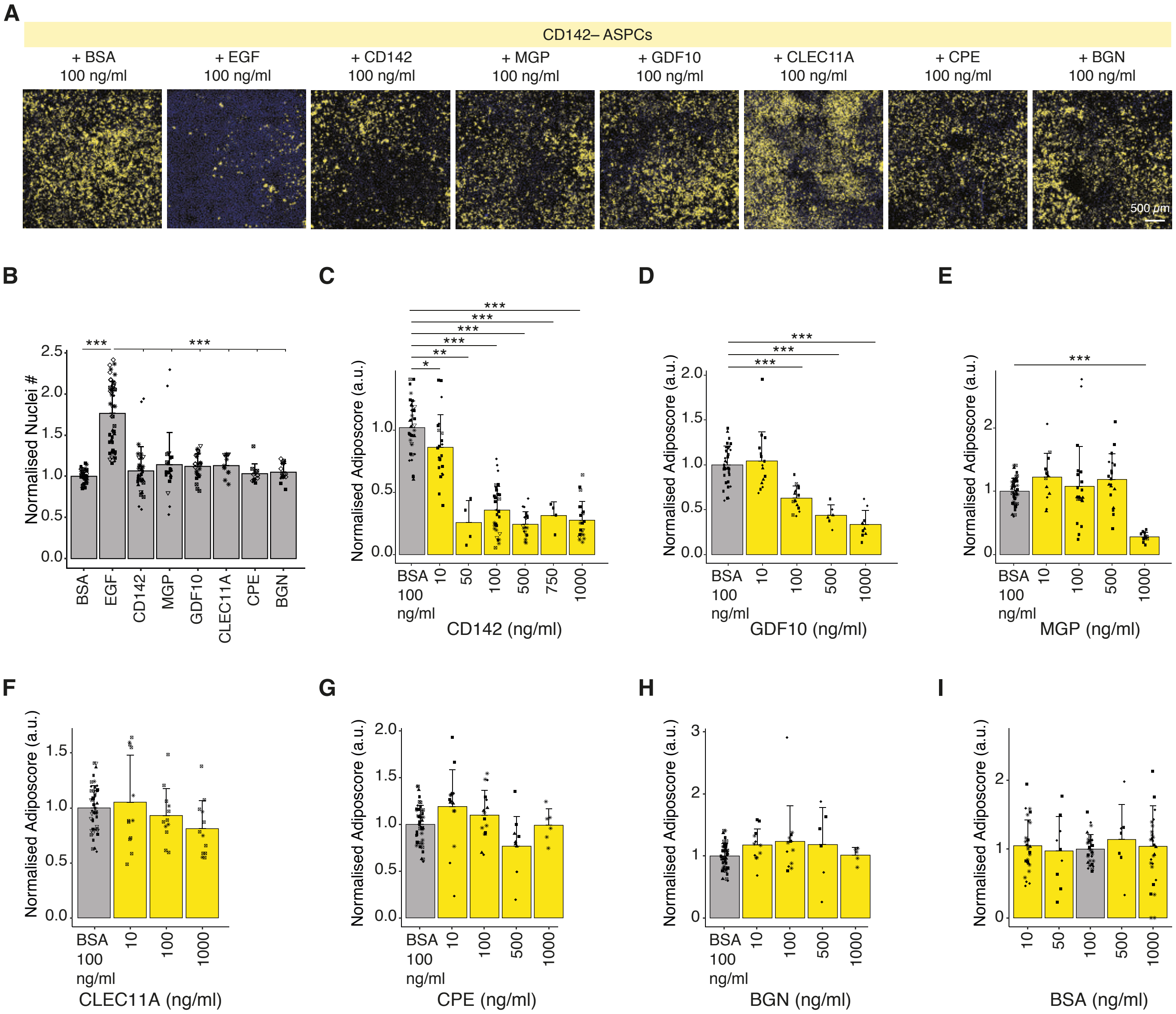


**Supplementary Figure 16. Adipogenic inhibition of adult-derived CD142– ASPCs using recombinant proteins**

**(A)** Representative fluorescence microscopy images of adult-derived CD142– ASPCs after *in vitro* adipogenic differentiation; the induction cocktail was supplemented with recombinant proteins corresponding to controls and to the selected CD142+ ASPC (Areg)-specific candidates: BSA (negative control), EGF (positive control), CD142, MGP, GDF10, CLEC11A, CPE and BGN at 100 ng/ml; the presented images correspond to 25 tiled and thresholded 20x images containing 7-8 z-stacks in order to capture most of the well surface (**Methods**);

**(B)** Bar plots showing the nuclei numbers for cellular fractions shown in **A**, marker shapes correspond to different biological replicates, n=11-59, 2-10 biological replicates, 3-9 independent wells for each;

**(C-I)** Bar plots showing the fraction of differentiated adult-derived CD142– ASPCs treated with the indicated recombinant proteins at the indicated concentrations, marker shapes correspond to different biological replicates;

In all images, nuclei are stained with Hoechst (blue) and lipids are stained with Bodipy (yellow); scale bars, 500 μm, bar colours: CD142– ASPCs treated with the indicated recombinant protein of interest — yellow, nuclei (**B**) or recombinant BSA treatment (negative control) (**C-I**) — light grey; *P ≤ 0.05, **P ≤ 0.01, ***P ≤ 0.001, pairwise two-sided *t*-test, for statistical details see **Methods**.

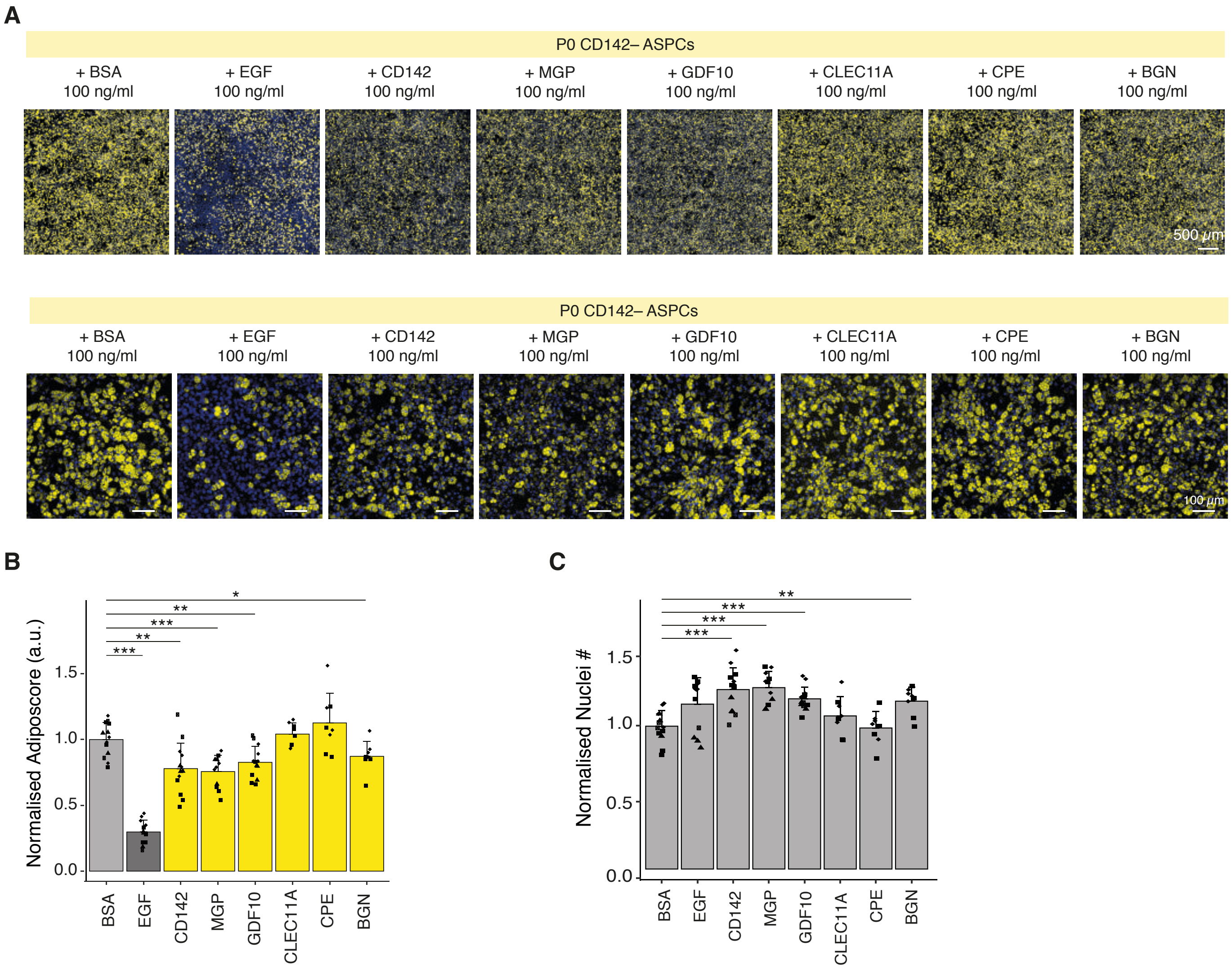


**Supplementary Figure 17. Adipogenic inhibition of P0-derived CD142– ASPCs with recombinant proteins**

- **(A)** Representative fluorescence microscopy images of P0-derived CD142– ASPCs after *in vitro* adipogenic differentiation; the induction cocktail was supplemented with recombinant proteins corresponding to controls and to the selected CD142+ ASPC (Areg)-specific candidates: BSA (negative control), EGF (positive control), CD142, MGP, GDF10, CLEC11A, CPE and BGN at 100 ng/ml; the presented images correspond to 25 tiled and thresholded 20x images containing 7-8 z-stacks in order to capture most of the well surface (**Methods**); scale bars 500 µm (**top**), 100 µm (**bottom**);
- **(B)** Fraction of differentiated CD142– ASPC cells treated with the indicated recombinant proteins shown in **A**; marker shapes correspond to different biological replicates, n=8-13, 1-3 biological replicates, 2-4 independent wells for each;
- **(C)** Bar plots showing the nuclei numbers for the cellular fractions shown in **A**; marker shapes correspond to different biological replicates, n=11-59, 2-10 biological replicates, 3-9 independent wells for each;

In all images, nuclei are stained with Hoechst (blue) and lipids are stained with Bodipy (yellow); bar colours: CD142– ASPCs treated with the indicated recombinant protein of interest — yellow, recombinant BSA treatment (negative control) (**B**) or nuclei (**C**) — light grey, EGF treatment (positive control) (**B**) — dark grey; *P ≤ 0.05, **P ≤ 0.01, ***P ≤ 0.001, pairwise two-sided *t*-test, for statistical details see **Methods**.

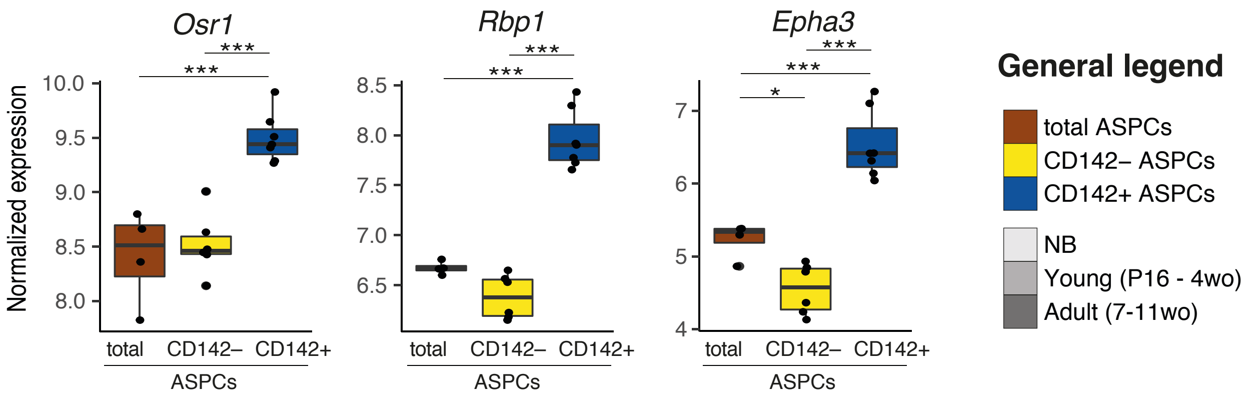


**Supplementary Figure 18. Expression profiles of RA-related CD142+ ASPC (Areg) markers**

Bulk RNA-seq-derived expression plots of Areg-specific genes involved in retinoic acid (RA) signalling: *Rbp1* (Retinol-binding protein 1), *Osr1* (Odd-skipped related transcription factor 1) and *Epha3* (Eph receptor A3);

*P ≤ 0.05, **P ≤ 0.01, ***P ≤ 0.001, one-way ANOVA and Tukey HSD *post hoc* test, for statistical details see **Methods**.

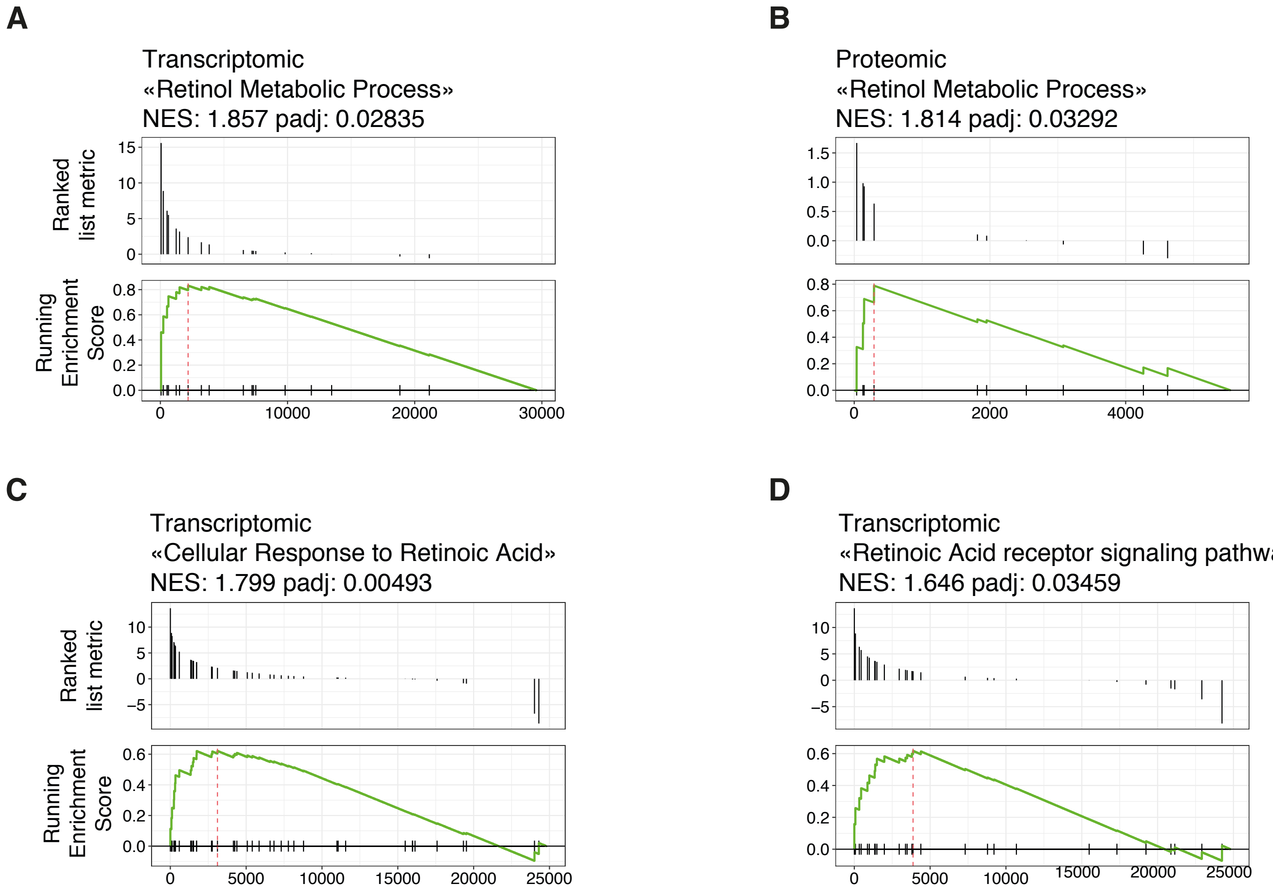


**Supplementary Figure 19. Retinoic acid-related terms are enriched in CD142+ ASPCs (Aregs) compared to their negative counterparts**

- **(A)** The term "retinol metabolic process" (GO:0042572) was identified as significantly enriched in freshly isolated CD142+ *versus* CD142– ASPCs by GSEA when performed on differential expression data based on bulk RNA-seq of freshly sorted ASPCs;
- **(B)** The term "retinol metabolic process" (GO:0042572) was identified as significantly enriched in freshly isolated CD142+ *versus* CD142– ASPCs when performed on differential expression data based on mass spectrometry of freshly sorted ASPCs;
- **(C)** The term "cellular response to retinoic acid (RA)" (GO:0071300) was identified as significantly enriched in CD142+ *versus* CD142– ASPCs after adipogenic differentiation when performed on differential expression data based on bulk RNA-seq of ASPCs after adipogenic differentiation;
- **(D)** The term "RA receptor signalling pathway" (GO:0048384) was identified as significantly enriched in CD142+ *versus* CD142– ASPCs after adipogenic differentiation when performed on differential expression data based on bulk RNA-seq of ASPCs after adipogenic differentiation.

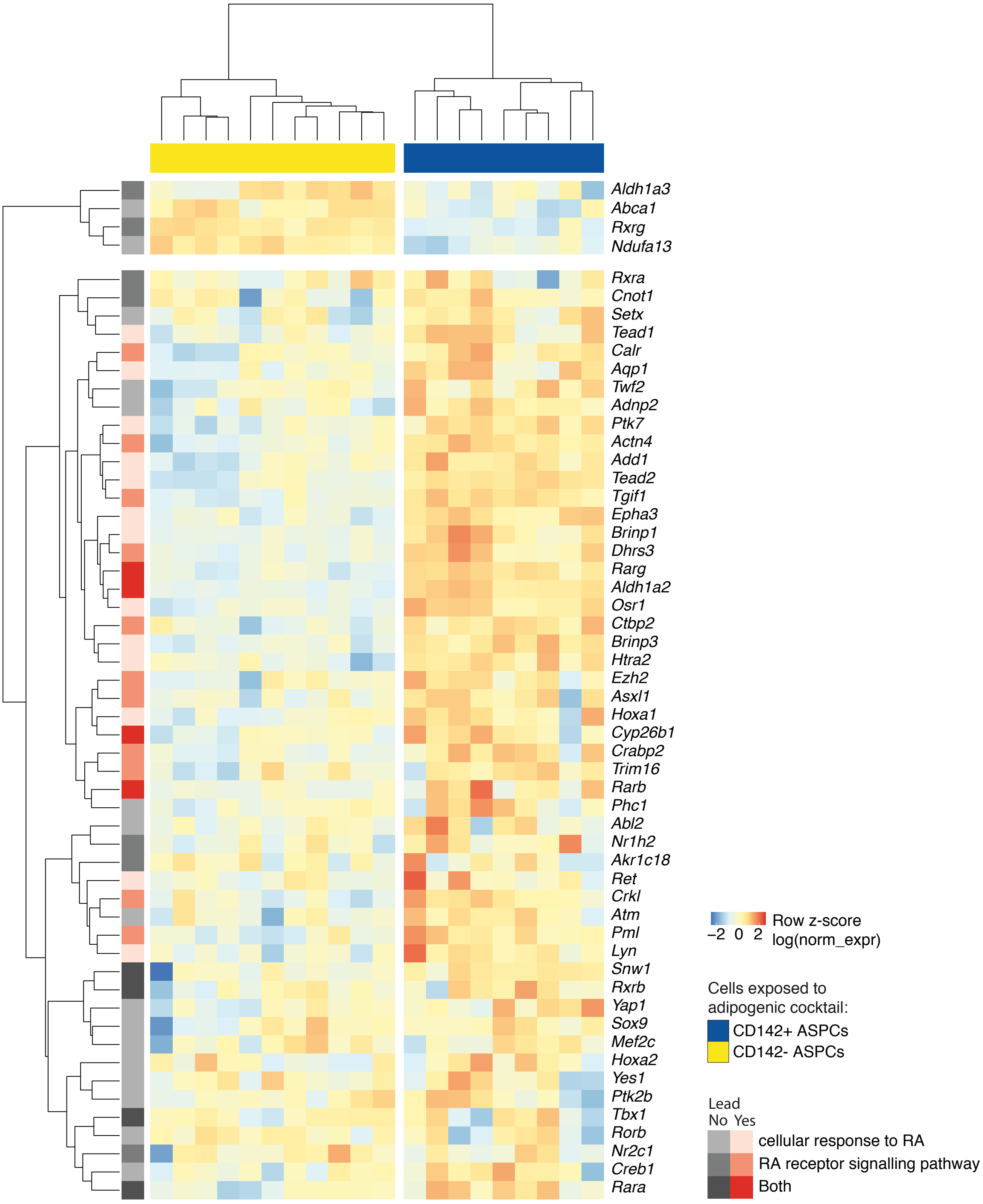


**Supplementary Figure 20. Expression of genes linked to RA-associated terms in CD142– and CD142+ ASPCs (Aregs) post-differentiation**

Expression heatmap of genes linked to “Cellular response to RA” (GO:0071300) and “RA receptor signalling pathway” (GO:0048384) across bulk RNA-seq samples of CD142– and CD142+ ASPCs (Aregs) post-differentiation; genes identified as lead by GSEA are indicated in red (**Methods**); log normalized expression scaled by row.

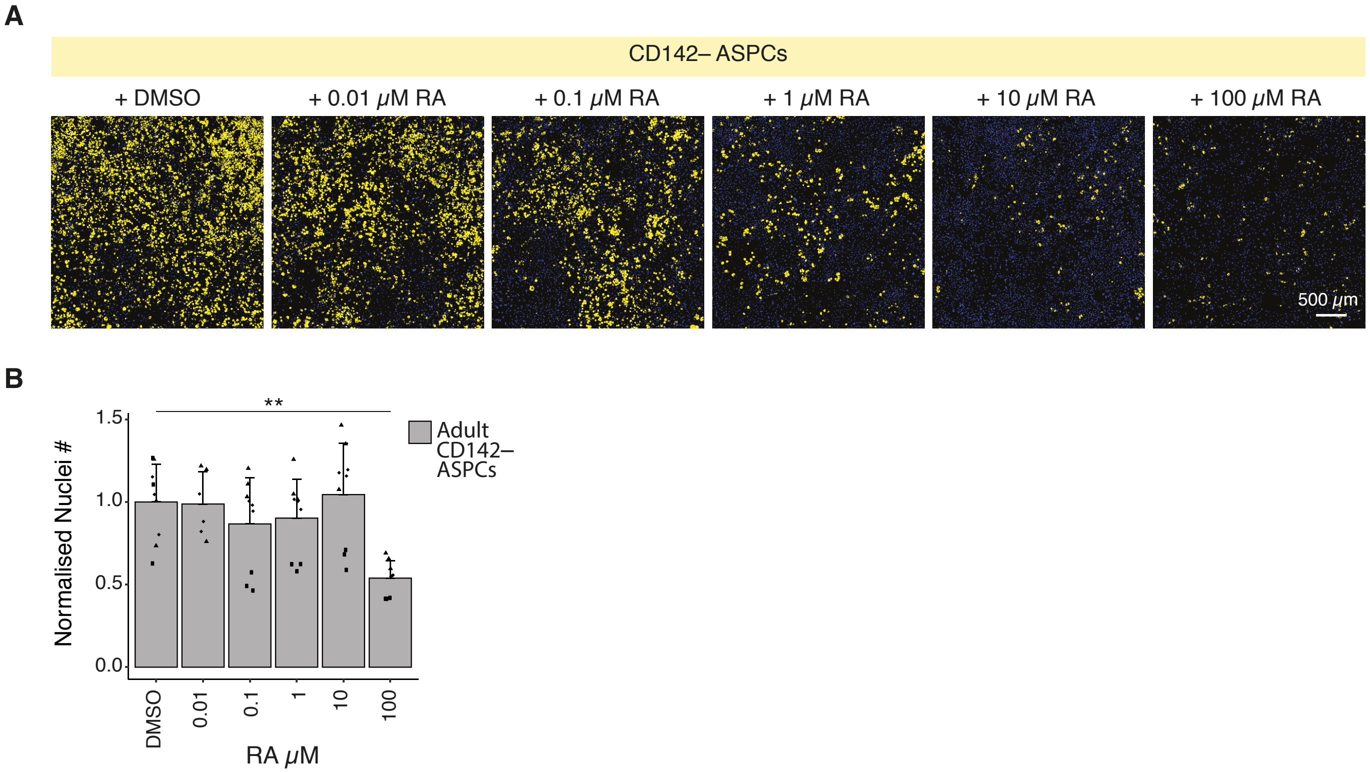


**Supplementary Figure 21. Retinoic acid treatment inhibits CD142– ASPC differentiation into adipocytes**

**(A)** Representative fluorescence microscopy images of CD142– ASPCs after *in vitro* adipogenic differentiation with the differentiation cocktail supplemented with DMSO (RA carrier) and RA at the indicated concentrations; the presented images correspond to 25 tiled and thresholded 20x images containing 7-8 z-stacks in order to capture most of the well surface (**Methods**);

**(B)** Bar plots showing the nuclei numbers for the cellular fractions shown in **A**; marker shapes correspond to different biological replicates, n=9, 3 biological replicates, 3 independent wells for each;

In all images, nuclei are stained with Hoechst (blue) and lipids are stained with Bodipy (yellow); scale bars, 500 μm; *P ≤ 0.05, **P ≤ 0.01, ***P ≤ 0.001, pairwise two-sided *t*-test, for statistical details see **Methods**.

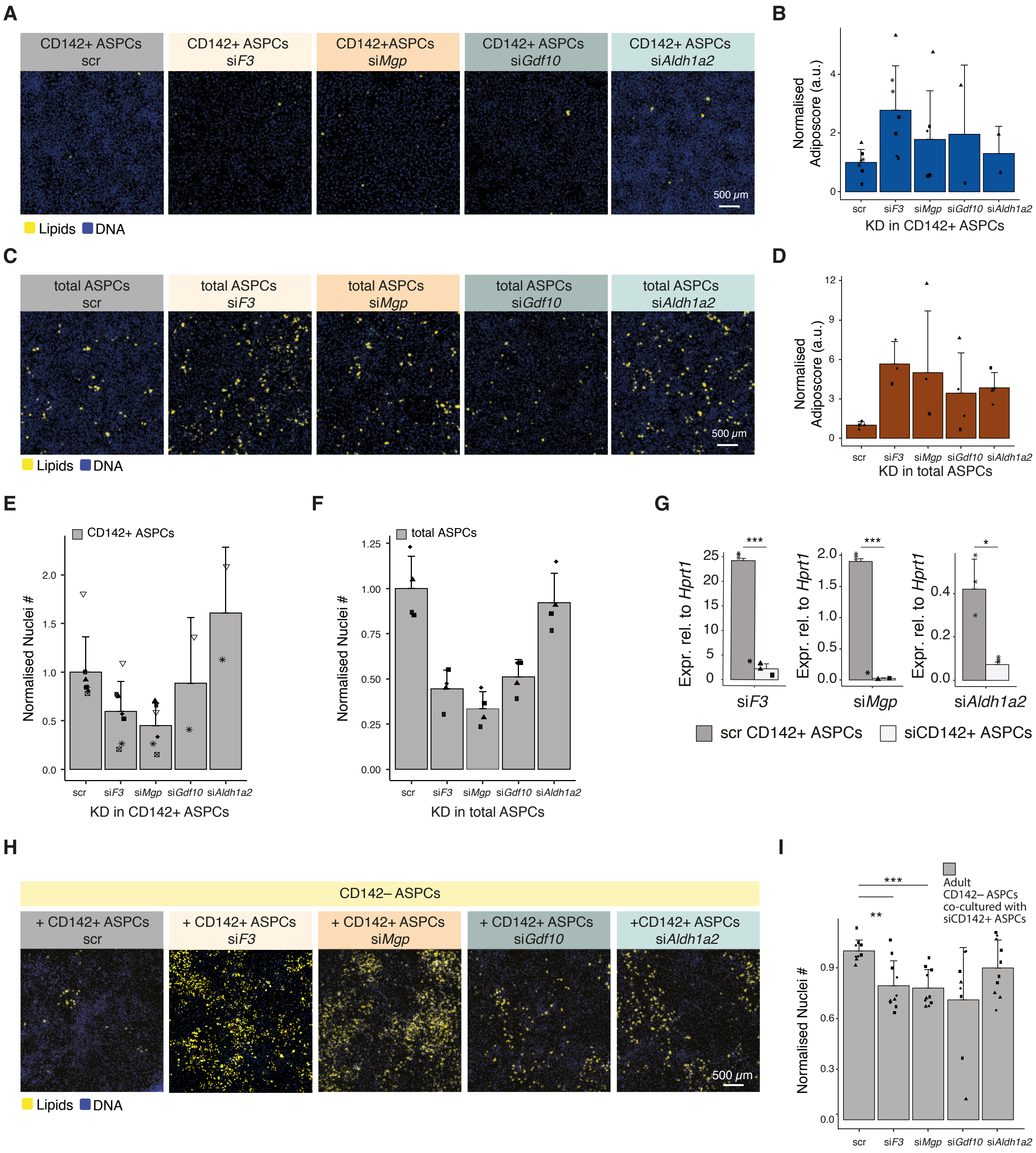


**Supplementary Figure 22** | See next page for caption

**Supplementary Figure 22. siRNA mediated knockdown of CD142+ ASPC (Areg) candidates**

- **(A)** Representative fluorescence microscopy images of CD142+ ASPCs (Aregs) after *in vitro* adipogenic differentiation with control (scr) or siRNA-mediated knockdowns of selected CD142+ ASPC (Areg)-specific candidate genes: *F3*, *Mgp*, *Gdf10* and *Aldh1a2*;
- **(B)** Fraction of differentiated CD142+ ASPCs (Aregs), as quantified by the “adiposcore”, for each siRNA knockdown assay shown in **A**; marker shapes correspond to different biological replicates, n=2-7, 2-5 biological replicates;
- **(C)** Representative fluorescence microscopy images of total ASPCs after in *vitro* adipogenic differentiation with control (scr) or siRNA-mediated knockdowns of selected CD142+ ASPC (Areg)-specific candidate genes: *F3*, *Mgp*, *Gdf10* and *Aldh1a2*;
- **(D)** Fraction of differentiated total ASPCs, as quantified by the “adiposcore”, for each siRNA knockdown assay shown in **C**; marker shapes correspond to different biological replicates, n=4, 3 biological replicates;
- **(E)** Bar plot showing the nuclei numbers for the cellular fractions shown in **A**; marker shapes correspond to different biological replicates, n=2-7, 2-5 biological replicates;
- **(F)** Bar plot showing the nuclei numbers for the cellular fractions shown in **C**; marker shapes correspond to different biological replicates, n=4, 3 biological replicates;
- **(G)** qPCR-measured gene expression of *F3*, *Mgp* and *Aldh1a2* in CD142+ ASPCs (Aregs) carrying control (scr) or siRNA-mediated knockdown of the indicated gene, 1-2 biological replicates, 3 independent wells for each;
- **(H)** Representative fluorescence microscopy images of CD142– ASPCs, co-cultured with CD142+ ASPCs (Aregs) carrying control (scr) or siRNA-mediated knockdowns of selected CD142+ ASPCs (Areg)-specific candidate genes: *F3*, *Mgp*, *Gdf10* and *Aldh1a2* after adipogenic differentiation; the presented images correspond to 25 tiled and thresholded 20x images containing 7-8 z-stacks in order to capture most of the well surface (**Methods**);
- **(I)** Bar plot showing the nuclei numbers for the cellular fractions shown in **H**; marker shapes correspond to different biological replicates, n=10, 4 biological replicates;

In all images, nuclei are stained with Hoechst (blue) and lipids are stained with Bodipy (yellow); scale bars, 500μm, bar colours: total ASPCs — brown, CD142+ ASPCs — blue, nuclei (**E**-**I**) - grey; *P ≤ 0.05, **P ≤ 0.01, ***P ≤ 0.001, pairwise two-sided *t*-test, for statistical details see **Methods**.

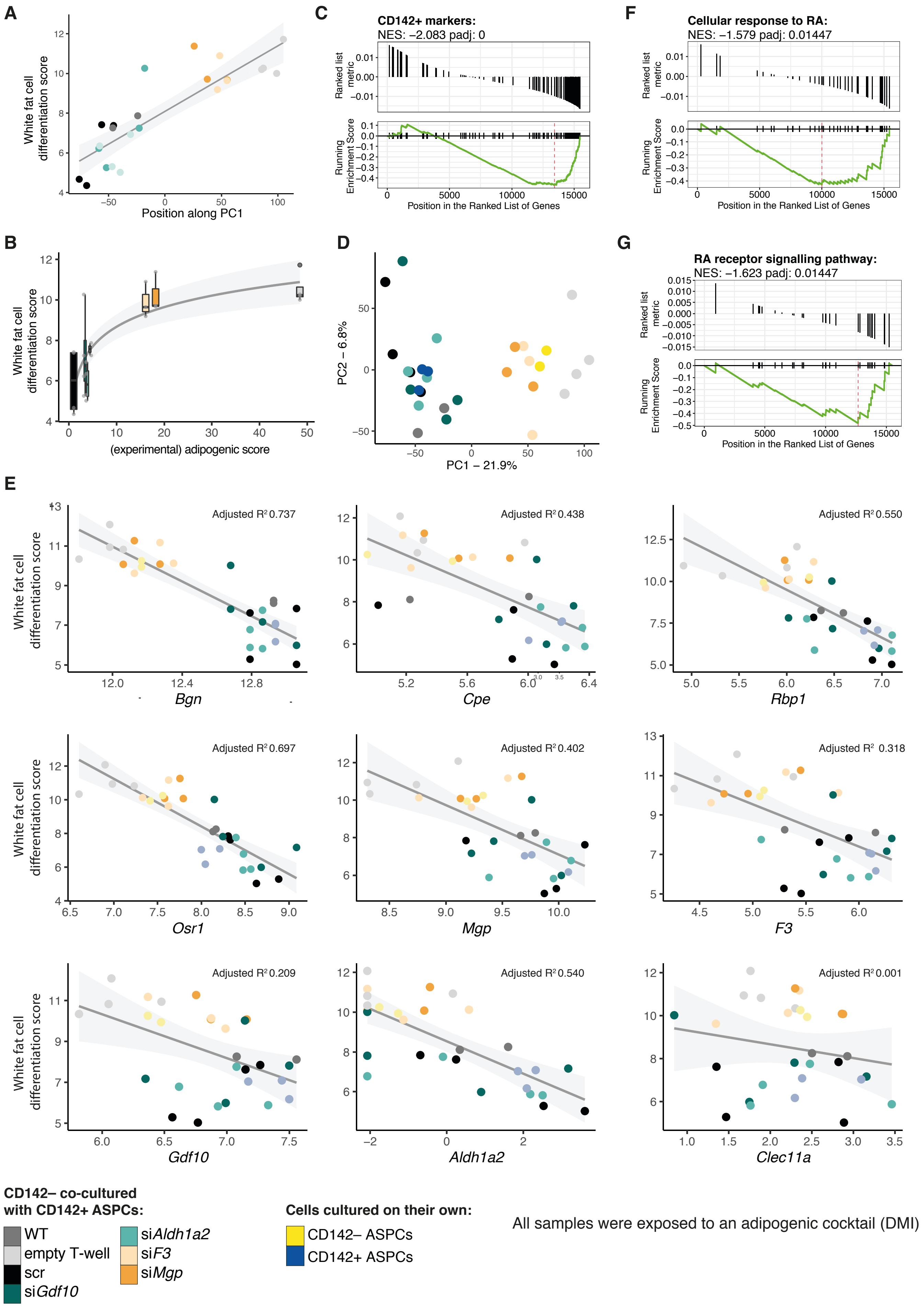


**Supplementary Figure 23** | See next page for caption

**Supplementary Figure 23. CD142– ASPCs exposed to the secretome of "active Aregs" (CD142+ ASPCs) exhibit upregulation of CD142+ ASPC (Areg) markers and retinoic acid-related genes**

- **(A)** Correlation of the position on the PC1 of the PCA shown in **Fig. 4C** and the "white fat cell differentiation score" (GO:0050872) (**Methods**); the colour legend is listed at the bottom of the figure;
- **(B)** Correlation of the experimental adipogenic score *versus* the "white fat cell differentiation score" (GO:0050872) (**Methods**);
- **(C)** GSEA results of the top CD142+ ASPC (Aregs) markers (**Suppl. Table 1**) performed on the genes driving PC1 of the PCA shown in **Fig. 4C**;
- **(D)** PCA of the bulk RNA-seq of CD142– ASPCs that are exposed (*via* transwell (T-well)) to distinct types of CD142+ ASPCs (knockdown and control), together with CD142+ ASPCs (Aregs) and CD142– ASPCs post-differentiation samples;
- **(E)** Correlation of the normalized gene expression of specific CD142+ ASPC (Areg) candidates *versus* the "white fat cell differentiation score";
- **(F)** GSEA results of "cellular response to retinoic acid (RA)" (GO:0071300) performed on the genes driving the PC1 of the PCA shown in **Fig. 4C**;
- **(G)** GSEA results of "RA receptor signalling pathway" (GO:0048384) performed on the genes driving PC1 of the PCA shown in **Fig. 4C**.

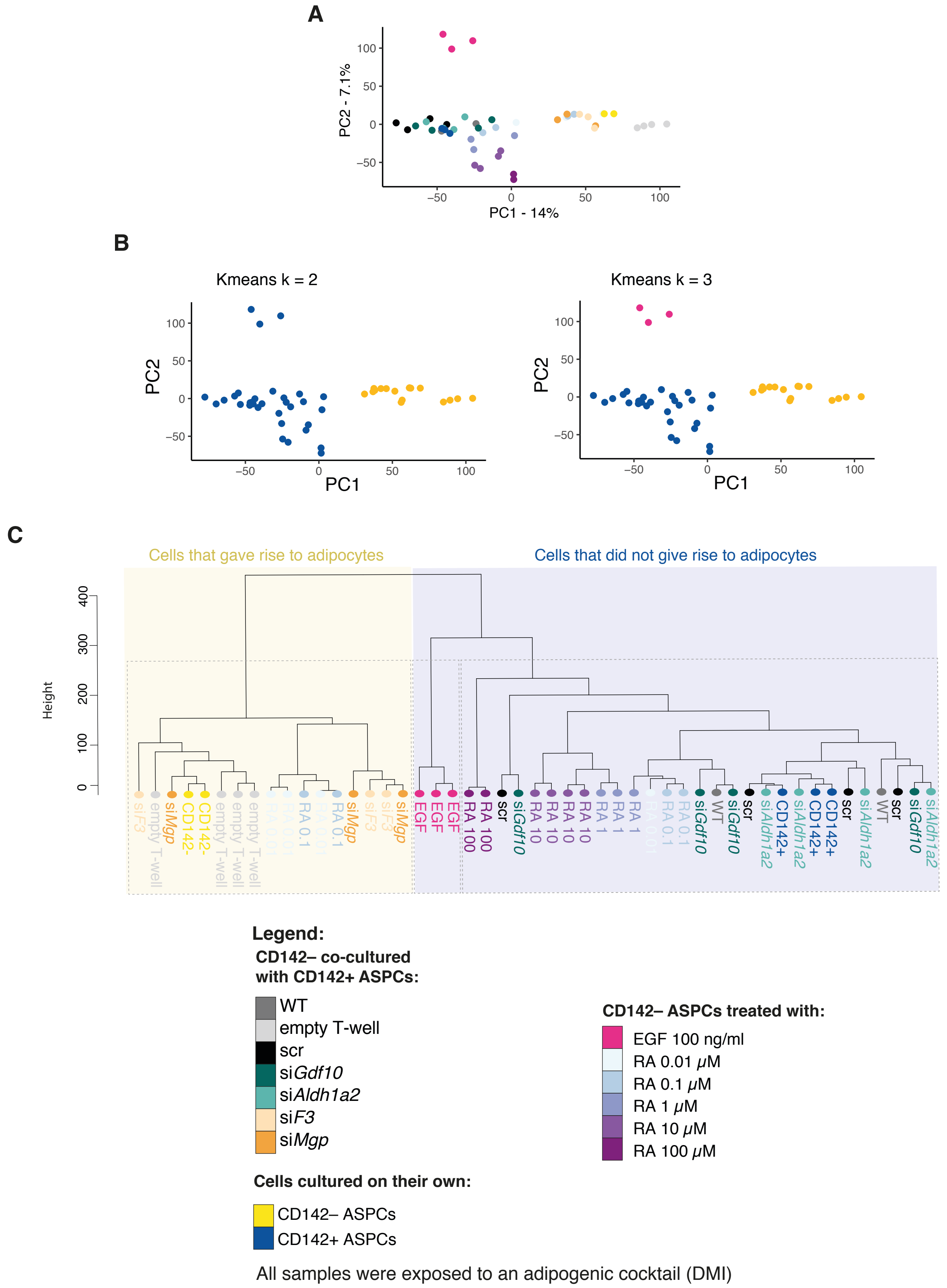


**Supplementary Figure 24. CD142– ASPCs co-cultured with "active Aregs" CD142+ ASPCs are transcriptionally similar to CD142– ASPCs treated with RA**

- **(A)** PCA of the bulk RNA-seq of CD142– ASPCs that were exposed (*via* transwell (T-well)) to the secretome of distinct CD142+ ASPC (Areg) types (knockdown or control) as listed in the colour legend, together with CD142+ (Aregs) and CD142– ASPCs having been exposed to differentiation medium on their own as well as CD142– ASPCs treated with EGF or different concentrations of RA;
- **(B)** k-means clustering results performed on the first five principal components of the PCA shown in **A**, for k = 2 (**left**) or k = 3 (**right**);
- **(C)** Hierarchical clustering tree based on bulk RNA-seq samples described in **A**.

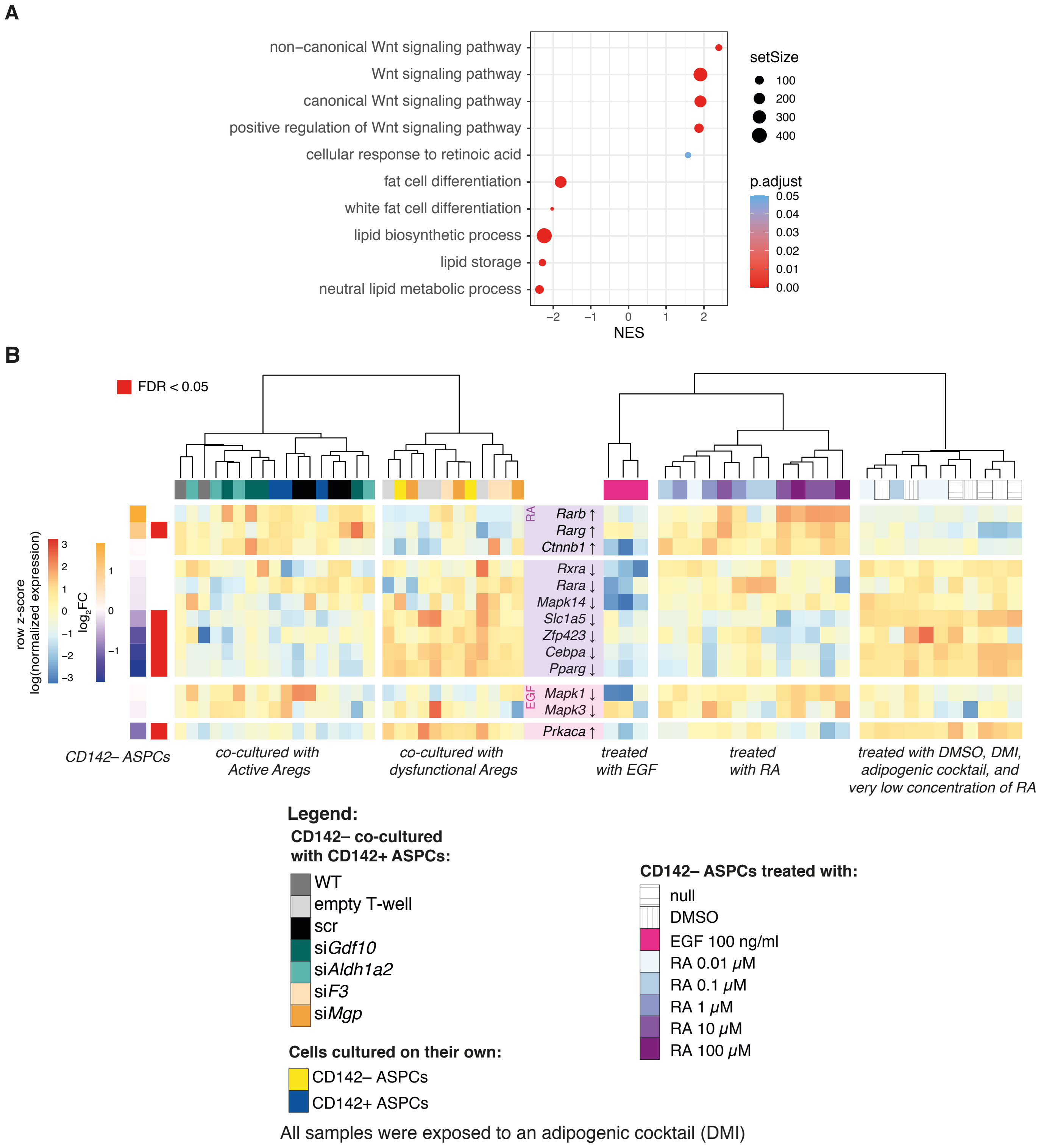


**Supplementary Figure 25. CD142– ASPCs exposed to the secretome of "active Aregs" (CD142+ ASPCs) exhibit upregulation of genes known to be involved in RA-mediated inhibition of adipogenesis**

**(A)** GSEA results of selected significant terms performed on the genes driving PC1 of the PCA shown in **Fig. 4C**;

**(B)** Expression heatmap of genes identified as involved in the RA- or EGF-mediated inhibition of adipogenesis, based on manual curation of the literature. The arrow next to each gene indicates the reported expression response to either RA or EGF. The heatmap on the **left** shows the log normalized expression by row across bulk RNA-seq samples of CD142– ASPCs co-cultured (*via* transwell (T-well)) with "active Aregs" (WT, scr, si*Aldh1a2*, si*Gdf10*) and standard CD142+ ASPCs (Aregs) post-differentiation (left cluster) and CD142– ASPCs co-cultured with "dysfunctional Aregs" (si*F3*, si*Mgp*) and CD142– ASPCs post-differentiation (right cluster); the heatmap on the **right** shows the normalized expression by row across bulk RNA-seq samples of CD142– ASPCs treated with EGF (left cluster), with RA (middle cluster) or with DMI with or without DMSO (RA carrier) (right cluster); for the two gene categories (involved in RA (purple) or EGF (pink)), the genes are ordered from top to bottom by the $\log_2$FC of CD142– ASPCs co-cultured with "active Aregs" over CD142– ASPCs co-cultured with "dysfunctional Aregs"; significantly differentially expressed genes (FDR < 0.05) are indicated by the red vertical bar.

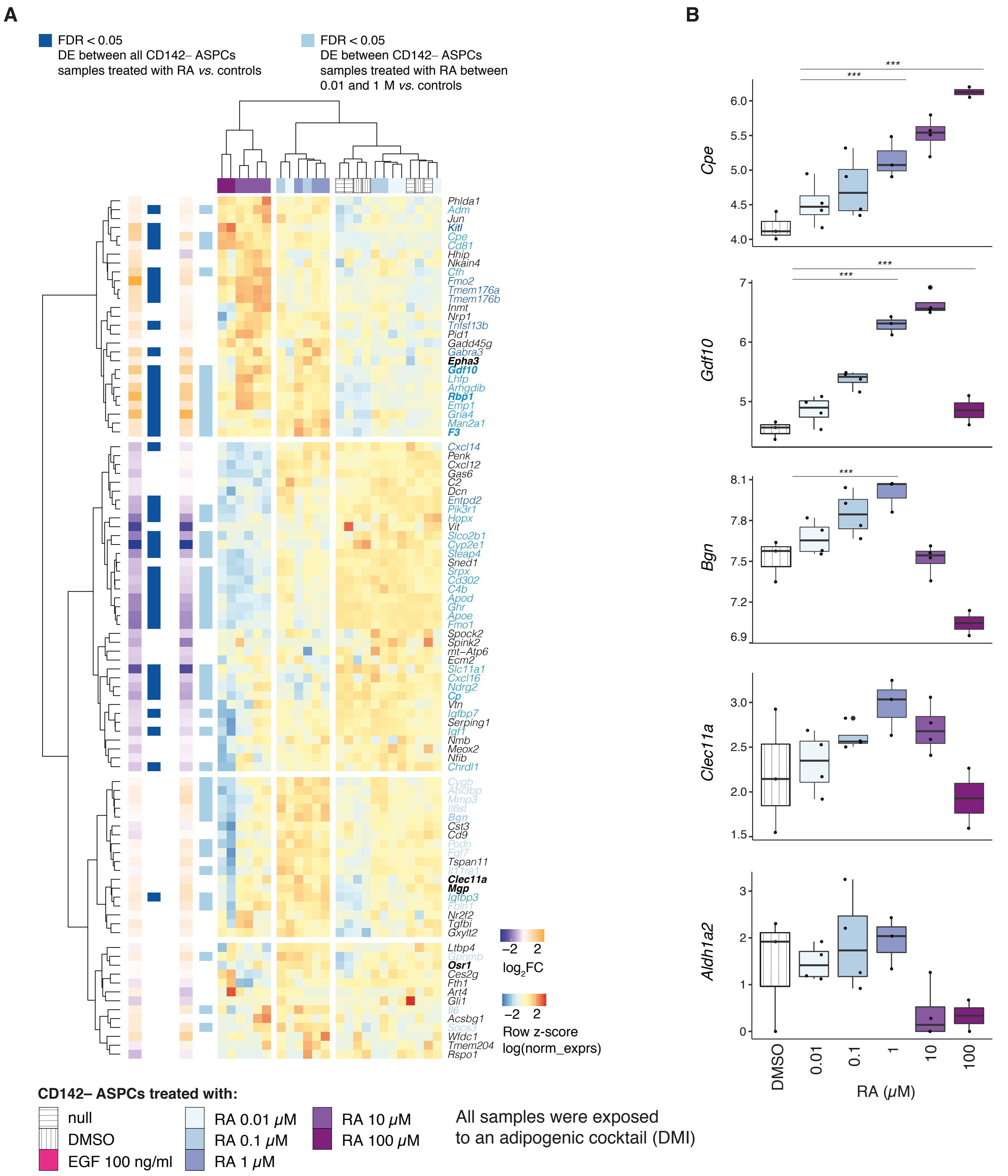


**Supplementary Figure 26. Key CD142+ ASPC (Areg) markers are upregulated in CD142– ASPCs treated with RA**

**(A)** Expression heatmap of the top CD142+ ASPC (Areg) markers (**Suppl. Table 1**) across bulk RNA-seq of CD142– ASPCs exposed to adipogenic cocktail (null) together with DMSO (RA carrier), EGF or different RA concentrations. The $\log_2$FC and significance are based on the differential expression analysis between (**left**, dark blue) CD142– ASPCs treated with all RA concentrations or (**right**, light blue) treated with RA concentrations ranging between 0.01 and 1 $\mu$M *versus* controls (treatment with DMI with or without DMSO);

**(B)** Boxplots showing the distribution of the log normalized expression (y axis) of *Cpe*, *Gdf10*, *Bgn*, *Clec11a* and *Aldh1a2* across transcriptomic data from CD142– ASPCs treated with DMSO or different RA concentrations (0.01 to 100 $\mu$M); the indicated significance is based on the differential analysis result between all RA treated samples or samples treated with RA at a concentration between 0.01 and 1 $\mu$M *versus* controls (treatment with DMI with or without DMSO).

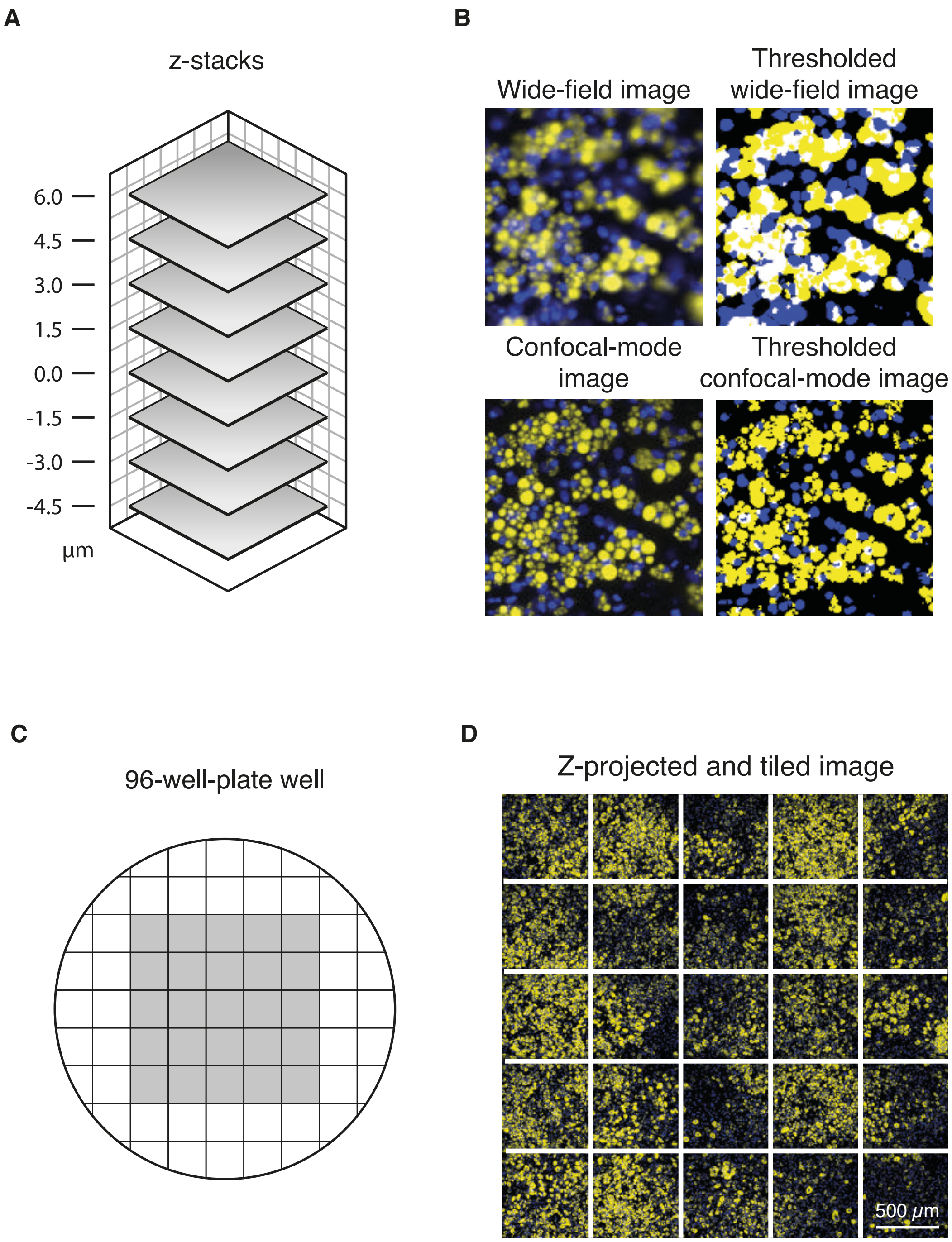


**Supplementary Figure 27. Imaging *in vitro* adipogenesis.**

- **(A)** Schematic representation of z-stacks acquired with Perkin Elmer Operetta platform during imaging wells that contain terminally differentiated *in vitro* adipocytes;
- **(B)** Representative images resulting from wide-field imaging (top) and confocal-mode imaging (bottom) and the respective thresholding result;
- **(C)** Schematic representation of the 25 images acquired within a 96-well-plate well;
- **(D)** A representative image resulting from confocal-mode imaging after tiling 25 acquired and z-projected images (the white grid has been added graphically for indicative purposes).

***Supplementary Table 1:*** Top 100 genes specific for CD142+ ASPCs (Aregs) issued from an integration of scRNA-seq (publicly available) and bulk RNA-seq (publicly available and generated in the presented study) datasets.

| **Aregs 100 Top Markers (1/3)** | | | |
|---|---|---|---|
| **gene ID** | **Single-cell RNA-seq avg logFC** | **Bulk RNA-seq $log_2FC$** | **Mass spectrometry $log_2FC$** |
| ***Gas6*** | 1.734934037 | 2.648 | 0.573588699 |
| ***Cxcl12*** | 1.695784366 | 1.306 | NA |
| ***Inmt*** | 1.69273622 | 2.106 | 2.232207245 |
| ***Apoe*** | 1.617141956 | NA | 0.523858909 |
| ***Gdf10*** | 1.485475157 | 2.51 | 1.002882979 |
| ***Clec11a*** | 1.339330278 | 3.383 | NA |
| ***F3*** | 1.306446139 | 4.364 | 3.852178637 |
| ***Mgp*** | 1.211678039 | 4.412 | 1.518067937 |
| ***C2*** | 1.197130218 | 2.295 | 0.870029683 |
| ***Fmo2*** | 1.16670468 | 4.944 | 3.451288912 |
| ***Igfbp7*** | 1.111835725 | NA | 0.929751786 |
| ***Igfbp3*** | 1.080026504 | NA | NA |
| ***Vit*** | 1.016376833 | 3.256 | NA |
| ***Bgn*** | 0.991099847 | 1.424 | 2.471041469 |
| ***Steap4*** | 0.986622284 | 1.661 | 3.100726969 |
| ***Gadd45g*** | 0.980075579 | NA | NA |
| ***Tmem176b*** | 0.972756487 | NA | NA |
| ***Igf1*** | 0.96280102 | NA | NA |
| ***Cygb*** | 0.931309354 | 1.784 | 2.216748912 |
| ***Tmem176a*** | 0.908769033 | NA | NA |
| ***Chrdl1*** | 0.892908836 | 2.42 | NA |
| ***Srpx*** | 0.890644606 | NA | NA |
| ***Nmb*** | 0.883789886 | 1.416 | NA |
| ***Abi3bp*** | 0.848341847 | 1.031 | -0.059743746 |
| ***Penk*** | 0.826923908 | NA | NA |
| ***Cp*** | 0.825599904 | NA | 2.109079548 |
| ***Nrp1*** | 0.8204081 | 1.223 | 0.247841773 |
| ***Vtn*** | 0.795788451 | NA | NA |
| ***Cxcl14*** | 0.783008446 | NA | 0.62827134 |
| ***Cst3*** | 0.77369243 | 1.874 | 1.244114617 |
| ***Mmp3*** | 0.738574153 | NA | NA |
| ***Il11ra1*** | 0.738383419 | NA | NA |
| ***Cd9*** | 0.735687305 | 1.422 | NA |
| ***C4b*** | 0.731043887 | -1.175 | -0.331515068 |
| ***Tgfbi*** | 0.729048091 | 2.618 | NA |

| Aregs 100 Top Markers (2/3) | | | |
|---|---|---|---|
| **gene ID** | **Single-cell RNA-seq avg logFC** | **Bulk RNA-seq $log_2FC$** | **Mass spectrometry $log_2FC$** |
| ***Cfh*** | 0.722718972 | NA | 0.547533656 |
| ***Socs3*** | 0.709242264 | NA | NA |
| ***Meox2*** | 0.678224608 | 3.734 | 1.593829126 |
| ***Il6*** | 0.66412741 | 1.652 | NA |
| ***Sepp1*** | 0.640493416 | 1.739 | NA |
| ***Cpe*** | 0.625167315 | 2.179 | NA |
| ***Osr1*** | 0.612115059 | 1.456 | 0.199888736 |
| ***Rbp1*** | 0.610362899 | 1.805 | 1.669933451 |
| ***Gpnmb*** | 0.602833659 | 1.878 | 2.115085125 |
| ***Nfib*** | 0.57788856 | NA | 0.844491716 |
| ***Apod*** | 0.575275481 | NA | 0.685823996 |
| ***Pid1*** | 0.563071453 | 2.35 | 0.313006138 |
| ***Phlda1*** | 0.562879145 | NA | 0.036171197 |
| ***Serping1*** | 0.560522695 | NA | 0.341085822 |
| ***Sned1*** | 0.554291693 | NA | NA |
| ***Fbln1*** | 0.533888358 | 1.53 | NA |
| ***Ltbp4*** | 0.518973443 | 1.06 | NA |
| ***Ndrg2*** | 0.514044932 | 3.057 | 1.805160555 |
| ***Ecm2*** | 0.484665409 | NA | NA |
| ***Entpd2*** | 0.483039679 | 1.163 | 2.834169075 |
| ***Podn*** | 0.482052268 | 1.259 | NA |
| ***Lhfp*** | 0.478587415 | NA | NA |
| ***Nr2f2*** | 0.474489959 | 1.723 | 0.88969045 |
| ***Dcn*** | 0.474343194 | NA | 0.410972407 |
| ***Fth1*** | 0.473941488 | NA | 0.444517095 |
| ***Kitl*** | 0.440142536 | 1.596 | NA |
| ***Ghr*** | 0.432869152 | 1.169 | NA |
| ***Cd302*** | 0.428252464 | NA | NA |
| ***Ces2g*** | 0.424280683 | NA | NA |
| ***Slco2b1*** | 0.410231034 | 1.509 | 0.823355932 |
| ***AI607873*** | 0.400312532 | NA | NA |
| ***Arhgdib*** | 0.393423297 | 4.014 | 2.192112505 |
| ***Fgf7*** | 0.384580273 | 1.086 | NA |
| ***Rspo1*** | 0.383282553 | NA | NA |
| ***Il6st*** | 0.383014607 | 1.062 | 0.77442495 |
| ***Gxylt2*** | 0.371476281 | NA | NA |
| ***mt-Atp6*** | 0.36908521 | NA | NA |
| ***Cd81*** | 0.367270304 | NA | 0.466254044 |
| ***Jun*** | 0.365838934 | NA | -0.309885662 |

| Aregs 100 Top Markers (3/3) | | | |
|---|---|---|---|
| **gene ID** | **Single-cell RNA-seq avg logFC** | **Bulk RNA-seq $log_2FC$** | **Mass spectrometry $log_2FC$** |
| ***Tspan11*** | 0.35906473 | 4.391 | NA |
| ***Emp1*** | 0.357989155 | NA | NA |
| ***Man2a1*** | 0.343316508 | NA | -0.163599593 |
| ***Adm*** | 0.339422608 | NA | NA |
| ***Cxcl16*** | 0.324837373 | NA | NA |
| ***Epha3*** | 0.312033992 | 2.171 | 1.239960604 |
| ***Fmo1*** | 0.304309182 | 1.679 | 1.319270289 |
| ***Tnfsf13b*** | 0.301936673 | 2.169 | NA |
| ***Gabra3*** | 0.299116342 | 1.848 | 0.488352387 |
| ***Spock2*** | 0.289710678 | 2.589 | NA |
| ***Pik3r1*** | 0.265155293 | NA | 0.003332054 |
| ***Acsbg1*** | NA | 6.015 | 0.926057088 |
| ***Spink2*** | NA | 5.451 | NA |
| ***Art4*** | NA | 4.883 | NA |
| ***Hopx*** | NA | 4.794 | 2.102595259 |
| ***Tmem204*** | NA | 4.763 | NA |
| ***Nkain4*** | NA | 4.643 | NA |
| ***Cldn22*** | NA | 4.32 | NA |
| ***Wfdc1*** | NA | 4.261 | NA |
| ***Hhip*** | NA | 4.214 | NA |
| ***Gria4*** | NA | 4.208 | NA |
| ***Cyp2e1*** | NA | 4.023 | NA |
| ***Slc11a1*** | NA | 4.015 | NA |
| ***Iyd*** | NA | 3.983 | NA |
| ***Gli1*** | NA | 3.964 | NA |
| ***Cav3*** | NA | 3.921 | NA |

***Supplementary Table 2:*** Summary of bulk RNA-seq and proteomic experiments conducted in the presented study.

| **TRANSCRIPTOMICS** | | | | | | |
|---|---|---|---|---|---|---|
| **Name** | **Fresh ASPCs** | **Expanded ASPCs** | **Post-diff ASPCs** | **RA & EGF treatment** | **KD assay** | **Age Assay** |
| **Design** | Sequenced were freshly purified murine ASPC cells isolated by FACS using four different anti-CD142 antibodies (Methods) | Sequenced were FACS-isolated murine CD142+ and CD142− ASPCs plated and expanded in culture for 4-5 days | Sequenced were FACS-isolated (with four different anti-CD142 antibodies) murine CD142+ and CD142− ASPCs plated, cultured to confluence and differentiated into adipocytes | Sequenced were CD142− ASPCs differentiated into adipocytes with retinoic acid (RA) or recombinant EGF treatment | Sequenced were CD142− APSCs differentiated into adipocytes in the presence of wild-type CD142+ ASPCs or CD142+ ASPCs with indicated KDs or an empty well | Sequenced were total, CD142− and CD142+ ASPCs isolated from P0, P16, 4wo, 7wo and 11wo mice |
| **Samples** | CD142+<br>CD142−<br>SCA-1+ | CD142+<br>CD142− | CD142+<br>CD142−<br>SCA-1+ | CD142− ASPCs treated with:<br>-0.01<br>-0.1<br>-1<br>-10<br>-100 µM of RA<br>-100 ng/µL of EGF | CD142− ASPCs co-cultured with CD142+ ASPCs:<br>-scr<br>-si*Mgp*<br>-siAldh1a2<br>-si*Gdf10*<br>-si*F3*<br>-WT<br>- with an empty well | CD142+<br>CD142−<br>SCA-1+ |

| **PROTEOMICS** | | |
|---|---|---|
| **Name** | **Fresh ASPCs** | **Expanded ASPCs** |
| **samples** | CD142+<br>CD142− | CD142+<br>CD142− |

***Supplementary Table 3:*** List of all antibodies employed in the experiments conducted in the presented study.

| Primary antibodies | | | | | | | | | | | |
|---|---|---|---|---|---|---|---|---|---|---|---|
| **Antibody** | **Clone** | **Clonality** | **Host** | **Reactivity** | **Conjugate** | **Application** | **Company** | **Ref#** | **Tested applications** | **Publication DOI** | **Dilution** |
| **anti-CD31** | MEC13.3 | monoclonal | Rat | Mouse | AF488 | FACS | BioLegend | 102514 | FC | 10.1038/ncomms 11302 | 1:500 |
| **anti-CD45** | 30-F11 | monoclonal | Rat | Mouse | AF488 | FACS | BioLegend | 103122 | FC | 10.1038/ncomms 11533 | 1:500 |
| **anti-CD142** | 001 | monoclonal | Rabbit | Mouse | PE | FACS | SinoBiological | 50413-R001-P | FC | 10.1038/s41586-018-0226-8 | 1:50 |
| **anti-CD142** | 001 | monoclonal | Rabbit | Mouse | Un-conjugated | FACS | SinoBiological | 50413-R001 | FC | 10.1126/sciadv.aa x9605 | 1:50 |
| **anti-CD142** | HTF-1 | monoclonal | Mouse | Mouse | PE | FACS | BiOrbyt | ORB507 485 | FC | N/A | 1:20 |
| **anti-CD142** | - | polyclonal | Goat | Mouse | PE | FACS | R&D Systems | FAB317 8P | FC | 10.3324/haematol .2019.220210 | 1:20 |
| **anti-SCA-1** | E13-161.7 | monoclonal | Rat | Mouse | PE-Cy7 | FACS | BioLegend | 122514 | FC | 10.1371/journal.p bio.1002562 | 1:500 |
| **anti-TER119** | TER-119 | monoclonal | Rat | Mouse | AF488 | FACS | BioLegend | 116215 | FC | 10.1016/j.bbalip.2 016.06.010 | 1:500 |

***Supplementary Table 4:*** Sequences of siRNA probes employed for Areg-related knock down experiments conducted in the presented study.

| siRNA name | Sequence |
|---|---|
| **Scrambled** | 5' /5Phos/rCrUrUrCrCrUrCrUrCrUrUrUrCrUrCrUrCrCrCrUrUrGrUGA 3' +<br>5' rUrCrArCrArArGrGrGrArGrArGrArArArGrArGrArGrGrArArGrGrA 3' - |
| **mm.Ri.F3.13.1** | 5' rGrGrArUrArUrCrArArCrUrGrArUrUrUrCrArArGrArCrAAT 3' +<br>5' rArUrUrGrUrCrUrUrGrArArArUrCrArGrUrUrGrArUrArUrCrCrArA 3' - |
| **mm.Ri.F3.13.2** | 5' rGrUrArArArArGrArCrUrCrArCrUrUrArCrArUrUrArGrUCA 3' +<br>5' rUrGrArCrUrArArUrGrUrArArGrUrGrArGrUrCrUrUrUrUrArCrArA 3' - |
| **mm.Ri.F3.13.3** | 5' rArUrGrUrUrArArArUrGrCrArGrGrArUrArUrUrUrCrUrGCT 3' +<br>5' rArGrCrArGrArArArUrArUrCrCrUrGrCrArUrUrUrArArCrArUrGrU 3' - |
| **mm.Ri.Mgp.13.1** | 5' rCrUrCrCrUrGrCrArGrUrArGrCrArUrUrArCrUrGrArArGTA 3' +<br>5' rUrArCrUrUrCrArGrUrArArUrGrCrUrArCrUrGrCrArGrGrArGrArU 3' - |
| **mm.Ri.Mgp.13.2** | 5' rCrCrUrArUrGrArArArUrCrArGrUrCrCrCrUrUrCrArUrCAA 3' +<br>5' rUrUrGrArUrGrArArGrGrGrArCrUrGrArUrUrUrCrArUrArGrGrArC 3' - |
| **mm.Ri.Mgp.13.3** | 5' rGrGrCrArArCrCrCrUrGrUrGrCrUrArCrGrArArUrCrUrCAC 3' +<br>5' rGrUrGrArGrArUrUrCrGrUrArGrCrArCrArGrGrGrUrUrGrCrCrArC 3' - |
| **mm.Ri.Gdf10.13.1** | 5' rGrArGrUrArUrUrUrUrCrUrCrUrUrArCrGrUrUrUrCrArCTA 3' +<br>5' rUrArGrUrGrArArArCrGrUrArArGrArGrArArArArUrArCrUrCrArC 3' - |
| **mm.Ri.Gdf10.13.2** | 5' rUrArArGrArUrGrGrCrUrUrCrArArGrArUrArGrArArGrACA 3' +<br>5' rUrGrUrCrUrUrCrUrArUrCrUrUrGrArArGrCrCrArUrCrUrUrArCrC 3' - |
| **mm.Ri.Gdf10.13.3** | 5' rCrGrGrGrUrGrGrArArUrGrArArUrGrGrArUrCrArUrCrUCT 3' +<br>5' rArGrArGrArUrGrArUrCrCrArUrUrCrArUrUrCrCrArCrCrCrGrArU 3' - |
| **mm.Ri.Aldh1a2.13.1** | 5' rGrArGrGrUrUrGrGrArArArGrCrUrUrArUrUrCrArArGrAAG 3' +<br>5' rCrUrUrCrUrUrGrArArUrArArGrCrUrUrUrCrCrArArCrCrUrCrArG 3' - |
| **mm.Ri.Aldh1a2.13.2** | 5' rCrArArGrArArArUrUrUrUrGrArGrGrUrUrUrArArGrArCTA 3' +<br>5' rUrArGrUrCrUrUrArArArCrCrUrCrArArArArUrUrUrCrUrUrGrArA 3' - |
| **mm.Ri.Aldh1a2.13.3** | 5' rGrGrUrGrArArUrArCrCrGrUrUrUrArUrUrUrArArArArATG 3' +<br>5' rCrArUrUrUrUrUrArArArUrArArArCrGrGrUrArUrUrCrArCrCrCrA 3' - |
